# Efficient *de novo* assembly of eleven human genomes using PromethION sequencing and a novel nanopore toolkit

**DOI:** 10.1101/715722

**Authors:** Kishwar Shafin, Trevor Pesout, Ryan Lorig-Roach, Marina Haukness, Hugh E. Olsen, Colleen Bosworth, Joel Armstrong, Kristof Tigyi, Nicholas Maurer, Sergey Koren, Fritz J. Sedlazeck, Tobias Marschall, Simon Mayes, Vania Costa, Justin M. Zook, Kelvin J. Liu, Duncan Kilburn, Melanie Sorensen, Katy M. Munson, Mitchell R. Vollger, Evan E. Eichler, Sofie Salama, David Haussler, Richard E. Green, Mark Akeson, Adam Phillippy, Karen H. Miga, Paolo Carnevali, Miten Jain, Benedict Paten

**Affiliations:** UC Santa Cruz Genomics Institute, Santa Cruz, CA 95064, USA; Chan Zuckerberg Initiative, Redwood City, CA 94063, USA; Oxford Nanopore Technologies, Oxford Science Park, OX4 4DQ, UK; Genome Informatics Section, Computational and Statistical Genomics Branch, National Human Genome Research Institute, Bethesda, MD 20892, USA; Baylor College of Medicine, Human Genome Sequencing Center, Houston, TX 77030, USA; Max Planck Institute for Informatics, 66123 Saarbrücken, Germany; Howard Hughes Medical Institute, University of California, Santa Cruz, CA 95064, USA; National Institute of Standards and Technology, Gaithersburg, MD 20899, USA; Circulomics Inc, Baltimore, MD 21202, USA; Department of Genome Sciences, University of Washington School of Medicine, Seattle, WA 98195, USA

**Keywords:** Nanopore, Assembly, Polishing, PromethION, Human Genomes, Shasta, MarginPolish, HELEN

## Abstract

Present workflows for producing human genome assemblies from long-read technologies have cost and production time bottlenecks that prohibit efficient scaling to large cohorts. We demonstrate an optimized PromethION nanopore sequencing method for eleven human genomes. The sequencing, performed on one machine in nine days, achieved an average 63x coverage, 42 Kb read N50, 90% median read identity and 6.5x coverage in 100 Kb+ reads using just three flow cells per sample. To assemble these data we introduce new computational tools: Shasta - a *de novo* long read assembler, and MarginPolish & HELEN - a suite of nanopore assembly polishing algorithms. On a single commercial compute node Shasta can produce a complete human genome assembly in under six hours, and MarginPolish & HELEN can polish the result in just over a day, achieving 99.9% identity (QV30) for haploid samples from nanopore reads alone. We evaluate assembly performance for diploid, haploid and trio-binned human samples in terms of accuracy, cost, and time and demonstrate improvements relative to current state-of-the-art methods in all areas. We further show that addition of proximity ligation (Hi-C) sequencing yields near chromosome-level scaffolds for all eleven genomes.

## Introduction

Short-read sequencing reference-assembly mapping methods only assay about 90% of the current reference human genome assembly [1], and closer to 80% at high-confidence [2]. The latest incarnations of these methods are highly accurate with respect to single nucleotide variants (SNVs) and short insertions and deletions (indels) within this mappable portion of the reference genome [3]. However, short reads are much less able to *de novo* assemble a new genome [4], to discover structural variations (SVs) [5, 6] (including large indels and base-level resolved copy number variations), and are generally unable to resolve phasing relationships without exploiting transmission information or haplotype panels [7].

Third generation sequencing technologies, including linked-reads [8, 9, 10] and long-read technologies [11, 12], get around the fundamental limitations of short-read sequencing for genome inference by providing more information per sequencing observation. In addition to increasingly being used within reference guided methods [1, 13, 14, 15], long-read technologies can generate highly contiguous *de novo* genome assemblies [16].

Nanopore sequencing, as commercialized by Oxford Nanopore Technologies (ONT), is particularly applicable to *de novo* genome assembly because it can produce high yields of very long 100+ kilobase (Kb) reads [17]. Very long reads hold the promise of facilitating contiguous, unbroken assembly of the most challenging regions of the human genome, including centromeric satellites, acrocentric short arms, rDNA arrays, and recent segmental duplications [18, 19, 20]. We contributed to the recent consortium-wide effort to perform the *de novo* assembly of a nanopore sequencing based human genome [17]. This earlier effort required considerable resources, including 53 ONT MinION flow cells and an assembly process that required over 150,000 CPU hours and weeks of wall-clock time, quantities that are unfeasible for production scale replication.

Making nanopore long-read *de novo* assembly easy, cheap and fast will enable new research. It will permit both more comprehensive and unbiased assessment of human variation, and creation of highly contiguous assemblies for a wide variety of plant and animal genomes. Here we report the *de novo* assembly of eleven diverse human genomes at near chromosome scale using a combination of nanopore and proximity-ligation (HiC) sequencing [8]. We demonstrate a substantial improvement in yields and read lengths for human genome sequencing at reduced time, labor, and cost relative to earlier efforts. Coupled to this, we introduce a toolkit for nanopore data assembly and polishing that is orders of magnitude faster than state-of-the-art methods.

## Results

### Nanopore sequencing eleven human genomes in nine days

We selected for sequencing eleven, low-passage (six passages), human cell lines of the offspring of parent-child trios from the 1000 Genomes Project (1KGP) [21] and Genome-in-a-Bottle (GIAB) [22] sample collections. The subset of 1KGP samples were selected to maximize allelic diversity and minimize passage (see Online Methods).

We performed PromethION nanopore sequencing and HiC Illumina sequencing for the eleven genomes. Briefly, we isolated HMW DNA from flash-frozen 50 million cell pellets using the QIAGEN Puregene kit, with some modifications to the standard protocol to ensure DNA integrity (see Online Methods). For nanopore sequencing, we performed a size selection to remove fragments <10 kilobases (Kb) using the Circulomics SRE kit, followed by library preparation using the ONT ligation kit (SQK-LSK109). We used three flow cells per genome, with each flow cell receiving a nuclease flush every 20-24 hours. This flush removed long DNA fragments that could cause the pores to become blocked over time. Each flow cell received a fresh library of the same sample after the nuclease flush. A total of two nuclease flushes were performed per flow cell, and each flow cell received a total of three sequencing libraries. We used Guppy version 2.3.5 with the high accuracy flipflop model for basecalling (see Online Methods).

The nanopore sequencing for these eleven genomes was performed in nine days, producing 2.3 terabases of sequence. This was made possible by running up to 15 flow cells in parallel during these sequencing runs. Results are shown in Fig. 1 and Supplementary Tables 1, 2, and 3. Nanopore sequencing yielded an average of 69 gigabases (Gb) per flow cell, with the total throughput per individual genome ranging between 48x (158 Gb) and 85x (280 Gb) coverage per genome (Fig. 1a). The read N50s for the sequencing runs ranged between 28 Kb and 51 Kb (Fig. 1b). We aligned nanopore reads to the human reference genome (GRCh38) and calculated their alignment identity to assess sequence quality (see Online Methods). We observed that the median and modal alignment identity was 90% and 93% respectively (Fig. 1c). The sequencing data per individual genome included an average of 55x coverage arising from 10 Kb+ reads, and 6.5x coverage from 100 Kb+ reads (Fig. 1d). This was in large part due to size-selection which yielded an enrichment of reads longer than 10 Kb.

**Figure 1:**
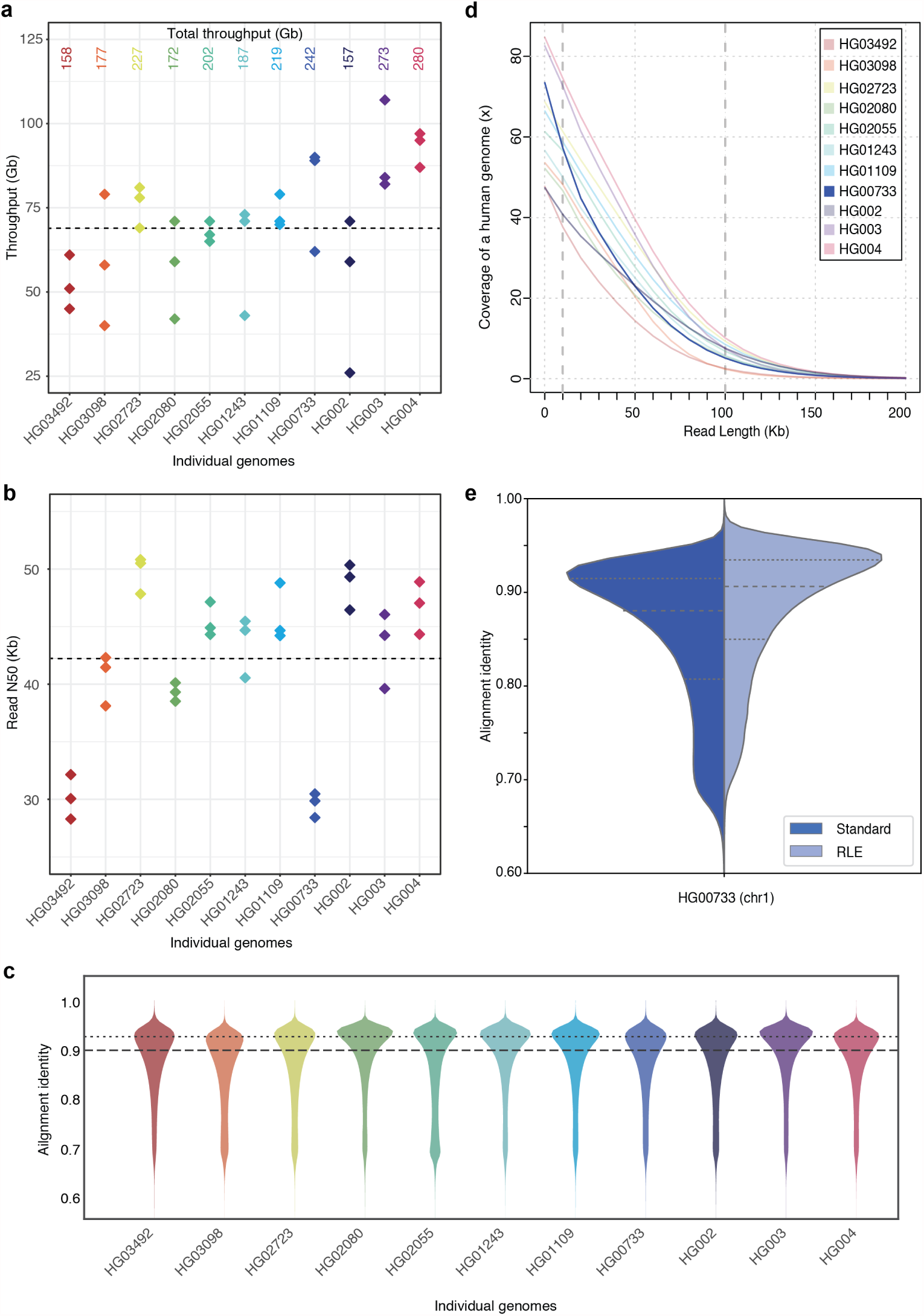
Nanopore sequencing results. **(a)** Throughput in gigabases from each of three flowcells for eleven samples, with total throughput at top. **(b)** Read N50s for each flowcell. **(c)** Alignment identities against GRCh38. Medians in a, b and c shown by dashed lines, dotted line in c is mode. **(d)** Genome coverage as a function of read length. Dashed lines indicate coverage at 10 and 100 Kb. HG00733 is bolded as an example. **(e)** Alignment identity for standard and run-length encoded (RLE) reads. Data for HG00733 chromosome 1 are shown. Dashed lines denote quartiles.

### Shasta: assembling a human genome from nanopore reads in under 6 hours

To assemble the genomes, we developed a new *de novo* assembly algorithm, Shasta. Shasta was designed to be orders of magnitude faster and cheaper at assembling a human-scale genome from nanopore reads than the Canu assembler used in our earlier work [17]. A detailed description of algorithms and computational techniques used is provided in the Online Methods section. Here we summarize key points:

- During most Shasta assembly phases, reads are stored in a homopolymer-compressed (HPC) form using *Run-Length Encoding* (RLE) [23, 24, 25]. In this form, identical consecutive bases are collapsed, and the base and repeat count are stored. For example, GATTTACCA would be represented as (`GATACA`, 113121). This representation is insensitive to errors in the length of homopolymer runs, thereby addressing the dominant error mode for Oxford Nanopore reads [11]. As a result, assembly noise due to read errors is decreased, and significantly higher identity alignments are facilitated (Fig. 1e).
- A *marker representation* of reads is also used, in which each read is represented as the sequence of occurrences of a predetermined, fixed subset of short *k*-mers (*marker representation*) in its run-length representation.
- A modified *MinHash* [26, 27] scheme is used to find candidate pairs of overlapping reads, using as *MinHash* features consecutive occurrences of *m* markers (default *m* = 4).
- Optimal alignments in marker representation are computed for all candidate pairs. The computation of alignments in marker representation is very efficient, particularly as various banded heuristics are used.
- A *Marker Graph* is created in which each vertex represents a marker found to be aligned in a set of several reads. The marker graph is used to assemble sequence after undergoing a series of simplification steps.
- The assembler runs on a single machine with a large amount of memory (typically 1-2 TB for a human assembly). All data structures are kept in memory, and no disk I/O takes place except for initial loading of the reads and final output of assembly results.

To validate Shasta, we compared it against three contemporary assemblers: Wtdbg2 [28], Flye [29] and Canu [30]. We ran all four assemblers on available read data from two diploid human samples, HG00733 and HG002, and one haploid human sample, CHM13. HG00733 and HG002 were part of our collection of eleven samples, and data for CHM13 came from the T2T consortium [31].

Canu consistently produced the most contiguous assemblies, with contig NG50s of 39.0, 31.3, and 85.8 Mb, for samples HG00733, HG002, and CHM13, respectively (Fig. 2a). Flye was the second most contiguous, with contig NG50s of 24.2, 24.9, and 34.2 Mb, for the same samples. Shasta was next with contig NG50s of 20.3, 19.3, and 37.8 Mb. Wtdbg2 produced the least contiguous assemblies, with contig NG50s of 14.5, 12.2, and 13.6 Mb.

**Figure 2:**
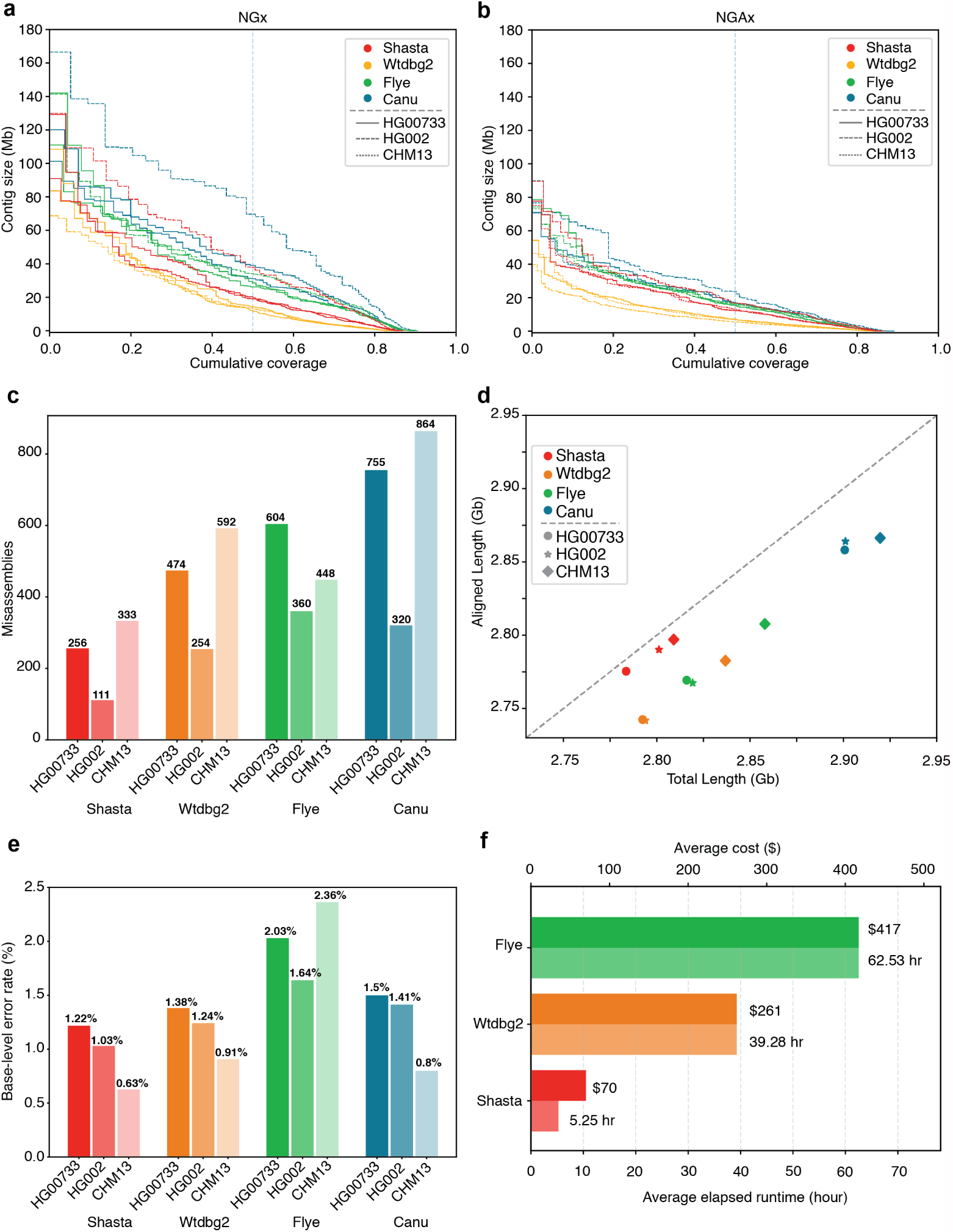
Assembly results for four assemblers and three human samples, before polishing. **(a)** NGx plot showing contig length distribution. The intersection of each line with the dashed line is the NG50 for that assembly. **(b)** NGAx plot showing the distribution of *aligned* contig lengths. Each horizontal line represents an aligned segment of the assembly unbroken by a misassembly or unmappable sequence with respect to GRCh38. The intersection of each line with the dashed line is the aligned NGA50 for that assembly. **(c)** Misassembly counts for regions outside of centromeres, segmental duplications and, for HG002, known SVs. **(d)** Total generated sequence length vs. total aligned sequence length (against GRCh38). **(e)** Balanced base-level error rates for assembled sequences. **(f)** Average runtime and cost for assemblers (Canu not shown).

Conversely, aligning the samples to GRCh38 and evaluating with QUAST [32], Shasta had between 3.6 to 7.9x fewer misassemblies per assembly than the other assemblers (Supplementary Tables 4 and 5). Breaking the assemblies at these misassemblies and unaligned regions with respect to GRCh38, we observe much smaller absolute variation in contiguity (Fig. 2b, avg. NGA50s (Mb): Canu 18.5, Flye 15.2, Shasta 13.7, Wtdbg2 6.4). These results imply that Shasta trades some contiguity for a smaller overall misassembly rate vs. Canu and Flye. However, a substantial fraction of the misassemblies identified likely reflect SVs with respect to GRCh38. To address this we discounted misassemblies within centromeres and known segmental duplications, which are enriched in SVs, and, in the case of HG002, a set of known SVs [33]; we still observe between 1.3 and 2.9x fewer misassemblies in Shasta relative to the other assemblers (Fig. 2c).

For HG00733 and CHM13 we examined a library of available bacterial artificial chromosome (BAC) assemblies (see Online Methods). The BACs were largely targeted at known segmental duplications (473 of 520 BACs lie within 10 Kb of a known duplication). Examining the subset of BACs for CHM13 and HG00733 that map to unique regions of GRCh38 (see Online Methods), we find Shasta contiguously assembles 46 of 47 BACs, with Canu and Flye performing similarly (Supplementary Table 6). In the full set we observe that Canu (401) and Flye (280) contigiously assemble a larger subset of these BACs than Shasta (131) and Wtdbg2 (107), confirming the notion that Shasta is relatively conservative in these duplicated regions (Supplementary Table 7). Examining the fraction of contiguously assembled BACs of those attempted (that is, having a substantial, unique overlap with only a single a contig in the assembly), we can derive a proxy to the specificity of these assemblies. In this regard, Canu (0.88), Shasta (0.87) and Flye (0.84) perform similarly, with Wtdbg2 (0.65) the outlier.

Canu consistently assembled the largest genomes (avg. 2.91 Gb), followed by Flye (avg. 2.83 Gb), Wtdbg2 (avg. 2.81 Gb) and Shasta (avg. 2.80 Gb). We would expect the vast majority of this assembled sequence to map to another human genome. Discounting unmapped sequence, the differences are smaller: Canu produced an avg. 2.86 Gb of mapped sequence per assembly, followed by Shasta (avg. 2.79 Gb), Flye (avg. 2.78 Gb) and Wtdbg2 (avg. 2.76 Gb) (Fig. 2d; see Online Methods). Again, this analysis supports the notion that Shasta is currently relatively conservative vs. its peers, producing the highest proportion of directly mapped assembly per sample.

Shasta produced the most base-level accurate assemblies (avg. balanced error rate 1.13% on diploid and 0.63% on haploid), followed by Wtbdg2 (1.31% on diploid and 0.91% on haploid), Canu (1.46% on diploid and 0.8% on haploid) and Flye (1.84% on diploid and 2.36% on haploid) (Fig. 2e); see Online Methods, Supplementary Table 8.

Shasta, Wtdbg2 and Flye were run on a commercial cloud, allowing us to reasonably compare their cost and run time (Fig. 2e; see Online Methods). Shasta took an average of 5.25 hours to complete each assembly at an average cost of $70 per sample. In contrast, Wtdbg2 took 7.5x longer and cost 3.7x as much, and Flye took 11.9x longer and cost 6.0x as much. The Canu assemblies were run on a large compute cluster, consuming up to $19,000 (estimated) of compute and took around 4-5 days per assembly (see Online Methods, Supplementary Table 9).

### Contiguously assembling MHC haplotypes

The Major Histocompatibility Complex (MHC) region is difficult to resolve using short reads due to its repetitive and highly polymorphic nature [34], but recent efforts to apply long read sequencing to this problem have shown promise [17, 35]. We analyzed the assemblies of CHM13 and HG00733 to see if they spanned the region. For the haploid assembly of CHM13 we find MHC is entirely spanned by a single contig in all 4 assemblers’ output, and most closely resembles the GL000251.2 haplogroup among those provided in GRCh38 (Fig. 3a; Supplementary Fig. 1 and Supplementary Table 10). In the diploid assembly of HG00733 two contigs span the large majority of the MHC for Shasta and Flye, while Canu and Wtdbg2 span the region with one contig (Fig. 3b; Supplementary Fig. 2). However, we note that the chimeric diploid assembly leads to sequences that do not closely resemble any haplogroup (see Online Methods).

**Figure 3:**
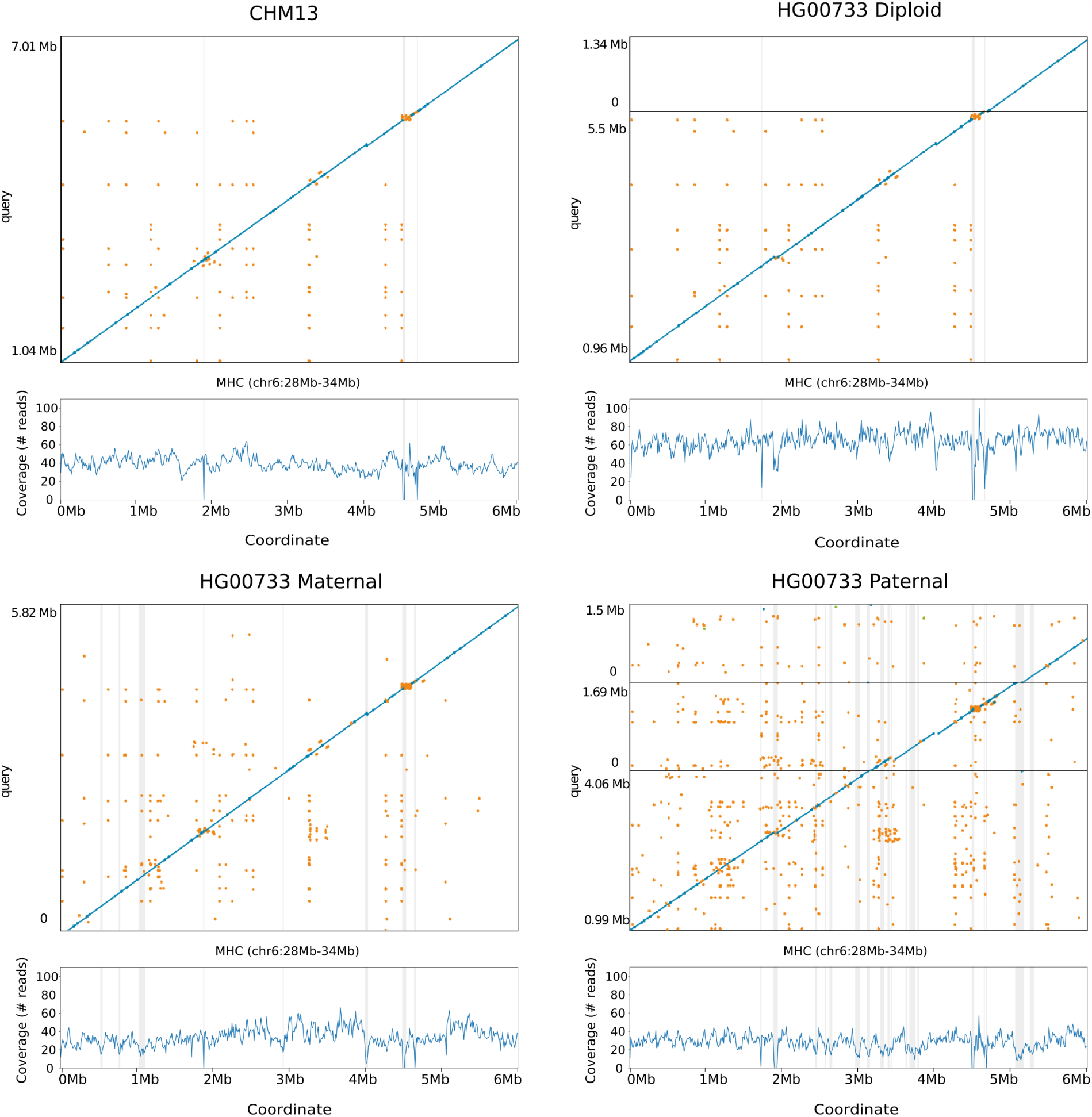
Shasta MHC assemblies vs GRCh38. Unpolished Shasta assembly for CHM13 and HG00733, including HG00733 trio-binned maternal and paternal assemblies. Shaded gray areas are regions in which coverage (as aligned to GRCh38) drops below 20. Horizontal black lines indicate contig breaks. Blue and green describe unique alignments (aligning forward and reverse, respectively) and orange describes multiple alignments.

To attempt to resolve haplotypes of HG00733 we performed trio-binning [36], where we partitioned all the reads for HG00733 into two sets based on likely maternal or paternal lineage and assembled the haplotypes (see Online Methods). For all haplotype assemblies the global contiguity worsened significantly (as the available read data coverage was approximately halved, and further, not all reads could be partitioned), but the resulting misassembly count decreased (Supplementary Table 11). When using haploid trio-binned assemblies, the MHC was spanned by a single contig for the maternal haplotype (Fig. 3c, Supplementary Fig. 3, Supplementary Table 12), with high identity to GRCh38 and having the greatest contiguity and identity with the GL000255.1 haplotype. For the paternal haplotype, low coverage led to discontinuities (Fig. 3d) breaking the region into three contigs.

### Deep neural network based polishing achieves QV30 long-read only polishing accuracy

Accompanying Shasta, we developed a deep neural network based consensus sequence polishing pipeline designed to improve the base-level quality of the initial assembly. The pipeline consists of two modules: MarginPolish and HELEN. MarginPolish uses a banded form of the forward-backward algorithm on a pairwise hidden Markov model (pair-HMM) to generate pairwise alignment statistics from the RLE alignment of each read to the assembly [37]. From these statistics MarginPolish generates a weighted RLE Partial Order Alignment (POA) graph [38] that represents potential alternative local assemblies. MarginPolish iteratively refines the assembly using this RLE POA, and then outputs the final summary graph for consumption by HELEN. HELEN employs a multi-task recurrent neural network (RNN) [39] that takes the weights of the MarginPolish RLE POA graph to predict a nucleotide base and run-length for each genomic position. The RNN takes advantage of contextual genomic features and associative coupling of the POA weights to the correct base and run-length to produce a consensus sequence with higher accuracy.

To demonstrate the effectiveness of MarginPolish and HELEN, we compared them with the state-of-the-art nanopore assembly polishing workflow: four iterations of Racon polishing [40] followed by Medaka [41]. Here MarginPolish is analogous in function to Racon, both using pair-HMM based methods for alignment and POA graphs for initial refinement. Similarly, HELEN is analogous to Medaka, in that both use a deep neural network and both work from summary statistics of reads aligned to the assembly.

Figure 4a and Supplementary Tables 13, 14 and 15 detail error rates for the four methods performed on the HG00733 and CHM13 Shasta assemblies (see Online Methods) using Pomoxis [42]. For the diploid HG00733 sample MarginPolish and HELEN achieve a balanced error rate of 0.501% (QV 23.00), compared to 0.579% (QV 22.37) by Racon and Medaka. For both polishing pipelines, a significant fraction of these errors are likely due to true heterozygous variations. For the haploid CHM13 we restrict comparison to a highly curated X chromosome sequence provided by the T2T consortium [31]. We achieve a balanced error rate of 0.095% (QV 30.22), compared to Racon and Medaka’s 0.127% (QV 28.96).

**Figure 4:**
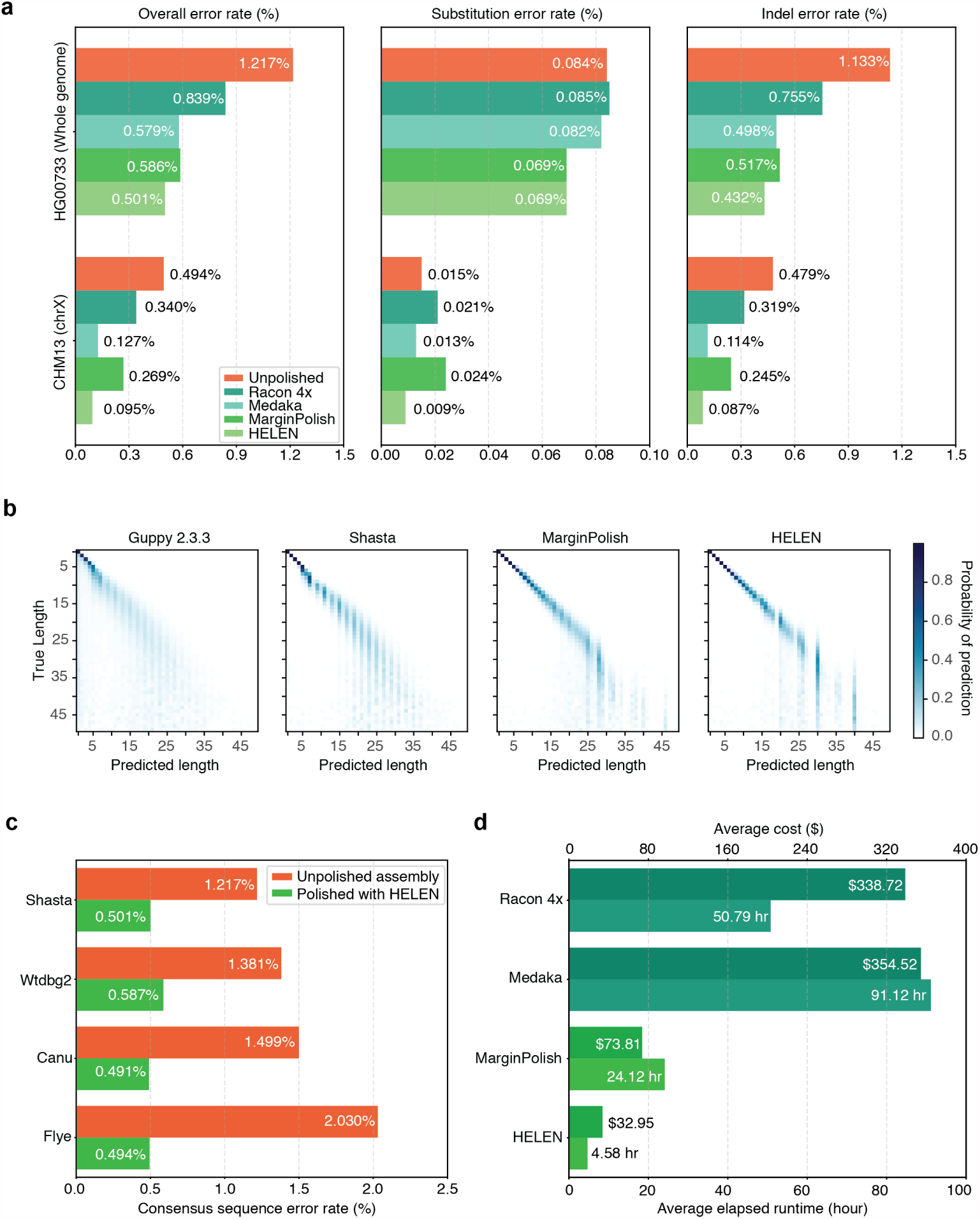
Polishing Results. **(a)** Balanced error rates for the four methods on HG00733 and CHM13. **(b)** Row-normalized heatmaps describing the predicted run-lengths (x-axis) given true run lengths (y-axis) for four steps of the pipeline on HG00733. **(c)** Error rates for MarginPolish and HELEN on four assemblies. **(d)** Average runtime and cost.

For all assemblies, errors were dominated by indel errors, e.g. substitution errors are 6.26x and 9.67x fewer than indels in the MarginPolish and HELEN on HG000733 and CHM13 assemblies, respectively. Many of these errors relate to homopolymer length confusion; Fig. 4b analyzes the homopolymer error rates for various steps of the polishing workflow for HG00733. Each panel shows a heatmap with the true length of the homopolymer run on the y-axis and the predicted run length on the x-axis, with the color describing the likelihood of predicting each run length given the true length. Note that the dispersion of the diagonal steadily decreases. The vertical streaks at high run lengths in the MarginPolish and HELEN confusion-matrix are the result of infrequent numerical and encoding artifacts (see Online Methods, Supplementary Fig. 4)

Figure 4c and Supplementary Table 16 show the overall error rate after running MarginPolish and HELEN on HG00733 assemblies generated by different tools. The consistency in post-polishing error rates is evidence that the models used are not strongly biased towards Shasta and that they can be usefully employed to polish assemblies generated by other tools.

Figure 4d and Supplementary Table 17 describe average runtimes and costs for the methods (see Online Methods). MarginPolish and HELEN cost a combined $108 and took 29 hours of wall-clock time on average, per sample. In comparison Racon and Medaka cost $693 and took 142 wall-clock hours on average, per sample.

### Long-read assemblies contain nearly all human coding genes

To evaluate the accuracy and completeness of an assembled transcriptome we ran the Comparative Annotation Toolkit [43], which can annotate a genome assembly using the human GENCODE [44] reference human gene set (Table 1, Online Methods, Supplementary Tables 18, 19, 20, and 21.).

**Table 1:** CAT transcriptome analysis of human protein coding genes for HG00733 and CHM13.

| Sample | Assembler | Polisher | Genes Found % | Missing Genes | Complete Genes % |
| --- | --- | --- | --- | --- | --- |
| HG00733 | Canu | HELEN | 99.741 | 51 | 67.038 |
|  | Flye | HELEN | 99.405 | 117 | 71.768 |
|  | Wtdbg2 | HELEN | 97.429 | 506 | 66.143 |
|  | Shasta | HELEN | 99.228 | 152 | 68.069 |
|  | Shasta | Medaka | 99.141 | 169 | 66.27 |
| CHM13 | Shasta | HELEN | 99.111 | 175 | 74.202 |
|  | Shasta | Medaka | 99.035 | 190 | 73.836 |

For the HG00733 and CHM13 samples we found that Shasta assemblies polished with MarginPolish and HELEN were close to representing nearly all human protein coding genes, having, respectively, an identified ortholog for 99.23% (152 missing) and 99.11% (175 missing) of these genes. Using the restrictive definition that a coding gene is complete in the assembly only if it is assembled across its full length, contains no frameshifts, and retains the original intron/exon structure, we found that 68.07% and 74.20% of genes, respectively, were complete in the HG00733 and CHM13 assemblies. Polishing the Shasta assemblies alternatively with the Racon-Medaka pipeline achieved similar but uniformly less complete results.

Comparing the MarginPolish and HELEN polished assemblies for HG00733 generated with Flye, Canu and Wtdbg2 to the similarly polished Shasta assembly we found that Canu had the fewest missing genes (just 51), but that Flye, followed by Shasta, had the most complete genes. Wtdbg2 was clearly an outlier, with notably larger numbers of missing genes (506). For comparison we additionally ran BUSCO [45] using the eukaryote set of orthologs on each assembly, a smaller set of 303 expected single-copy genes (Supplementary Tables 22 and 23). We find comparable performance between the assemblies, with small differences largely recapitulating the pattern observed by the larger CAT analysis.

### Comparing to a PacBio HiFi Assembly

We compared the CHM13 Shasta assembly polished using MarginPolish and HELEN with the recently released Canu assembly of CHM13 using PacBio HiFi reads [46]; HiFi reads being based upon circular consensus sequencing technology that delivers significantly lower error rates. The HiFi assembly has lower NG50 (29.0 Mb vs. 41.0 Mb) than the Shasta assembly (Supplementary Fig. 5). Consistent with our other comparisons to Canu, the Shasta assembly also contains a much lower misassembly count (1107) than the Canu based HiFi assembly (8666), a difference which remains after discounting all misassemblies in centromeres and known segmental duplications (314 vs. 893). The assemblies have an almost equal NGAx (~20.0Mb), but the Shasta assembly covers a smaller fraction of GRCh38 (95.28% vs. 97.03%) (Supplementary Fig. 6, Supplementary Table 24). Predictably, the HiFi assembly has ~4.7 fold fewer inserts and ~3.3 fold fewer deletes than the Shasta assembly when aligned to the highly curated X chromosome assembly from v0.6 T2T consortium [31]. Although the HiFi assembly has less indels, both have comparable mismatches with HiFi assembly having ~1.4 fold fewer mismatches (Supplementary Table 25).

### Assembling, polishing and scaffolding 11 human genomes at near chromosome scale

The median NG50 of the 11 Shasta assemblies is 18.5Mb (Fig. 5a), generally below the level required to achieve complete chromosomes; to resolve the remaining contig breaks in the assemblies we scaffolded all of the polished Shasta assemblies with HiC proximity-ligation data using HiRise [47] (see Online Methods). On average, 891 joins were made per assembly. This increased the scaffold NG50s to near chromosome scale, with a median of 129.96 Mb, as shown in Fig. 5a, with additional assembly metrics in Supplementary Table 26. Aligning HG00733 to GRCh38, we find no major rearrangements and all chromosomes are spanned by one or a few contigs (Fig. 5b), with the exception of chrY, which is not present because HG00733 is female. Similar results were observed for HG002 (Supplementary Fig. 7).

**Figure 5:**
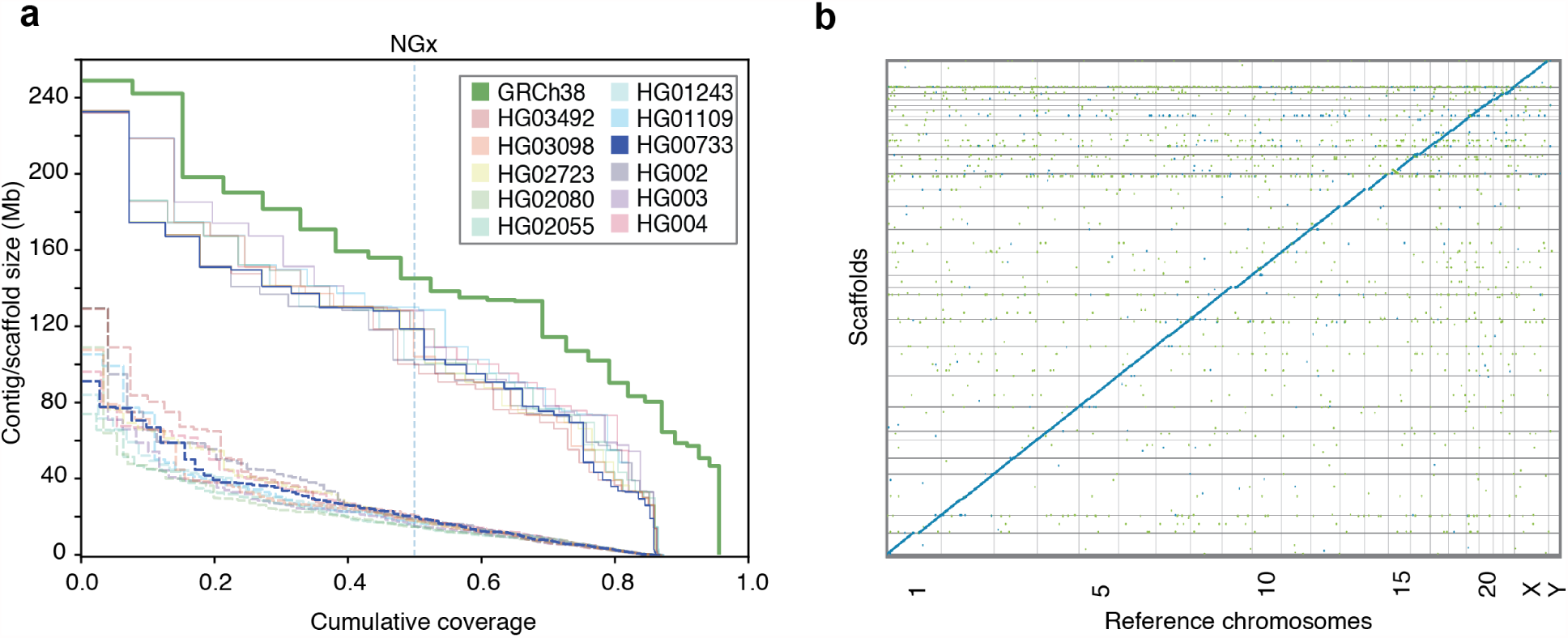
HiRise scaffolding for 11 genomes. **(a)** NGx plots for each of the 11 genomes, before (dashed) and after (solid) scaffolding with HiC sequencing reads, GRCh38 minus alternate sequences is shown for comparison. **(b)** Dot plot showing alignments between the scaffolded HG00733 Shasta assembly and GRCh38 chromosome scaffolds. Blue indicates forward aligning segments, green indicates reverse, with both indicating unique alignments.

## Discussion

In this paper we demonstrate the sequencing and assembly of eleven diverse human genomes in a time and cost efficient manner using a combination of nanopore and proximity ligation sequencing.

The PromethION realizes dramatic improvements in yield per flow cell, allowing the sequencing of each genome with just three flow cells at an average coverage of 63x. This represents a large reduction in associated manual effort and a dramatic practical improvement in parallelism; a single PromethION allows up to 48 flow cells to be run concurrently. Here we completed all 2.3 terabases of nanopore data collection in nine days on one PromethION, running up to 15 flow cells simultaneously (it is now possible to run 48 concurrently). In terms of contemporary long-read sequencing platforms, this throughput is unmatched.

Due to the length distribution of human transposable elements, we found it better to discard reads shorter than 10 Kb to prevent multi-mapping. The Circulomics SRE kit reduced the fraction of reads <10 Kb to around 13%, making the majority usable for assembly. Conversely, the right tail of the read length distribution is long, yielding an average of 6.5x coverage per genome in 100 Kb+ reads. This represents an enrichment of around 7 fold relative to our earlier MinION effort [17]. In terms of assembly, the result was an average NG50 of 18.5 Mb for the 11 genomes, ~3x higher than in that initial effort, and comparable with the best achieved by alternative technologies [12, 48]. We found the addition of HiC sequencing for scaffolding necessary to achieve chromosome scale, making 891 joins on average per assembly. However, our results are consistent with previous modelling based on the size and distribution of large repeats in the human genome, which predicts that an assembly based on 30x coverage of such 100 Kb+ reads would approach the continuity of complete human chromosomes [17, 31].

Relative to alternate long-read and linked-read sequencing, the read identity of nanopore reads has proven lower [11, 17]. However, original reports of 66% identity [11] for the original MinION are now historical footnotes: we observe modal read identity of 92.5%, resulting in QV30 base quality for haploid polished assembly from nanopore reads alone. The accurate resolution of highly repetitive and recently duplicated sequence will depend on long-read polishing, because short-reads are generally not uniquely mappable. Further polishing using complementary data types, including PacBio HiFi reads [48] and 10x Chromium [49], will likely prove useful in achieving QV40+ assemblies.

The advent of third generation technologies has dramatically lowered the cost of high-contiguity long-read *de novo* assembly relative to earlier methods [50]. This cost reduction is still clearly underway. The first MinION human assembly cost ~$40,000 in flow cells and reagents [17]. After a little over a year, the equivalent cost per sample here was ~$6,000. At bulk with current list-pricing, this cost would be reduced to ~$3,500 per genome. It is not unreasonable to expect further yield growth and resulting cost reduction of nanopore and competing platforms such that we foresee $1,000 total sequencing cost high-contiguity *de novo* plant and animal genome assembly being achieved - a milestone that will likely make many ambitious comparative genomic efforts economic [51, 52].

With sequencing efficiency for long-reads improving, computational considerations are paramount in figuring overall time, cost and quality. Simply put, large genome *de novo* assembly will not become ubiquitous if the requirements are weeks of assembly time on large computational clusters. We present three novel methods that provide a pipeline for the rapid assembly of long nanopore reads. Shasta can produce a draft human assembly in around six hours and $70 using widely available commercial cloud nodes. This cost and turnaround time is much more amenable to rapid prototyping and parameter exploration than even the fastest competing method (Wtdbg2), which was on average 7.5x slower and 3.7x more expensive. Connected together, the three tools presented allow a polished assembly to be produced in ~24 hours and for ~$180, against the fastest comparable combination of Wtdbg2/Racon and Medaka which costs 5.4x more and is 5.3x slower while producing measurably worse results in terms of misassemblies, contiguity and base-level accuracy. Substantial further parallelism of polishing, the dominant time component in our current pipeline, is easily possible, such that we are now working to demonstrate a half-day turn around of our complete pipeline. With real-time base calling, a DNA-to-*de novo* assembly could be achieved in less than 96 hours with little difficulty. Such speed could make these techniques practical for screening human genomes for abnormalities in difficult-to-sequence regions.

All three presented computational methods employ run-length encoding of reads. By operating on homopolymer-compressed nucleotide sequences, we mitigate effects of the dominant source of error in nanopore reads [53] and enable the use of different models for addressing alignment and run-length estimation orthogonally.

Shasta produces a notably more conservative assembly than competing tools, trading greater correctness for contiguity and total produced sequence. For example, the ratio of total length to aligned length is relatively constant for all other assemblers, where approximately 1.6% of sequence produced does not align across the three evaluated samples. In contrast, on average just 0.38% of Shasta’s sequence does not align to GRCh38, representing a more than 4x reduction in unaligned sequence. Additionally, we note substantially lower misassembly counts, resulting in much smaller differences between the raw NGx and corrected NGAx values. Shasta also produces substantially more base-level accurate assemblies than the other competing tools.

MarginPolish and HELEN provide a consistent improvement of base quality over all tested assemblers, with more accurate results than the current state-of-the-art long read polishing workflow. We note the marginalization over alignments performed by MarginPolish as a likely source of this improvement, reinforcing conclusions from previous work [1].

We have assembled and compared haploid, trio-binned and diploid samples. Trio binned samples show great promise for haplotype assembly, for example contiguously assembling an MHC haplogroup, but the halving of effective coverage resulted in ultimately less contiguous human assemblies with higher base-error rates than the related, chimeric diploid assembly. This can potentially be rectified by merging the haplotype assemblies to produce a pseudo-haplotype or increasing sequencing coverage. Indeed the improvements in contiguity and base accuracy in CHM13 over the diploid samples illustrate what can be achieved with higher coverage of a haploid sample. We believe that one of the most promising directions for the assembly of diploid samples is the integration of phasing into the assembly algorithm itself, as pioneered by others [16, 54, 55]. We anticipate that the novel tools we’ve described here are suited for this next step: the Shasta framework is well placed for producing phased assemblies over structural variants, MarginPolish is built off of infrastructure designed to phase long reads [1], and the HELEN model could be improved to include haplotagged features for the identification of heterozygous sites.

## Acknowledgements

The authors are grateful for support from the following individuals. Dan Turner, David Stoddart, Androo Markham, and Jonathan Pugh (ONT) provided advice on method development and basecalling. Chris Wright (ONT) provided advice for Medaka. Daniel Garalde, and Rosemary Dokos (ONT) provided advice on the PromethION for parallelized DNA sequencing and basecalling.

The authors are grateful to Amazon Web Services (AWS) for hosting the data via their AWS Public Dataset Program.

AP and SK were supported by the Intramural Research Program of the National Human Genome Research Institute, National Institutes of Health. This work utilized the computational resources of the NIH HPC Biowulf cluster (https://hpc.nih.gov).

Sidney Bell and Charlotte Weaver from Chan Zuckerberg Initiative (CZI) provided support on development and documentation. CZI further supported this effort by funding the usage of Amazon Web Services (AWS) for the project.

Certain commercial equipment, instruments, or materials are identified to specify adequately experimental conditions or reported results. Such identification does not imply recommendation or endorsement by the National Institute of Standards and Technology, nor does it imply that the equipment, instruments, or materials identified are necessarily the best available for the purpose.

This work was supported, in part, by the National Institutes of Health (award numbers: 5U54HG007990, 5T32HG008345-04, 1U01HL137183, R01HG010053, U01HL137183, and U54HG007990 to BP and DH; R01HG010329 to SRS and DH), by Oxford Nanopore Research Grant SC20130149 (MA), and the Howard Hughes Medical Institute (DH).

## Online Methods

### Sample selection

The goal of sample selection was to select a set of individuals that collectively captured the maximum amount of weighted allelic diversity [56]. To do this, we created a list of all low-passage lymphoblastoid cell lines that are part of a trio available from the 1000 Genomes Project collection [57] (We selected trios to allow future addition of pedigree information, and low-passage line to minimize acquired variation). In some cases, we considered the union of parental alleles in the trios due to not having genotypes for the offspring. Let a weighted allele be a variant allele and its frequency in the 1000 Genomes Project Phase 3 VCF. We selected the first sample from our list that contained the largest sum of frequencies of weighted alleles, reasoning that this sample should have the largest expected fraction of variant alleles in common with any other randomly chosen sample. We then removed the variant alleles from this first sample from the set of variant alleles in consideration and repeated the process to pick the second sample, repeating the process recursively until we had selected seven samples. This set greedily, heuristically optimizes the maximum sum of weighted allele frequencies in our chosen sample subset. We also added the three Ashkenazim Trio samples and the Puerto Rican individual (HG00733). These four samples were added for the purposes of comparison with other studies that are using them [22].

### Cell culture

Lymphoblastoid cultures for each individual were obtained from the Coriell Institute Cell Repository (coriell.org) and were cultured in RPMI 1640 supplemented with 15% fetal bovine serum (Life Technologies). The cells underwent a total of six passages (p3+3). After expansion, cells were harvested by pelleting at 300xg for 5 minutes. Cells were resuspended in 10 ml PBS and a cell count was taken using a BiRad TC20 cell counter. Cells were aliquoted into 50 ml conical tubes containing 50 million cells, pelleted as above and washed with 10 ml PBS before a final pelleting after which the PBS was removed and the samples were flash frozen on dry ice and stored at −80°C until ready for further processing.

### DNA extraction and size-selection

We extracted high-molecular weight (HMW) DNA using the QIAGEN Puregene kit. We followed the standard protocol with some modifications. Briefly, we lysed the cells by adding 3 ml of Cell Lysis Solution per 10 million cells, followed by incubation at 37°C for up to 1 hour. We performed mild shaking intermediately by hand, and avoided vortexing. Once clear, we split the lysate into 3 ml aliquots and added 1 ml of Protein Precipitation Solution to each of the tubes. This was followed by pulse vortexing three times for five seconds each time. We next spun this at 2000 x g for 10 minutes. We added the supernatant from each tube to a new tube containing 3 ml of isopropanol, followed by 50x inversion. The HMW DNA precipitated and formed a dense thread-like jelly. We used a disposable inoculation loop to extract the DNA precipitate. We then dipped the DNA precipitate, while it was on the loop, into ice-cold 70% ethanol. After this, the DNA precipitate was added to a new tube containing 50-250 *µ*l 1x TE buffer. The tubes were heated at 50°C for 2 hours and then left at room temperature overnight to allow resuspension of the DNA. The DNA was then quantified using Qubit and NanoDrop.

We used the Circulomics Short Read Eliminator (SRE) kit to deplete short-fragments from the DNA preparation. We size-selected 10 *µ*g of DNA using the Circulomics recommended protocol for each round of size-selection.

### Nanopore sequencing

We used the SQK-LSK109 kit and its recommended protocol for making sequencing libraries. We used 1 *µ*g of input DNA per library. We prepared libraries at a 3x scale since we performed a nuclease flush on every flow cell, followed by the addition of a fresh library.

We used the standard PromethION scripts for sequencing. At around 24 hours, we performed a nuclease flush using the ONT recommended protocol. We then re-primed the flow cell, and added a fresh library corresponding to the same sample. After the first nuclease flush, we restarted the run setting the voltage to −190 mV. We repeated the nuclease flush after another around 24 hours (i.e. around 48 hours into sequencing), re-primed the flow cell, added a fresh library, and restarted the run setting the run voltage to −200 mV.

We performed basecalling using Guppy v.2.3.5 on the PromethION tower using the GPUs. We used the MinION DNA flipflop model (`dna_r9.4.1_450bps_flipflop.cfg`), as recommended by ONT.

### Chromatin Crosslinking and Extraction from Human Cell Lines

We thawed the frozen cell pellets and washed them twice with cold PBS before resuspension in the same buffer. We transferred Aliquots containing five million cells by volume from these suspensions to separate microcentrifuge tubes before chromatin crosslinking by addition of paraformaldehyde (EMS Cat. No. 15714) to a final concentration of one percent. We briefly vortexed the samples and allowed them to incubate at room temperature for fifteen minutes. We pelleted the crosslinked cells and washed them twice with cold PBS before thoroughly resuspending in lysis buffer (50 mM Tris-HCl, 50 mM NaCl, 1 mM EDTA, 1% SDS) to extract crosslinked chromatin.

### The Hi-C Method

We bound the crosslinked chromatin samples to SPRI beads, washed three times with SPRI wash buffer (10 mM Tris-HCl, 50 mM NaCl, 0.05% Tween-20), and digested by DpnII (20 U, NEB Catalog No. R0543S) for 1 hour at 37°C in an agitating thermal mixer. We washed the bead-bound samples again before incorporation of Biotin-11-dCTP (ChemCyte Catalog No. CC-6002-1) by DNA Polymerase I, Klenow Fragment (10 U, NEB Catalog No. M0210L) for thirty minutes at 25°C with shaking. Following another wash, we carried out blunt-end ligation by T4 DNA Ligase (4000 U, NEB Catalog No. M0202T) with shaking overnight at 16°C. We reversed the chromatin crosslinks, digested the proteins, eluted the samples by incubation in crosslink reversal buffer (5 mM CaCl 2, 50 mM Tris-HCl, 8% SDS) with Proteinase K (30 *µ*g, Qiagen Catalog No. 19133) for fifteen minutes at 55°C followed by forty-five minutes at 68°C.

### Sonication and Illumina Library Generation with Biotin Enrichment

After SPRI bead purification of the crosslink-reversed samples, we transferred DNA from each to Covaris® microTUBE AFA Fiber Snap-Cap tubes (Covaris Cat. No. 520045) and sonicated to an average length of 400 ± 85 bp using a Covaris® ME220 Focused-Ultrasonicator™. Temperature was held stably at 6°C and treatment lasted sixty-five seconds per sample with a peak power of fifty watts, ten percent duty factor, and two-hundred cycles per burst. The average fragment length and distribution of sheared DNA was determined by capillary electrophoresis using an Agilent® FragmentAnalyzer 5200 and HS NGS Fragment Kit (Agilent Cat. No. DNF-474-0500). We ran sheared DNA samples twice through the NEBNext® Ultra™ II DNA Library Prep Kit for Illumina® (Catalog No. E7645S) End Preparation and Adaptor Ligation steps with custom Y-adaptors to produce library preparation replicates. We purified ligation products via SPRI beads before Biotin enrichment using Dynabeads® MyOne™ Streptavidin C1 beads (ThermoFisher Catalog No. 65002). We performed indexing PCR on streptavidin beads using KAPA HiFi HotStart ReadyMix (Catalog No. KK2602) and PCR products were isolated by SPRI bead purification. We quantified the libraries by Qubit™ 4 fluorometer and FragmentAnalyzer 5200 HS NGS Fragment Kit (Agilent Cat. No. DNF-474-0500) before pooling for sequencing on an Illumina HiSeq X at Fulgent Genetics.

### Analysis methods

#### Read alignment identities

To generate the identity violin plots (Fig. 1c/e) we aligned all the reads for each sample and flowcell to GRCh38 using `minimap2` [23] with the `map-ont` preset. Using a custom script `get_summary_stats.py` in the repository https://github.com/rlorigro/nanopore_assembly_and_polishing_assessment, we parsed the alignment for each read and enumerated the number of matched (*N*_=_), mismatched (*N*_*X*_), inserted (*N*_*I*_), and deleted (*N*_*D*_) bases. From this, we calculated *alignment identity* as *N*_=_*/*(*N*_=_ + *N*_*X*_ + *N*_*I*_ + *N*_*D*_). These identities were aggregated over samples and plotted using the `seaborn` library with the script `plot_summary_stats.py` in the same repository. This method was used to generate both Figure 1c and Figure 1e. For Figure 1e, we selected reads from HG00733 flowcell1 aligned to GRCh38 chr1. The “Standard” identities are used from the original reads/alignments. To generate identity data for the “RLE” portion, we extracted the reads above, run-length encoded the reads and chr1 reference, and followed the alignment and identity calculation process described above. Sequences were run-length encoded using a simple script github.com/rlorigro/runlength_analysis/blob/master/runlength_encode_fasta.py) and aligned with minimap2 using the `map-ont` preset and `–k 19`.

#### Base-level error-rate analysis with Pomoxis

We analyzed the base-level error-rates of the assemblies using the `assess_assembly` tool of Pomoxis toolkit developed by Oxford Nanopore Technology https://github.com/nanoporetech/pomoxis. The assess assembly tool is tailored to compute the error rates in a given assembly compared to a truth assembly. It reports an identity error rate, insertion error rate, deletion error rate, and an overall error rate. The identity error rate indicates the number of erroneous substitutions, the insertion error rate is the number of incorrect insertions, and the deletion error rate is the number of deleted bases averaged over the total aligned length of the assembly to the truth. The overall error rate is the sum of the identity, insertion, and deletion error rates. For the purpose of simplification, we used the indel error rate, which is the sum of insertion and deletion error rates.

The `assess_assembly` script takes an input assembly and a reference assembly to compare against. The assessment tool chunks the reference assembly to 100 Kb regions and aligns it back to the input assembly to get a trimmed reference. Next, the input is aligned to the trimmed reference sequence with the same alignment parameters to get an input assembly to the reference assembly alignment. The total aligned length is the sum of the lengths of the trimmed reference segments where the input assembly has an alignment. The total aligned length is used as the denominator while averaging each of the error categories to limit the assessment in only correctly assembled regions. Then the tool uses `stats_from_bam`, which counts the number of mismatch bases, insert bases, and delete bases at each of the aligned segments and reports the error rate by averaging them over the total aligned length.

The Pomoxis section in Supplementary Notes describe the commands we ran to perform this assessment.

**Table 2:** The truth assembly files with download URLs.

| Sample name | Region | File type | URL |
| --- | --- | --- | --- |
| HG002 | Whole genome | fasta | HG002_GRCh38_h1.fa |
|  |  | bed | HG002_GRCh38.bed |
| HG00733 | Whole genome | fasta | hg00733_truth_assembly.fa |
| CHM13 | Whole genome | fasta | CHM13_truth_assembly.fa |
|  | Chr-X | fasta | CHRX_CHM13_truth_assembly.fa |

#### Truth assemblies for base-level error-rate analysis

We used HG002, HG00733, and CHM13 for base-level error-rate assessment of the assembler and the polisher. These three assemblies have high-quality assemblies publicly available, which are used as the ground truth for comparison. Two of the samples, HG002 and HG00733, are diploid samples; hence, we picked one of the two possible haplotypes as the truth. The reported error rate of HG002 and HG00733 include some errors arising due to the zygosity of the samples. The complete hydatidiform mole sample CHM13 is a haploid human genome which is used to assess the applicability of the tools on haploid samples. We have gathered and uploaded all the files we used for assessment in one place: https://console.cloud.google.com/storage/browser/kishwar-helen/truth_assemblies/.

To generate the HG002 truth assembly, we gathered the publicly available Genome-in-a-bottle (GIAB) high-confidence variant set (VCF) against GRCh38 reference sequence. Then we used `bedtools` to create an assembly (FASTA) file from the GRCh38 reference and the high-confidence variant set. We got two files using this process for each of the haplotypes, and we picked one randomly as the truth. All the diploid HG002 assembly is compared against this one chosen assembly. GIAB also provides a bed file annotating high-confidence regions where the called variants are highly precise and sensitive. We used this bed file with `assess_assembly` to ensure that we compare the assemblies only in the high confidence regions.

The HG00733 truth is from the publicly available phased PacBio high-quality assembly of this sample [58]. We picked phase0 as the truth assembly and acquired it from NCBI under accession `GCA_003634895.1`. We note that the assembly is phased but not haplotyped, such that portions of phase0 will include sequences from both parental haplotypes and is not suitable for trio-binned analyses. Furthermore, not all regions were fully phased; regions with variants that are represented as some combination of both haplotypes will result in lower QV and a less accurate truth.

For CHM13, we used the v0.6 release of CHM13 assembly by the T2T consortium [31]. The reported quality of this truth assembly in Q-value is QV 39. One of the attributes of this assembly is chromosome X. As reported by the T2T assembly authors, chromosome X of CHM13 is the most complete (end-to-end) and high-quality assembly of any human chromosome. We obtained the chromosome X assembly, which is the highest-quality truth assembly (QV >= 40) we have.

## QUAST / BUSCO

To quantify contiguity, we primarily depended on the tool QUAST [32]. QUAST identifies misassemblies as major rearrangement events in the assembly relative to the reference. For our assemblies, we quantified all contiguity stats against GRCh38, using autosomes plus chromosomes X and Y only. We report the total misassemblies given that their relevant “size” descriptor was greater than 1 Kb, as is the default behavior in QUAST. QUAST provides other contiguity statistics in addition to misassembly count, notably total length and total aligned length as reported in Figure 2d. To determine total aligned length (and unaligned length), QUAST performs collinear chaining on each assembled contig to find the best set of non-overlapping alignments spanning the contig. This process contributes to QUAST’s misassembly determination. We consider unaligned sequence to be the portions of the assembled contigs which are not part of this best set of non-overlapping alignments. All statistics are recorded in Supplementary Table 4. For all QUAST analyses, we used the flags `min-identity` 80 and `fragmented`.

QUAST also produces an NGAx plot (similar to an NGx plot) which shows the aligned segment size distribution of the assembly after accounting for misassemblies and unalignable regions. The intermediate segment lengths that would allow NGAx plots to be reproduced across multiple samples on the same axis (as is shown in Figure 2b) are not stored, so we created a GitHub fork of QUAST to store this data during execution: https://github.com/rlorigro/quast. Finally, the assemblies and the output of QUAST were parsed to generate figures with an NGx visualization script, `ngx_plot.py`, found at github.com/rlorigro/nanopore_assembly_and_polishing_assessment/.

BUSCO [45] is a tool which quantifies the number of Benchmarking Universal Single-Copy Orthologs present in an assembly. We ran BUSCO via the option within QUAST, comparing against the *eukaryota* set of orthologs from OrthoDB v9.

### Misassembly assessments

To analyze the QUAST-reported misassemblies for different regions of the genome, we gathered the known segmental duplication (SD) regions [7], centromeric regions for GRCh38, and known regions in GRCh38 with structural variation for HG002 from GIAB [33]. We used a python script `quast_sv_extractor.py` that compares each reported misassembly of QUAST to the SD, SV and centromeric regions and discounts any misassembly that overlaps with these regions. The `quast_sv_extractor.py` script can be found at https://github.com/kishwarshafin/helen/blob/master/modules/python/helper/.

The segmental duplication regions of GRCh38 defined in the `ucsc.collapsed.sorted.segdups` file can be downloaded from https://github.com/mvollger/segDupPlots/.

The defined centromeric regions of GRCh38 for all chromosomes are used from the available summary at https://www.ncbi.nlm.nih.gov/grc/human.

For GIAB HG002, known SVs for GRCh38 are available in `NIST_SVs_Integration_v0.6/` under ftp://ftp-trace.ncbi.nlm.nih.gov/giab/ftp/data/AshkenazimTrio/analysis/. We used Tier1+2 bed file availabe in the GIAB ftp site.

### Trio-binning

We performed trio-binning on two samples HG002 and HG00733 [36]. For HG00733, we obtained the parental read sample accessions (HG00731, HG00732) from 1000 genome database. Then we counted k-mers with meryl to create maternal and paternal k-mer sets. Based on manual examination of the k-mer count histograms to determine an appropriate threshold, we excluded k-mers occurring less than 6 times for maternal set and 5 times for paternal set. We subtracted the paternal set from the maternal set to get k-mers unique to the maternal sample and similarly derived unique paternal k-mer set. Then for each read, we counted the number of occurrences of unique maternal and paternal k-mers and classified the read based on the highest occurrence count. During classification, we avoided normalization by k-mer set size. This resulted in 35.2x maternal, 37.3x paternal, and 5.6x unclassified for HG00733. For HG002, we used the Illumina data for the parental samples (HG003, HG004) from GIAB project [22]. We counted k-mers using meryl and derived maternal paternal sets using the same protocol. We filtered k-mers that occur less than 25 times in both maternal and paternal sets. The classification resulted in 24x maternal, 23x paternal, and 3.5x unknown. The commands and data source are detailed in the Supplementary Notes.

### Transcript analysis with comparative annotation toolkit

We ran the Comparative Annotation Toolkit [43] to annotate the polished assemblies in order to analyze how well Shasta assembles transcripts and genes. Each assembly was individually aligned to the GRCh38 reference assembly using Cactus [59] to create the input alignment to CAT. The GENCODE [60] V30 annotation was used as the input gene set. CAT was run in the transMap mode only, without Augustus refinement, since the goal was only to evaluate the quality of the projected transcripts. All transcripts on chromosome Y were excluded from the analysis since some samples lacked a Y chromosome.

### Run-Length Confusion Matrix

To generate run-length confusion matrices from reads and assemblies, we run-length encoded (RLE) the assembly/read sequences and reference sequences using a purpose-built python script, `measure_runlength_distribution_from_fasta.py`. The script requires a reference and sequence file, and can be found in the GitHub repo https://github.com/rlorigro/runlength_analysis/. The RLE nucleotides were aligned to the RLE reference nucleotides with `minimap2`. As RLE sequences cannot have identical adjacent nucleotides, the number of unique k-mers is diminished with respect to standard sequences. As `minimap2` uses empirically determined sizes for seed k-mers, we used a k-mer size of 19 to approximately match the frequency of the default size (15) used by the presets for standard sequences. For alignment of reads and assemblies we used the `map-ont` and `asm20` presets respectively.

By iterating through the alignments, each match position in the cigar string (mismatched nucleotides are discarded) was used to find a pair of lengths (*x, y*) such that *x* is a predicted length and *y* is the true (reference) length. For each pair, we updated a matrix which contains the frequency of every possible pairing of prediction vs truth, from length 1bp to 50bp. Finally, this matrix is normalized by dividing each element by the sum of the observations for its true run length, 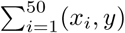, and plotted as a heatmap. Each value represents the probability of predicting a length for a given true length.

### Runtime and Cost Analysis

Our runtime analysis was generated with multiple methods detailing the amount of time the processes took to complete. These methods include the unix command time and a home-grown resource tracking script which can be found in the https://github.com/rlorigro/TaskManager repository. We note that the assembly and polishing methods have different resource requirements, and do not all fully utilize available CPUs, GPUs, and memory over the program’s execution. As such, we report runtimes using wall clock time and the number of CPUs the application was configured to use, but do not convert to CPU hours. Costs reported in the figures are the product of the runtime and AWS instance price. Because portions of some applications do not fully utilize CPUs, cost could potentially be reduced by running on a smaller instance which would be fully utilized, and runtime could be reduced by running on a larger instance which can be fully utilized for some portion of execution. We particularly note the long runtime of Medaka and found that for most of the total runtime, only a single CPU was used. Lastly, we note that data transfer times are not reported in runtimes. Some of the data required or generated exceeds hundreds of gigabytes, which could be potentially significant in relation to the runtime of the process. Notably, the images generated by MarginPolish and consumed by HELEN were often greater than 500 GB in total.

All recorded runtimes are reported in the supplement. For Shasta, times were recorded to the tenth of the hour. All other runtimes were recorded to the minute. All runtimes reported in figures were run on the Amazon Web Services cloud platform (AWS).

Shasta runtime reported in Fig. 2f was determined by averaging across all 12 samples. Wtdbg2 runtime was determined by summing runtimes for wtdbg2 and wtpoa-cns and averaging across the HG00733, HG002, and CHM13 runs. Flye runtime was determined by averaging across the HG00733, HG002, and CHM13 runs. Precise Canu runtimes are not reported, as they were run on the NIH Biowulf cluster. Each run was restricted to nodes with 28 cores (56 hyperthreads) (2×2680v4 or 2×2695v3 Intel CPUs) and 248GB of RAM or 16 cores (32 hyperthreads) (2×2650v2 Intel CPUs) and 121GB of RAM. Full details of the cluster are available at https://hpc.nih.gov. The runs took between 219 and 223 thousand CPU hours (4-5 wall-clock days). No single job used more than 80GB of RAM/12 CPUs. We find the r5.4xlarge ($1.008 per hour) to be the cheapest AWS instance type possible considering this resource usage, which puts estimated cost between $18,000 and $19,000 per genome.

For MarginPolish, we recorded all runtimes, but used various thread counts that did not always fully utilize the instance’s CPUs. The runtime reported in the figure was generated by averaging across 8 of the 12 samples, selecting runs that used 70 CPUs (of the 72 available on the instance). These samples this was true for were GM24385, HG03492, HG01109, HG02055, HG02080, HG01243, HG03098, and CHM13. Runtimes for read alignments used by MarginPolish were not recorded. Because MarginPolish requires an aligned BAM, we found it unfair to not report this time in the figure as it is a required step in the workflows for MarginPolish, Racon, and Medaka. As a proxy for the unrecorded read alignment time used to generate BAMs for MarginPolish, we added the average alignment time recorded while aligning reads in preparation for Medaka runs. We note that the alignment for MarginPolish was done by piping output from `minimap2` directly into `samtools sort`, and piping this into `samtools view` to filter for primary and supplementary reads. Alignment for Medaka was done using `mini_align`, which is a wrapper for `minimap2` bundled in Medaka that simultaneously sorts output.

Reported HELEN runs were performed on GCP except for HG03098, but on instances that match the AWS instance type `p2.8xlarge` in both CPU count and GPU (NVIDIA Tesla P100). As such, the differences in runtime between the platforms should be negligible, and we have calculated cost based on the AWS instance price for consistency. The reported runtime is the sum of time taken by `call_consensus.py` and `stitch.py`. Unannotated runs were performed on UCSC hardware.

Racon runtimes reflect the sum of four series of read alignment and polishing. The time reported in the figure is the average of the runtime of this process run on the Shasta assembly for HG00733, HG002, and CHM13.

Medaka runtime was determined by averaging across the HG00733, HG002, and CHM13 runs after running Racon 4× on the Shasta assembly. We again note that this application in particular did not fully utilize the CPUs for most of the execution, and in the case of HG00733 appeared to hang and was restarted. The plot includes the average runtime from read alignment using `minialign`; this is separated in the tables in the supplementary results. We ran Medaka on an `x1.16xlarge` instance, which had more memory than was necessary. When determining cost, we chose to price the run based on the cheapest AWS instance type that we could have used accounting for configured CPU count and peak memory usage (`c5n.18xlarge`). This instance could have supported 8 more concurrent threads, but as the application did not fully utilize the CPUs we find this to be a fair representation.

### Assembly of MHC

Each of the 8 GRCh38 MHC haplotypes were aligned using `minimap2` (with preset `asm20`) to whole genome assemblies to identify spanning contigs. These contigs were then extracted from the genomic assembly and used for alignment visualization. For dot plots, Nucmer 4.0 [61] was used to align each assembler’s spanning contigs to the standard chr6:28000000-34000000 MHC region, which includes 500Mb flanks. Output from this alignment was parsed with Dot [62] which has a web-based GUI for visualization. All defaults were used in both generating the input files and drawing the figures. Coverage plots were generated from reads aligned to chr6, using a script, `find_coverage.py`, located at (github.com/rlorigro/nanopore_assembly_and_polishing_assessment/).

The best matching alt haplotype (to Shasta, Canu, and Flye) was chosen as a reference haplotype for quantitative analysis. Haplotypes with the fewest supplementary alignments across assemblers were top candidates for QUAST analysis. Candidates with comparable alignments were differentiated by identity. The highest contiguity/identity MHC haplotype was then analyzed with QUAST using `–min-identity 80`. For all MHC analyses regarding Flye, the unpolished output was used.

### BAC Analysis

At a high level, the BAC analysis was performed by aligning BACs to each assembly, quantifying their resolution, and calculating identity statistics on those that were fully resolved.

We obtained 341 BACs for CHM13 [63, 64] and 179 for HG00733 [7] (complete BAC clones of `VMRC62`), which had been selected primarily by targeting complex or highly duplicated regions. We performed the following analysis on the full set of of BACs (for CHM13 and HG00733), and a subset selected to fall within unique regions of the genome. To determine this subset, we selected all BACs which are greater than 10 Kb away from any segmental duplication, resulting in 16 of HG00733 and 31 of CHM13. This subset represents simple regions of the genome which we would expect all assemblers to resolve.

For the analysis, BACs were aligned to each assembly with the command `minimap2 –secondary=no -t 16 -ax asm20 assembly.fasta bac.fasta > assembly.sam` and converted to a PAF-like format which describes aligned regions of the BACs and assemblies. Using this, we calculated two metrics describing how resolved each BAC was: *closed* is defined as having 99.5% of the BAC aligned to a single locus in the assembly; *attempted* is defined as having an alignment of at least 5 Kb with at least 90% identity to only a single assembly contig. If multiple such alignments exist to a single contig, it counts as attempted; if multiple such alignments exist to different contigs, it does not count as attempted. We furthermore calculate median and mean identities (using alignment identity metric described above) of the closed BACs. The code for this can be found at https://github.com/skoren/bacValidation

### Shasta

The following describes Shasta version 0.1.0 (https://github.com/chanzuckerberg/shasta/releases/tag/0.1.0) which was used throughout our analysis. All runs were done on an AWS `x1.32xlarge` instance (1952 GB memory, 128 virtual processors). The runs used the Shasta recommended options for best performance (`–memoryMode filesystem –memoryBacking 2M`). Rather than using the distributed version of the release, the source code was rebuilt locally for best performance as recommended by Shasta documentation.

### Run-length encoding of input reads

Shasta represents input reads using run-length encoding. The sequence of each input read is represented as a sequence of bases, each with a repeat count that says how many times each of the bases is repeated. Such a representation has previously been used in biological sequence analysis [23, 24, 25].

For example, the following read

~~~
CGATTTAAGTTA
~~~

is represented as follows using run-length encoding:

~~~
CGATAGTA
11132121
~~~

Using run-length encoding makes the assembly process less sensitive to errors in the length of homopolymer runs, which are the most common type of errors in Oxford Nanopore reads. For example, consider these two reads:

~~~
CGATTTAAGTTA
CGATTAAGGGTTA
~~~

Using their raw representation above, these reads can be aligned like this:

~~~
CGATTTAAG--TTA
CGATT-AAGGGTTA
~~~

Aligning the second read to the first required a deletion and two insertions. But in run-length encoding, the two reads become:

~~~
CGATAGTA
11132121
CGATAGTA
11122321
~~~

The sequence portions are now identical and can be aligned trivially and exactly, without any insertions or deletions:

~~~
CGATAGTA
CGATAGTA
~~~

The differences between the two reads only appear in the repeat counts:

~~~
11132121
11122321
* *
~~~

The Shasta assembler uses one byte to represent repeat counts, and as a result it only represents repeat counts between 1 and 255. If a read contains more than 255 consecutive bases, it is discarded on input. In the data we have analyzed so far such reads are extremely rare.

### Some properties of base sequences in run-length encoding

- In the sequence portion of the run-length encoding, consecutive bases are always distinct. If they were not, the second one would be removed from the run-length encoded sequence, while increasing the repeat count for the first one.
- With ordinary base sequences, the number of distinct *k*-mers of length *k* is 4^*k*^. But with run-length base sequences, the number of distinct *k*-mers of length *k* is 4 × 3^*k−*1^. This is a consequence of the previous bullet.
- The run-length sequence is generally shorter than the raw sequence, and cannot be longer. For a long random sequence, the number of bases in the run-length representation is 3/4 of the number of bases in the raw representation.

### Markers

Even with run-length encoding, error in input reads are still frequent. To further reduce sensitivity to errors, and also to speed up some of the computational steps in the assembly process, the Shasta assembler also uses a read representation based on *markers*. Markers are occurrences in reads of a pre-determined subset of short *k*-mers. By default, Shasta uses for this purpose *k*-mers with *k* = 10 in run-length encoding, corresponding to an average approximately 13 bases in raw read representation.

Just for the purposes of illustration, consider a description using markers of length 3 in run-length encoding. There is a total 4 × 3^2^ = 36 distinct such markers. We arbitrarily choose the following fixed subset of the 36, and we assign an id to each of the kmers in the subset as follows:

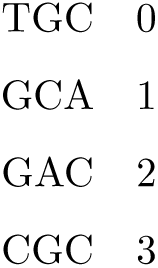

Consider now the following portion of a read in run-length representation (here, the repeat counts are irrelevant and so they are omitted):

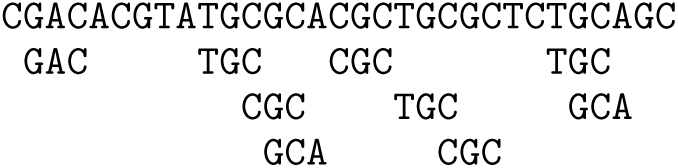

Occurrences of the *k*-mers defined in the table above are shown and define the markers in this read. Note that markers can overlap. Using the marker ids defined in the table above, we can summarize the sequence of this read portion as follows:

~~~
2 0 3 1 3 0 3 0 1
~~~

This is the marker representation of the read portion above. It just includes the sequence of markers occurring in the read, not their positions.

Note that the marker representation loses information, as it is not possible to reconstruct the complete initial sequence from the marker representation. This also means that the marker representation is insensitive to errors in the sequence portions that don’t belong to any markers.

The Shasta assembler uses a random choice of the *k*-mers to be used as markers. The length of the markers *k* is controlled by assembly parameter `Kmers.k` with a default value of 10. Each *k*-mer is randomly choosen to be used as a marker with probability determined by assembly parameter `Kmers.probability` with a default value of 0.1. With these default values, the total number of distinct markers is approximately 0.1 *×* 4 *×* 3^9^ *≈* 7900.

The only constraint used in selecting *k*-mers to be used as markers is that if a *k*-mer is a marker, its reverse complement should also be a marker. This makes it easy to construct the marker representation of the reverse complement of a read from the marker representation of the original read. It also ensures strand symmetry in some of the computational steps.

It is possible that the random selection of markers is not optimal, and that it may be best to select the markers based on their frequency in the input reads or other criteria. These possibilities have not yet been investigated.

Fig. 6 shows the run-length representation of a portion of a read and its markers, as displayed by the Shasta http server.

**Figure 6:**
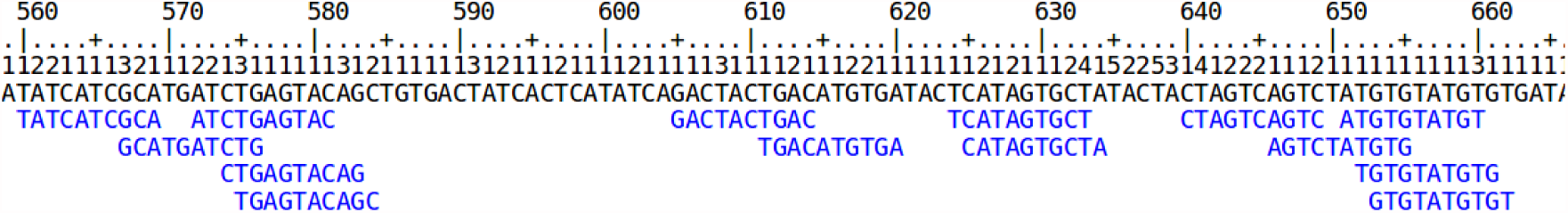
Markers aligned to a run length encoded read.

### Marker alignments

The marker representation of a read is a sequence in an alphabet consisting of the marker ids. This sequence is much shorter than the original sequence of the read, but uses a much larger alphabet. For example, with default Shasta assembly parameters, the marker representation is 10 times shorter than the run-length encoded read sequence, or about 13 times shorter than the raw read sequence. Its alphabet has around 8000 symbols, many more than the 4 symbols that the original read sequence uses.

Because the marker representation of a read is a sequence, we can compute an alignment of two reads directly in marker representation. Computing an alignment in this way has two important advantages:

- The shorter sequences and larger alphabet make the alignment much faster to compute.
- The alignment is insensitive to read errors in the portions that are not covered by any marker.

For these reasons, the marker representation is orders of magnitude more efficient than the raw base representation when computing read alignments. Fig. 7 shows an example alignment matrix.

**Figure 7:**
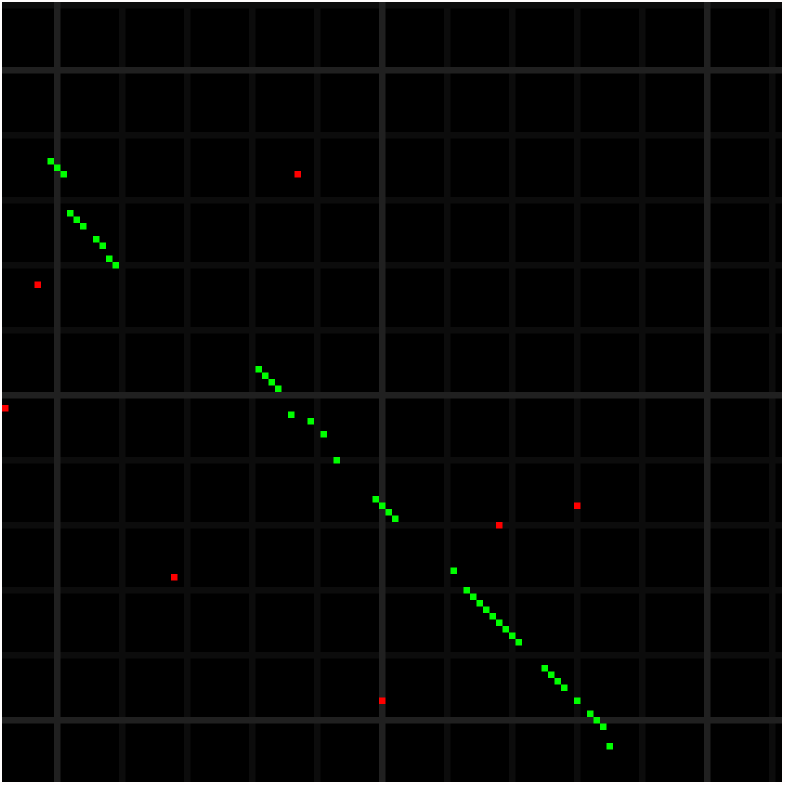
A marker alignment represented as a dot-plot. Elements that are identical between the two sequences are displayed in green or red - the ones in green are the ones that are part of the optimal alignment computed by the Shasta assembler. Because of the much larger alphabet, matrix elements that are identical between the sequences but are not part of the optimal alignment are infrequent. Each alignment matrix element here corresponds on average to a 13 *×* 13 block in the alignment matrix in raw base sequence.

### Computing optimal alignments in marker representation

To compute the (likely) optimal alignment (example highlighted in green in Fig. 7), the Shasta assembler uses a simple alignment algorithm on the marker representations of the two reads to be aligned. It effectively constructs an optimal path in the alignment matrix, but using some ‘banding’ heuristics to speed up the computation:

- The maximum number of markers that an alignment can skip on either read is limited to a maximum, under control of assembly parameter `Align.maxSkip` (default value 30 markers, corresponding to around 400 bases when all other Shasta parameters are at their default). This reflects the fact that Oxford Nanopore reads can often have long stretches in error. In the alignment matrix shown in Fig. 7, there is a skip of about 20 markers (2 light grey squares) following the first 10 aligned markers (green dots) on the top left.
- The maximum number of markers that an alignment can skip at the beginning or end of a read is limited to a maximum, under control of assembly parameter `Align.maxTrim` (default value 30 markers, corresponding to around 400 bases when all other Shasta parameters are at their default). This reflects the fact that Oxford Nanopore reads often have an initial or final portion that is not usable. These first two heuristics are equivalent to computing a reduced band of the alignment matrix.
- To avoid alignment artifacts, marker *k*-mers that are too frequent in either of the two reads being aligned are not used in the alignment computation. For this purpose, the Shasta assembler uses a criterion based on absolute number of occurrences of marker k-mers in the two reads, although a relative criterion (occurrences per Kb) may be more appropriate. The current absolute frequency threshold is under control of assembly parameter `Align.maxMarkerFrequency` (default 10 occurrences).

Using these techniques and with the default assembly parameters, the time to compute an optimal alignment is ~10^*−*3^ − 10^*−*2^ seconds in the Shasta implementation as of release 0.1.0 (April 2019). A typical human assembly needs to compute 10^8^ read alignments which results in a total compute time ~10^5^ − 10^6^ seconds, or ~10^3^ − 10^4^ seconds of elapsed time (~ 1-3 hours) on a machine with 128 virtual processors. This is one of the most computationally expensive portions of a Shasta assembly. Some additional optimizations are possible in the code that implement this computation, and may be implemented in future releases.

### Finding overlapping reads

Even though computing read alignments in marker representation is fast, it still is not feasible to compute alignments among all possible pairs of reads. For a human size genome with ~10^6^ − 10^7^ reads, the number of pairs to consider would be ~10^12^ − 10^14^, and even at ~10^*−*3^ seconds per alignment the compute time would be ~10^9^ − 10^11^ seconds, or ~ 10^7^ − 10^9^ seconds elapsed time (~ 10^2^ − 10^4^ days) when using 128 virtual processors.

Therefore some means of narrowing down substantially the number of pairs to be considered is essential. The Shasta assembler uses for this purpose a slightly modified MinHash [26, 27] scheme based on the marker representation of reads.

In overview, the MinHash algorithm takes as input a set of items each characterized by a set of features. Its goal is to find pairs of the input items that have a high Jaccard similarity index -that is, pairs of items that have many features in common. The algorithm proceeds by iterations. At each iteration, a new hash table is created and a hash function that operates on the feature set is selected. For each item, the hash function of each of its features is evaluated, and the minimum hash function value found is used to select the hash table bucket that each item is stored in. It can be proven that the probability of two items ending up in the same bucket equals the Jaccard similarity index of the two items - that is, items in the same bucket are more likely to be highly similar than items in different buckets. The algorithm then adds to the pairs of potentially similar items all pairs of items that are in the same bucket.

When all iterations are complete, the probability that a pair of items was found at least once is an increasing function of the Jaccard similarity of the two items. In other words, the pairs found are enriched for pairs that have high similarity. One can now consider all the pairs found (hopefully a much smaller set than all possible pairs) and compute the Jaccard similarity index for each, then keep only the pairs for which the index is sufficiently high. The algorithm does not guarantee that all pairs with high similarity will be found - only that the probability of finding all pairs is an increasing function of their similarity.

The algorithm is used by Shasta with items being oriented reads (a read in either original or reverse complemented orientation) and features being consecutive occurrences of *m* markers in the marker representation of the oriented read. For example, consider an oriented read with the following marker representation:

~~~
18,45,71,3,15,6,21
~~~

If *m* is selected equal to 4 (the Shasta default, controlled by assembly parameter `MinHash.m`), the oriented read is assigned the following features:

~~~
(18,45,71,3)
(45,71,3,15)
(71,3,15,6)
(3,15,6,21)
~~~

From the picture above of an alignment matrix in marker representation, we see that streaks of 4 or more common consecutive markers are relatively common. We have to keep in mind that, with Shasta default parameters, 4 consecutive markers span an average 40 bases in run-length encoding or about 52 bases in the original raw base representation. At a typical error rate around 10%, such a portion of a read would contain on average 5 errors. Yet, the marker representation in run-length space is sufficiently robust that these common “features” are relatively common despite the high error rate. This indicates that we can expect the MinHash algorithm to be effective in finding pairs of overlapping reads.

However, the MinHash algorithm has a feature that is undesirable for our purposes: namely, that the algorithm is good at finding read pairs with high Jaccard similarity index. For two sets *X* and *Y*, the Jaccard similarity index is defined as the ratio:

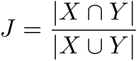

Because the read length distribution of Oxford Nanopore reads is very wide, it is very common to have pairs of reads with very different lengths. Consider now two reads with lengths *n*_*x*_ and *n*_*y*_, with *n*_*x*_ < *n*_*y*_, that overlap exactly over the entire length *n*_*x*_. The Jaccard similarity is in this case given by *n_x_/n_y_ <* 1. This means that, if one of the reads in a pair is much shorter than the other one, their Jaccard similarity will be low even in the best case of exact overlap. As a result, the unmodified MinHash algorithm will not do a good job at finding overlapping pairs of reads with very different length.

For this reason, the Shasta assembler uses a small modification to the MinHash algorithm: instead of just using the minimum hash for each oriented read for each iteration, it keeps all hashes below a given threshold (this is not the same as keeping a fixed number of the lowest hashes for each read). Each oriented read can be stored in multiple buckets, one for each low hash encountered. The average number of low hashes on a read is proportional to its length, and therefore this change has the effect of eliminating the bias against pairs in which one read is much shorter than the other. The probability of finding a given pair is no longer driver by the Jaccard similarity. The modified algorithm is referred to as *LowHash* in the Shasta source code. Note that it is effectively equivalent to an indexing approach in which we index all features with low hash.

The LowHash algorithm is controlled by the following assembly parameters:

- `MinHash.m` (default 4): the number of consecutive markers that define a feature.
- `MinHash.hashFraction`(default 0.01): The fraction of hash values that count as “low”.
- `MinHash.minHashIterationCount` (default 10): The number of iterations.
- `MinHash.maxBucketSize` (default 10): The maximum number of items for a bucket to be considered. Buckets with more than this number of items are ignored. The goal of this parameter is to mitigate the effect of common repeats, which can result in buckets containing large numbers of unrelated oriented reads.
- `MinHash.minFrequency` (default 2): the number of times a pair of oriented reads has to be found to be considered and stored as a possible pair of overlapping reads.

### Initial assembly steps

Initial steps of a Shasta assembly proceed as follows. If the assembly is setup for best performance (`--memoryMode filesystem --memoryBacking 2M` if using the Shasta executable), all data structures are stored in memory, and no disk activity takes place except for initial loading of the input reads, storing of assembly results, and storing a small number of small files with useful summary information.

- Input reads are read from Fasta files and converted to run-length representation.
- *K*-mers to be used as markers are randomly selected.
- Occurrences of those marker *k*-mers in all oriented reads are found.
- The LowHash algorithm finds candidate pairs of overlapping oriented reads.
- A marker alignment is computed for each candidate pair of oriented reads. If the marker alignment contains a minimum number of aligned markers, the pair is stored as an aligned pair. The minimum number of aligned markers is controlled by assembly parameter `Align.minAlignedMarkerCount`.

### Read graph

Using the methods covered so far, an assembly has created a list of pairs of oriented reads, each pair having a plausible marker alignment. How to use this type of information for assembly is a classical problem with a standard solution (Myers, 2005), the *string graph*.

However, the prescriptions in the Myers paper cannot be directly used here, the main reason being that the process used to find pairs of overlapping reads is probabilistic and does not guarantee that all overlapping pairs will be found. Direct application of the Myers approach in this context results in unnecessary breakages in continuity.

The approach currently used in the Shasta assembler is very simple, and can likely be improved. In the current simple approach, the Shasta assembler creates un undirected graph, the *Read Graph*, in which each vertex represents an oriented read (that is, a read in either original orientation or reverse complemented), and an undirected edge between two vertices is created if we have found an alignment between the corresponding oriented reads.

However, the read graph as constructed in this way suffers from high connectivity in repeat regions. Therefore, the Shasta assembler only keeps a *k*-Nearest-Neighbor subset of the edges. That is, for each vertex (oriented read) we only keep the *k* edges with the best alignments (greatest number of aligned markers). The number of edges kept for each vertex is controlled by assembly parameter `ReadGraph.maxAlignmentCount`, with a default value of 6. Note that, despite the *k*-Nearest-Neighbor subset, it remains possible for a vertex to have degree more than *k*.

Note that each read contributes two vertices to the read graph, one in its original orientation, and one in reverse complemented orientation. Therefore the read graph contains two strands, each strand at full coverage. This makes it easy to investigate and potentially detect erroneous strand jumps that would be much less obvious if using approaches with one vertex per read.

An example of one strand is shown in Fig. 8. Even though the graph is undirected, edges that correspond to overlap alignments are drawn with an arrow that points from the prefix oriented read to the suffix one, to represent the direction of overlap. Edges that correspond to containment alignments are drawn in red and without an arrow. Vertices are drawn with area proportional to the length of the corresponding reads.

**Figure 8:**
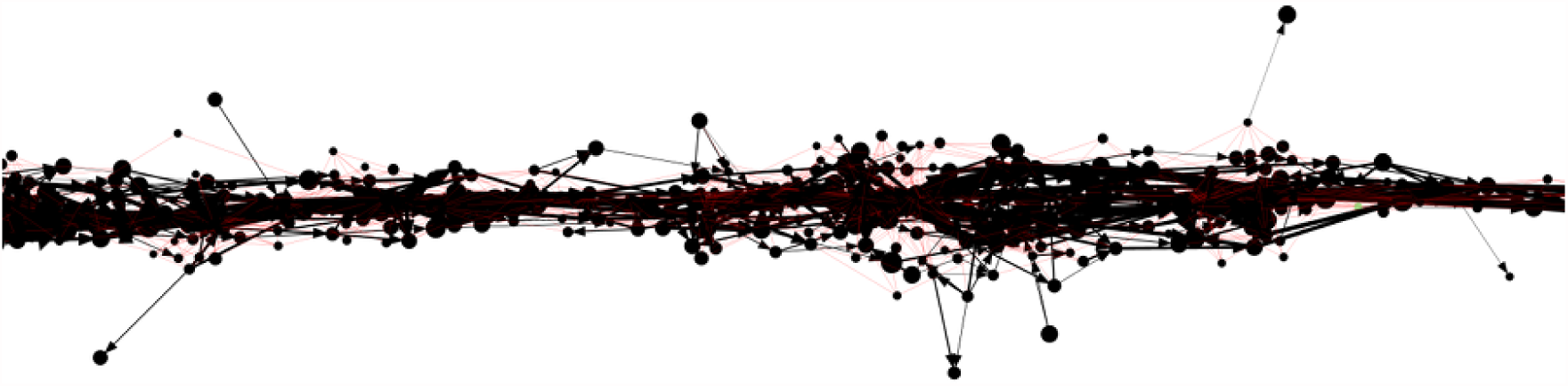
An example of a portion of the read graph, as displayed by the Shasta http server.

The linear structure of the read graph successfully reflects the linear arrangement of the input reads and their origin on the genome being assembled.

However, deviations from the linear structure can occur in the presence of long repeats (Fig. 9), typically for high similarity segment duplications.

**Figure 9:**
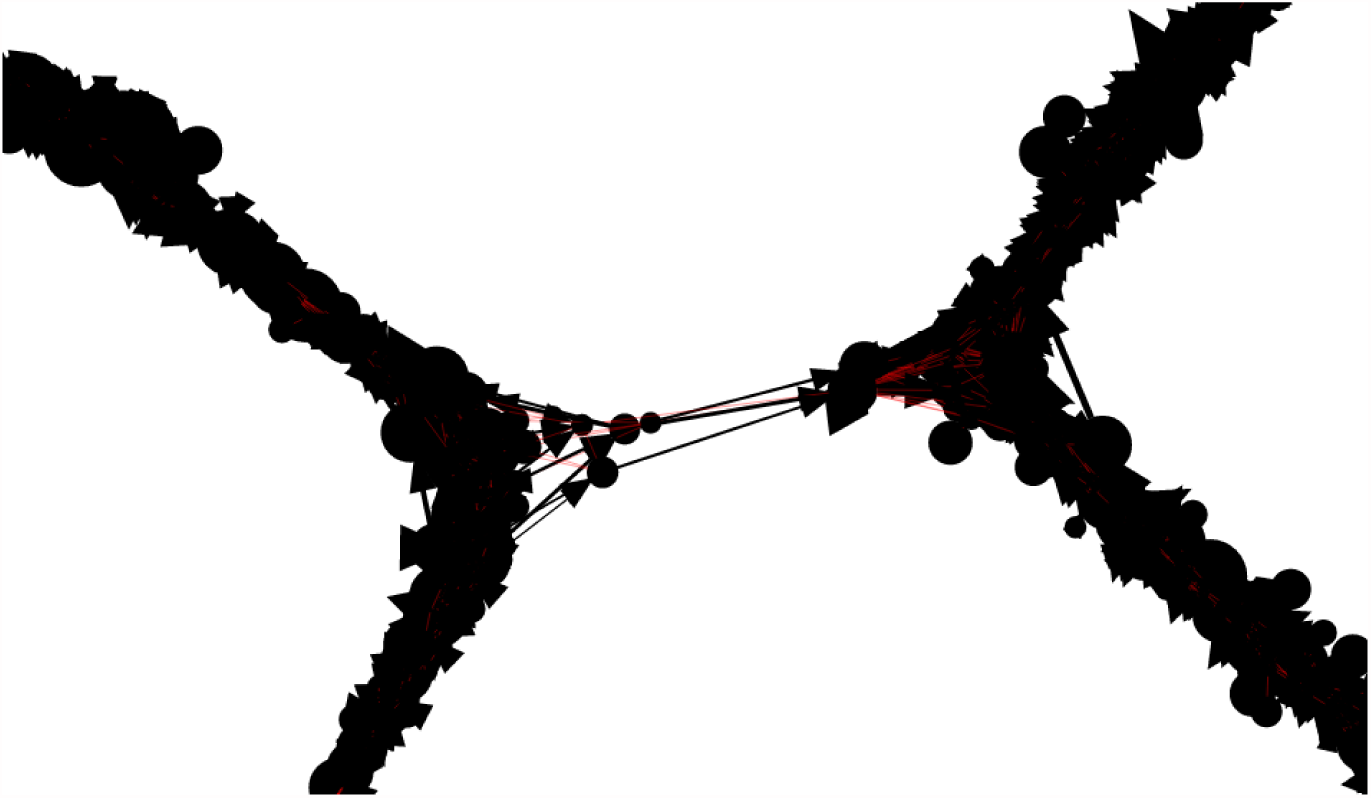
An example of a portion of the read graph showing obviously incorrect connections

The current Shasta implementation does not attempt to remove the obviously incorrect connections. This results in unnecessary breaks in assembly contiguity. Despite this, Shasta assembly contiguity is adequate and comparable to what other, less performant long read assemblers achieve. It is hoped that future Shasta releases will do a better job at handling these situations.

### Marker graph

Consider a read whose marker representation is:

~~~
a b c d e
~~~

We can represent this read as a directed graph that the describes the sequence in which its markers appear (Fig. 10).

**Figure 10:**
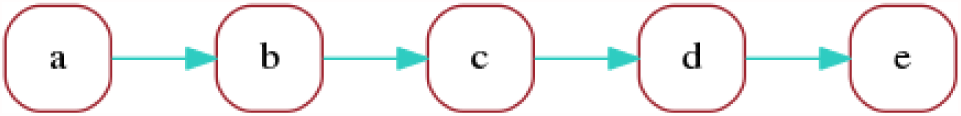
A marker graph representing a single read.

This is not very useful but illustrates the simplest form of a *marker graph* as used in the Shasta assembler. The marker graph is a directed graph in which each vertex represents a marker and each edge represents the transition between consecutive markers. We can associate sequence with each vertex and edge of the marker graph:

- Each vertex is associated with the sequence of the corresponding marker.
- If the markers of the source and target vertex of an edge do not overlap, the edge is associated with the sequence intervening between the two markers.
- If the markers of the source and target vertex of an edge do overlap, the edge is associated with the overlapping portion of the marker sequences.

Consider now a second read with the following marker representation, which differs from the previous one just by replacing marker c with `x`:

~~~
a b x d e
~~~

The marker graph for the two reads is Fig 11(A).

**Figure 11:**
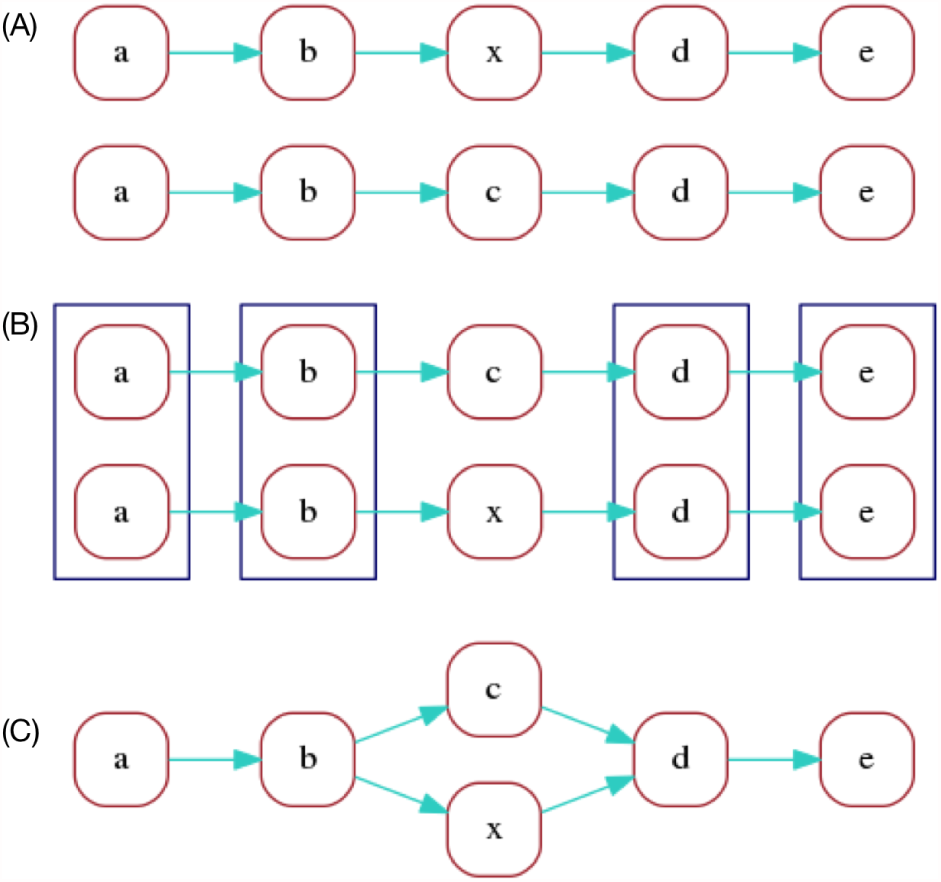
An illustration of marker graph construction for two sequences.

In the optimal alignment of the two reads, markers `a, b, d, e` are aligned. We can redraw the marker graph grouping together vertices that correspond to aligned markers as in Fig 11(B).

Finally, we can merge aligned vertices to obtain a marker graph describing the two aligned reads, shown in Fig 11(C).

Here, by construction each vertex still has a unique sequence associated with it - the common sequence of the markers that were merged (however the corresponding repeat counts can be different for each contributing read). An edge, on the other hand, can have different sequences associated with it, one corresponding to each of the contributing reads. In this example, edges `a->b` and `d->e` have two contributing reads, which can each have distinct sequence between the two markers.

We call coverage of a vertex or edge the number of reads “contributing” to it. In this example, vertices `a, b, d, e` have coverage 2 and vertices `c, x` have coverage 1. Edges `a->b` and `d->e` have coverage 2, and the remaining edges have coverage 1.

The construction of the marker graph was illustrated above for two reads, but the Shasta assembler constructs a global marker graph which takes into account all oriented reads:

- The process starts with a distinct vertex for each marker of each oriented read. Note that at this stage the marker graph is large (~ 2 × 10^10^ vertices for a human assembly using default assembly parameters).
- For each marker alignment corresponding to an edge of the read graph, we merge vertices corresponding to aligned markers.
- Of the resulting merged vertices, we remove those whose coverage in too low or two high, indicating that the contributing reads or some of the alignments involved are probably in error. This is controlled by assembly parameters `MarkerGraph.minCoverage` (default 10) and `MarkerGraph.maxCoverage` (default 100), which specify the minimum and maximum coverage for a vertex to be kept.
- Edges are created. An edge `v0->v1` is created if there is at least a read contributing to both v0 and v1 and for which all markers intervening between `v0` and `v1` belong to vertices that were removed.

Note that this does not mean that all vertices with the same marker sequence are merged - two vertices are only merged if they have the same marker sequence, and if there are at least two reads for which the corresponding markers are aligned.

Given the large number of initial vertices involved, this computation is not trivial. To allow efficient computation in parallel on many threads a lock-free implementation of the disjoint data set data structure [65], is used for merging vertices. Some code changes were necessary to permit large numbers of vertices, as the initial implementation by Wenzel Jakob only allowed for 32-bit vertex ids (https://github.com/wjakob/dset).

### Assembly graph

The Shasta assembly process also uses a compact representation of the marker graph, called the *assembly graph*, in which each linear sequence of edges is replaced by a single edge (Fig. 12).

**Figure 12:**
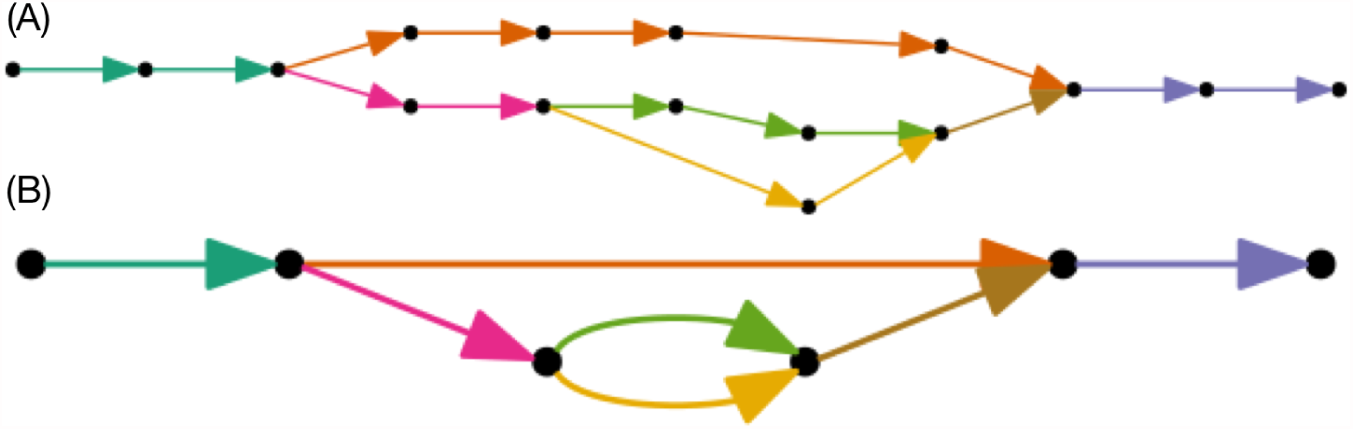
(A) A marker graph with linear sequence of edges colored. (B) The corresponding assembly graph. Colors were chosen to indicate the correspondence to marker graph edges.

The *length* of an edge of the assembly graph is defined as the number of marker graph edges that it corresponds to. For each edge of the assembly graph, an average coverage is also computed, by averaging the coverage of the marker graph edges it corresponds to.

### Using the marker graph to assemble sequence

The marker graph is a partial description of the multiple sequence alignment between reads and can be used to assemble consensus sequence. One simple way to do that is to only keep the “dominant” path in the graph, and then traverse that path from vertex to edge to vertex, assembling run-length encoded sequence as follows:

1. On a vertex, all reads have the same sequence, by construction: the marker sequence associated with the vertex. There is trivial consensus among all the reads contributing to a vertex, and the marker sequence can be used directly as the contribution of the vertex to assembled sequence.
2. For edges, there are two possible situations plus a hybrid case:
  - 2.1. If the adjacent markers overlap, in most cases all contributing reads have the same number of overlapping bases between the two markers, and we are again in a situation of trivial consensus, where all reads contribute the same sequence, which also agrees with the sequence of adjacent vertices. In cases where not all reads are in agreement on the number of overlapping bases, only reads with the most frequent number of overlapping bases are taken into account.
  - 2.2. If the adjacent markers don’t overlap, then each read can have a different sequence between the two markers. In this situation, we compute a multiple sequence alignment of the sequences and a consensus using the spoa library [38] (https://github.com/rvaser/spoa). The multiple sequence alignment is computed constrained at both ends, because all reads contributing to the edge have, by construction, identical markers at both sides.
  - 2.3. A hybrid situation occasionally arises, in which some reads have the two markers overlapping, and some do not. In this case we count reads of the two kinds and discard the reads of the minority kind, then revert to one of the two cases 2.1 or 2.2 above.

This is the process used for sequence assembly by the current Shasta implementation. It requires a process to select and define dominant paths, which is described in the next section. It is algorithmically simple, but its main shortcoming is that it does not use for assembly reads that contribute to the abundant side branches. This means that coverage is lost, and therefore the sequence of assembled accuracy is not as good as it could be if all available coverage was used. Means to eliminate this shortcoming and use information from the side branches of the marker graph could be a subject of future work on the Shasta assembler.

The process described above works with run-length encoded sequence and therefore assembles run-length encoded sequence. The final step to create raw assembled sequence is to compute the most likely repeat count for each sequence position in run-length encoding. After some experimentation, this is currently done by choosing as the most likely repeat count the one that appears the most frequently in the reads that contributed to each assembled position.

A simple Bayesian model for repeat counts resulted in a modest improvement in the quality of assembled sequence. But the model appears to sensitive to calibration errors, and therefore it is not used by default in Shasta assemblies. However, it is used by MarginPolish, as described below.

### Selecting assembly paths in Shasta

The sequence assembly procedure described in the previous section can be used to assemble sequence for any path in the marker graph. This section describes the selection of paths for assembly in the current Shasta implementation. This is done by a series of steps that “remove” edges (but not vertices) from the marker graph until the marker graph consists mainly of linear sections which can be used as the assembly paths. For speed, edges are not actually removed but just marked as removed using a set of flag bits allocated for this purpose in each edge. However, the description below will use the loose term “remove” to indicate that an edge was flagged as removed.

This process consists of the following three steps, described in more detail in the following sections:

- Approximate transitive reduction of the marker graph.
- Pruning of short side branches (leaves).
- Removal of bubbles and super-bubbles.

The last step, removal of bubbles and superbubbles, is consistent with Shasta’s current assembly goal which is to compute a mostly monoploid assembly, at least on short scales.

### Approximate transitive reduction of the marker graph

The goal of this step is to eliminate the side branches in the marker graph, which are the result of errors. Despite the fact that the number of side branches is substantially reduced thanks to the use of run-length encoding, side branches are still abundant. This step uses an approximate transitive reduction of the marker graph which only considers reachability up to a maximum distance, controlled by assembly parameter `MarkerGraph.maxDistance` (default 30 marker graph edges). Using a maximum distance makes sure that the process remains computationally affordable, and also has the advantage of not removing long-range edges in the marker graph, which could be significant.

In detail, the process works as follows. In this description, the edge being considered for removal is the edge v0→v1 with source vertex v0 and target vertex v1. The first two steps are not really part of the transitive reduction but are performed by the same code for convenience.

- All edges with coverage less than or equal to `MarkerGraph.lowCoverageThreshold` are unconditionally removed. The default value for this assembly parameter is 0, so this step does nothing when using default parameters.
- All edges with coverage 1 and for which the only supporting read has a large marker skip are unconditionally removed. The marker skip of an edge, for a given read, is defined as the distance (in markers) between the v0 marker for that read and the v1 marker for the same read. Most marker skips are small, and a large skip is indicative of an artifact. Keeping those edges could result in assembly errors. The marker skip threshold is controlled by assembly parameter `MarkerGraph.edgeMarkerSkipThreshold` (default 100 markers).
- Edges with coverage greater than `MarkerGraph.lowCoverageThreshold` (default 0) and less than `MarkerGraph.highCoverageThreshold` (default 256), and that were not previously removed, are processed in order of increasing coverage. Note that with the default values of these parameters all edges are processed, because edge coverage is stored using one byte and therefore can never be more than 255 (it is saturated at 255). For each edge `v0→v1`, a Breadth-First Search (BFS) in the marker graph is performed starting at source vertex v0 and with a limit of `MarkerGraph.maxDistance` (default 30) edges distance from vertex v0. The BFS is constrained to not use edge `v0→v1`. If the BFS reaches v1, indicating that an alternative path from v0 to v1 exists, edge `v0→v1` is removed. Note that the BFS does not use edges that have already been removed, and so the process is guaranteed not to affect reachability. Processing edges in order of increasing coverage makes sure that low coverage edges the most likely to be removed.

The transitive reduction step is intrinsically sequential and so it is currently performed in sequential code for simplicity. It could be in principle be parallelized, but that would require sophisticated locking of marker graph edges to make sure independent threads don’t step on each other, possibly reducing reachability. However, even with sequential code, this step is not computationally expensive, taking typically only a small fraction of total assembly time.

When the transitive reduction step is complete, the marker graph consists mostly of linear sections composed of vertices with in-degree and out-degree one, with occasional side branches and bubbles or superbubbles, which are handled in the next two phases described below.

### Pruning of short side branches (leaves)

At this stage, a few iterations of pruning are done by simply removing, at each iteration, edge v0→v1 if v0 has in-degree 0 (that is, is a backward-pointing leaf) or v1 has out-degree 0 (that is, is a forward-pointing leaf). The net effect is that all side branches of length (number of edges) at most equal to the number of iterations are removed. This leaves the leaf vertex isolated, which causes no problems. The number of iterations is controlled by assembly parameter `MarkerGraph.pruneIterationCount` (default 6).

### Removal of bubbles and superbubbles

The marker graph now consists of mostly linear section with occasional bubbles or superbubbles [66]. Most of the bubbles and superbubbles are caused by errors, but some of those are due to heterozygous loci in the genome being assembled. Bubbles and superbubbles of the latter type could be used for separating haplotypes (phasing) - a possibility that will be addressed in future Shasta releases. However, the goal of the current Shasta implementation is to create a monoploid assembly at all scales but the very long ones. Accordingly, bubbles and superbubbles at short scales are treated as errors, and the goal of the bubble/superbubble removal step is to keep the most significant path in each bubble or superbubble.

The Fig. 13 shows typical examples of a bubble and superbubble in the marker graph.

**Figure 13:**
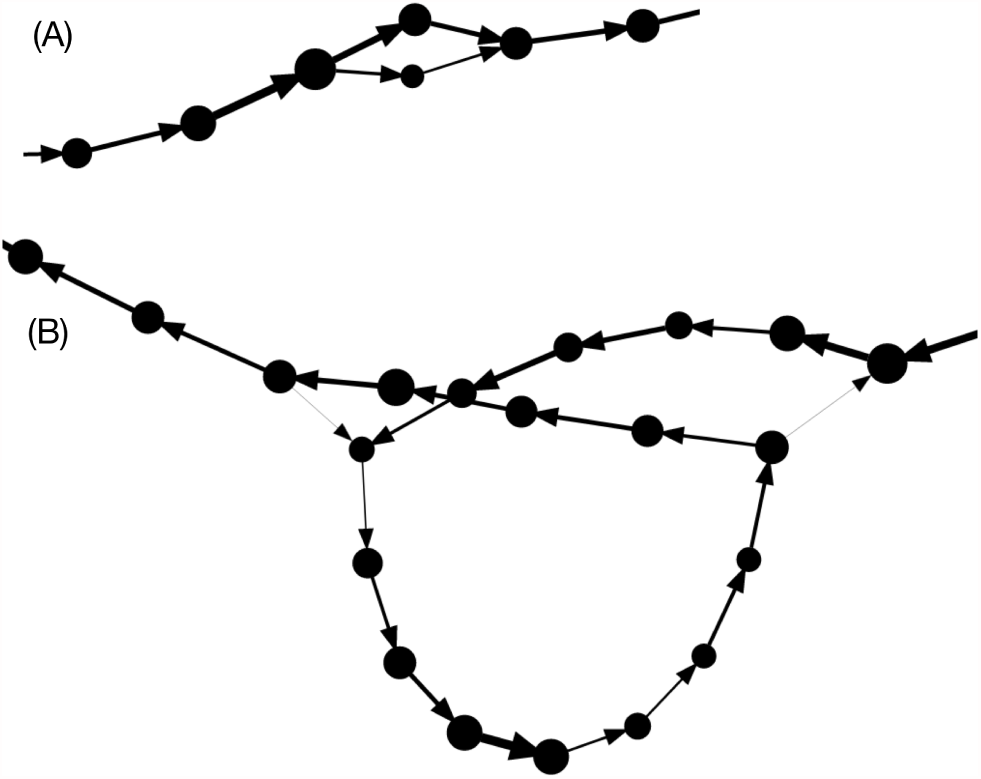
(A) A simple bubble. (B) A superbubble.

The bubble/superbubble removal process is iterative. Early iterations work on short scales, and late iterations fork on longer scales. Each iteration uses a length threshold that controls the maximum number of marker graph edges for features to be considered for removal. The value of the threshold for each iteration is specified using assembly parameter `MarkerGraph.simplifyMaxLength`, which consists of a comma-separated string of integer numbers, each specifying the threshold for one iteration in the process. The default value is 10,100,1000, which means that three iterations of this process are performed. The first iteration uses a threshold of 10 marker graph edges, and the second and third iterations use length thresholds of 100 and 1000 marker graph edges, respectively. The last and largest of the threshold values used determines the size of the smallest bubble or superbubble that will survive the process. The default 1000 markers is equivalent to roughly 13 Kb. To suppress more bubble/superbubbles, increase the threshold for the last iterarion. To see more bubbles/superbubbles, decrease the length threshold for the last iteration, or remove the last iteration entirely.

The goal of the increasing threshold values is to work on small features at first, and on larger features in the later iterations. The choice of `MarkerGraph.simplifyMaxLength` could be application dependent. The default value is a reasonable compromise useful if one desires a mostly monoploid assembly with just some large heterozygous features.

Each iteration consists of two steps. The first removes bubbles and the second removes superbubbles. Only bubbles/superbubbles consisting of features shorter than the threshold for the current iteration are considered:

#### 1. Bubble removal

- An assembly graph corresponding to the current marker graph is created.
- Bubbles are located in which the length of all branches (number of marker graph edges) is no more than the length threshold at the current iteration. In the assembly graph, a bubble appears as a set of parallel edges (edges with the same source and target).
- In each bubble, only the assembly graph edge with the highest average coverage is kept. Marker graph edges corresponding to all other assembly graph edges in the bubble are flagged as removed.

#### 2. Superbubble removal

- An assembly graph corresponding to the current marker graph is created.
- Connected components of the assembly graph are computed, but only considering edges below the current length threshold. This way, each connected component corresponds to a “cluster” of “short” assembly graph edges.
- For each cluster, entries in the cluster are located. These are vertices that have in-edges from a vertex outside the cluster. Similarly, out-edges are located (vertices that have out-edges outside the cluster).
- For each entry/exit pair, the shortest path is computed. However, in this case the “length” of an assembly graph edge is defined as the inverse of its average coverage - that is, the inverse of average coverage for all the contributing marker graph edges.
- Edges on each shortest path are marked as edges to be kept.
- All other edges internal to the cluster are removed.

When all iterations of bubble/superbubble removal are complete, the assembler creates a final version of the assembly graph. Each edge of the assembly graph corresponds to a path in the marker graph, for which sequence can be assembled using the method described above. Note, however, that the marker graph and the assembly graph have been constructed to contain both strands. Special care is taken during all transformation steps to make sure that the marker graph (and therefore the assembly graph) remain symmetric with respect to strand swaps. Therefore, the majority of assembly graph edges come in reverse complemented pairs, of which we assemble only one. It is however possible but rare for an assembly graph to be its own reverse complement.

### High performance computing techniques employed by Shasta

The Shasta assembler is designed to run on a single machine with an amount of memory sufficient to hold all of its data structures (1-2 TB for a human assembly, depending on coverage). All data structures are memory mapped and can be set up to remain available after assembly completes. Note that using such a large memory machine does not substantially increase the cost per CPU cycle. For example, on Amazon AWS the cost per virtual processor hour for large memory instances is no more than twice the cost for laptop-sized instances.

There are various advantages to running assemblies in this way:

- Running on a single machine simplifies the logistics of running an assembly, versus for example running on a cluster of smaller machines with shared storage.
- No disk input/output takes place during assembly, except for loading the reads in memory and writing out assembly results plus a few small files containing summary information. This eliminates performance bottlenecks commonly caused by disk I/O.
- Having all data structures in memory makes it easier and more efficient to exploit parallelism, even at very low granularity.
- Algorithm development is easier, as all data are immediately accessible without the need to read files from disk. For example, it is possible to easily rerun a specific portion of an assembly for experimentation and debugging without any wait time for data structures to be read from disk.
- When the assembler data structures are set up to remain in memory after the assembler completes, it is possible to use the Python API or the Shasta http server to inspect and analyze an assembly and its data structures (for example, display a portion of the read graph, marker graph, or assembly graph).
- For optimal performance, assembler data structures can be mapped to Linux 2 MB pages (“huge pages”). This makes it faster for the operating system to allocate and manage the memory, and improves TLB efficiency. Using huge pages mapped on the `hugetlbfs` filesystem (Shasta executable options `--memoryMode filesystem --memoryBacking 2M`) can result in a significant speed up (20-30%) for large assemblies. However it requires root privilege via `sudo`.

To optimize performance in this setting, the Shasta assembler uses various techniques:

- In most parallel steps, the division of work among threads is not set up in advance but decided dynamically (“Dynamic load balancing”). As a thread finishes a piece of work assigned to it, it grabs another chunk of work to do. The process of assigning work items to threads is lock-free (that is, it uses atomic memory primitives rather than mutexes or other synchronization methods provided by the operating system).
- Most large memory allocations are done via `mmap` and can optionally be mapped to Linux 2 MB pages backed by the Linux `hugetlbfs`. This memory is persistent until the next reboot and is resident (non-pageable). As a result, assembler data structures can be kept in memory and reaccessed repeatedly at very low cost. This facilitates algorithm development (e. g. it allows repeatedly testing a single assembly phase without having to rerun the entire assembly each time or having to wait for data to load) and postprocessing (inspecting assembly data structures after the assembly is complete). The Shasta http server and Python API take advantage of this capability.
- The Shasta code includes a C++ class for conveniently handling these large memory-mapped regions as C++ containers with familiar semantics (`class shasta::MemoryMapped::Vector`).
- In situations where a large number of small vectors are required, a two-pass process is used (`class shasta::MemoryMapped::VectorOfVectors`). In the first pass, one computes the length of each of the vectors. A single large area is then allocated to hold all of the vectors contiguously, together with another area to hold indexes pointing to the beginning of each of the short vectors. In a second pass, the vectors are then filled. Both passes can be performed in parallel and are entirely lock-free. This process eliminates memory allocation overhead that would be incurred if each of the vectors were to be allocated individually.

Thanks to these techniques, Shasta achieves close to 100% CPU utilization during its parallel phases, even when using large numbers of threads. However, a number of sequential phases remain, which typically result in average CPU utilization during a large assembly around 70%. Some of these sequential phases can be parallelized, which would result in increased average CPU utilization and improved assembly performance.

### MarginPolish

Throughout we used MarginPolish (https://github.com/ucsc-nanopore-cgl/MarginPolish) version 1.0.0.

MarginPolish is an assembly refinement tool designed to sum over (marginalize) read to assembly alignment uncertainty. It takes as input a genome assembly and set of aligned reads in BAM format.

It outputs a refined version of the input genome assembly after attempting to correct base-level errors in terms of substitutions and indels (insertions and deletions). It can also output a summary representation of the assembly and read alignments as a weighted partial order alignment graph (POA), which is used by the HELEN neural network based polisher described below.

It was designed and is optimized to work with noisy long ONT reads, although parameterization for other, similar read types is easily possible. It does not yet consider signal-level information from ONT reads. It is also currently a haploid polisher, in that it does not attempt to recognize or represent heterozygous polymorphisms or phasing relationships. For haploid genome assemblies of a diploid genome it will therefore fail to capture half of all heterozygous polymorphisms.

### Algorithm Overview

MarginPolish works in overview as follows:

1. Reads and the input assembly are converted to their run-length encoding (RLE) (see Shasta description above for description and rationale).
2. A restricted, weighted Partial Order Alignment [38] (POA) graph is constructed representing the RLE input assembly and potential edits to it in terms of substitutions and indels.
3. Within identified regions of the POA containing likely assembly errors:
  - A set of alternative sequences representing combinations of edits are enumerated by locally traversing the POA within the region.
  - The likelihood of the existing and each alternative sequence is evaluated given the aligned reads.
  - If an alternative sequence with higher likelihood than the current reference exists then the assembly at the location is updated with this higher likelihood sequence.
4. Optionally, the program loops back to step 2 to repeat the refinement process (by default it loops back once).
5. The modified RLE assembly is expanded by estimating the repeat count of each base given the reads using a simple Bayesian model. The resulting final, polished assembly is output. In addition, a representation of the weighted POA can be output.

### Innovations

Compared to existing tools MarginPolish is most similar to Racon [40], in that they are comparable in speed, both principally use small-parameter HMM like models and both do not currently use signal information. Compared to Racon MarginPolish has some key innovations that we have found to improve polishing accuracy:

- MarginPolish, as with our earlier tool in the Margin series [1], uses the forward-backward and forward algorithms for pair hidden Markov models (HMMs) to sum over all possible pairwise alignments between pairs of sequences instead of the single most probable alignment (Viterbi). Considering all alignments allows more information to be extracted per read.
- The MarginPolish POA construction does not have a read-order dependence. Earlier algorithms for constructing POA graphs have a well known explicit read order dependence that can result in undesirable topologies.
- MarginPolish works in run-length encoded space, which results in considerably less alignment uncertainty and correspondingly improved performance.
- MarginPolish, similarly to Nanopolish [67], evaluates the likelihood of each alternative sequence introduced into the assembly. This improves performance relative to a faster but less accurate algorithm that traces back a consensus sequence through the POA graph.
- MarginPolish employs a simple chunking scheme to break up the polishing of the assembly into overlapping pieces. This results in low memory usage per core and simple parallelism.

Below steps 2, 3 and 5 of the MarginPolish algorithm are described in detail. In addition, the parallelization scheme is described.

### Partial Order Alignment Graph Construction

To create the POA we start with the existing assembled sequence *s* = *s*_1_, *s*_2_,…*s*_*n*_ and for each read *r* = *r*_1_, *r*_2_, …, *r*_*m*_ in the set of reads *R* use the Forward-Backward algorithm with a standard 3-state, affine-gap pair-HMM to derive posterior alignment probabilities using the implementation described in [59]. The parameters for this model are specified in the `polish.hmm` subtree of the JSON formatted parameters file, including `polish.hmm.transitions`, and `polish.hmm.emissions`. Current defaults were tuned via EM [11] of R9.4 ONT reads from and aligned to a bacterial reference; we have observed the parameters for this HMM seem robust to small changes in base-caller versions. The result of running the Forward-backward algorithm is three sets of posterior probabilities:

- Firstly *match probabilities*: the set of posterior match probabilities, each the probability *P* (*r*_*i*_ ◊ *s*_*j*_) that a read base *r*_*i*_ is aligned to a base *s*_*j*_ in *s*.
- Secondly *insertion probabilities*: the set of posterior insertion probabilities, each the probability *P*(*r*_*i*_ ◊ −*j*) that a read base *r*_*i*_ is inserted between two bases *s*_*j*_ and *s*_*j+1*_ in *s*, or, if *j* = 0, inserted before the start of *s*, or, if *j* = *n*, after the end of *s*.
- Thirdly *deletion probabilities*, the set of posterior deletion probabilities, each the probability *P* (*s*_*j*_ ◊ −*r*_*i*_) that a base *s*_*j*_ in *s* is deleted between two read bases *r*_*i*_ and *r*_*i+1*_. (Note, because a read is generally an incomplete observation of *s* we consider the probability that a base in *s* is deleted before the first position or after the last position of a read as 0).

As most probabilities in these three sets are very small and yet to store and compute all the probabilities would require evaluating comparatively large forward and backward alignment matrices we restrict the set of probabilities heuristically as follows:

- We use a banded forward-backward algorithm, as originally described here [68]. To do this we use the original alignment of the read to *s* as in the input BAM file. Given that *s* is generally much longer than each read this allows computation of each forward-backward invocation in time linearly proportional to the length of each read, at the cost of restricting the probability computation to a sub-portion of the overall matrix, albeit one that contains the vast majority of the probability mass.
- We only store posterior probabilities above a threshold (`polish.pairwiseAlignmentParameters.threshold`, by default 0.01), treating smaller probabilities as equivalent as zero.

The result is that these three sets of probabilities are a very sparse subset of the complete sets.

To estimate the posterior probability of a multi-base insertion of a read substring *r*_*i*_, *r*_*i+1*_, … *r*_*k*_ at a given location *j* in *s* involves repeated summation over terms in the forward and backward matrices. Instead to approximate this probability we heuristically use:

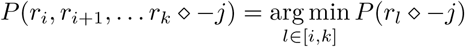

the minimum probability of any base in the multi-base insertion being individually inserted at the location in *s* as a proxy, a probability that is an upper-bound on the actual probability.

Similarly we estimate the posterior probability of a deletion involving more than one contiguous base *s* at a given location in a read using analogous logic. As we store a sparse subset of the single-base insertion and deletion probabilities and given these probability approximations it is easy to calculate all the multi-base indel probabilities with value greater than *t* by linear traversal of the single-based insertion and deletion probabilities after sorting them, respectively, by their read and *s* coordinates. The result of such calculation is expanded sets of insertion and deletion probabilities that include multi-base probabilities.

To build the POA we start from *s*, which we call the *backbone*. The backbone is a graph where each base *s*_*j*_ in *s* corresponds to a node, there are special source and sink nodes (which do not have a base label), and the directed edges connect the nodes for successive bases *s*_*j*_, *s*_*j+1*_ in *s*, from the source node to the node for *s*_1_, and, similarly, from the node for *s*_*n*_ to the sink node.

Each non-source/sink node in the backbone has a separate weight for each possible base *x* ∈ {*A, C, G, T*}. This weight:

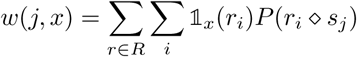

where 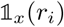 is an indicator function that is 1 if *r*_*i*_ = *x* and otherwise 0, corresponds to the sum of match probabilities of read elements of base *x* being aligned to *s*_*j*_. This weight has a probabilistic interpretation: it is the total number of expected observations of the base *x* in the reads aligned to *s*_*j*_, summing over all possible pairwise alignments of the reads to *s*. It can be fractional because of the inherent uncertainty of these alignments, e.g. we may predict only a 50% probability of observing such a base in a read.

We add *deletion edges*, which connect nodes in the backbone. Indexing the nodes in the backbone from 0 (the source) to the source *n* + 1 (the sink), a deletion edge between positions *j* and *k* in the backbone corresponds to the deletion of bases *j, j* + 1, … *k* − 1 in *s*. Each deletion edge has a weight equal to the sum of deletion probabilities for deletion events that delete the corresponding base(s) in *s*, summing over all possible deletion locations in all reads. Deletions with no weight are not included. Again, this weight has a probabilistic interpretation: it is the expected number of times we see the deletion in the reads, and again it may be fractional.

We represent insertions as nodes labelled with an insertion sequence. Each insertion node has a single incoming edge from a backbone node, and a single outgoing edge to the next backbone node in the backbone sequence. Each insertion is labeled with a weight equal to the sum of probabilities of events that insert the given insertion sequence between the corresponding bases in *s*. The resulting POA is a restricted form of a weighted, directed acyclic graph (Fig. 14(A) shows an example).

**Figure 14:**
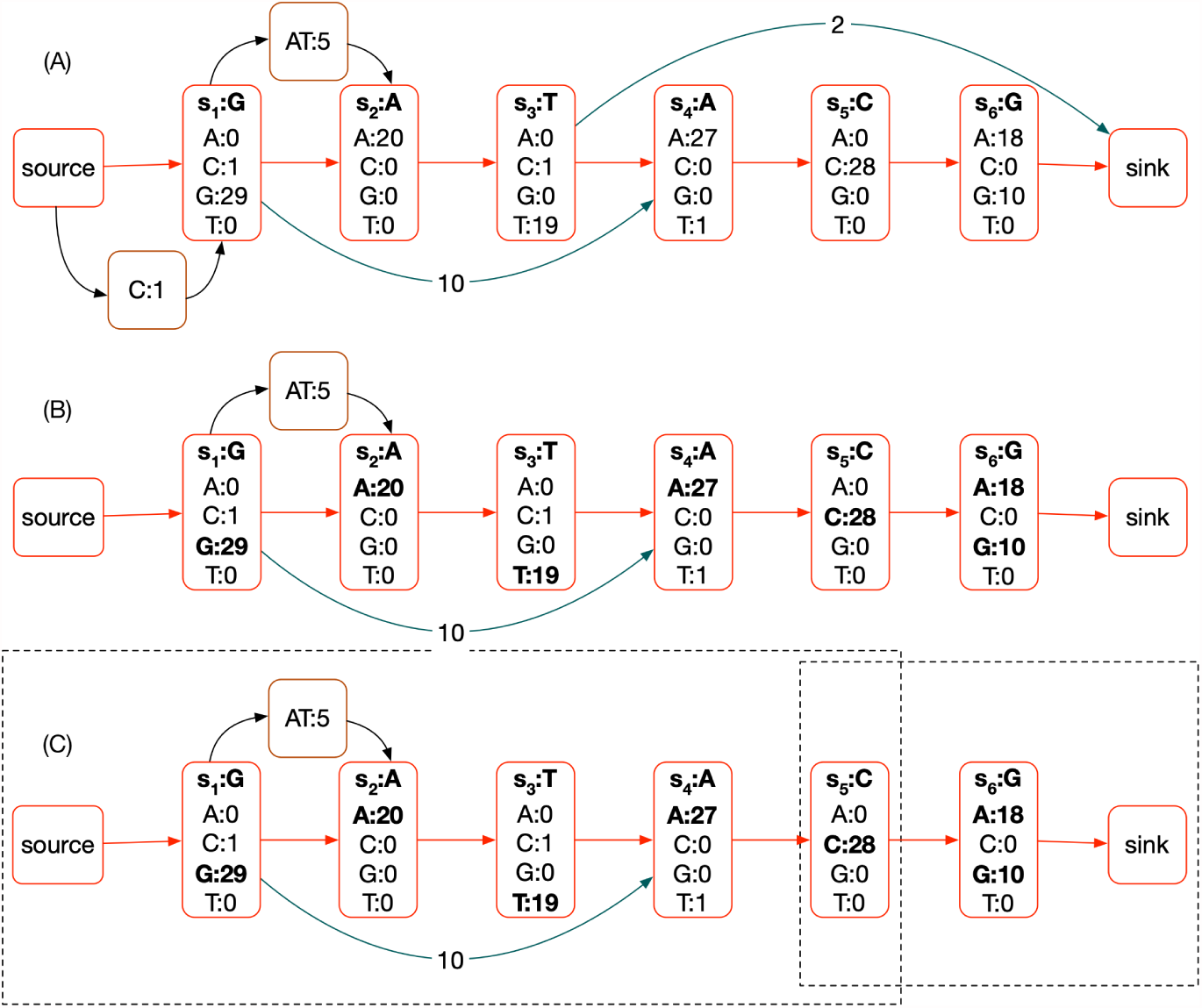
A) An example POA, assuming approximately 30x read coverage. The backbone is shown in red. Each non-source/sink node has a vector of weights, one for each possible base. Deletion edges are shown in teal, they also each have a weight. Finally insertion nodes are shown in brown, each also has a weight. (B) A pruned POA, removing deletions and insertions that have less than a threshold weight and highlighting plausible bases in bold. There are six plausible nucleotide sequences represented by paths through the POA and selections of plausible base labels: G;AT;A;T;A;C:A, G;AT;A;T;A;C:G, G;A;T;A;C:A, G;A;T;A;C:G, G;A;C:A, G;A;C:G. To avoid the combinatorial explosion of such enumeration we identify subgraphs (C) and locally enumerate the possible subsequences in these regions independently (dotted rectangles identify subgraphs selected). In each subgraph there is a source and sink node that does not overlap any proposed edit.

Frequently either an insertion or deletion can be made between different successive bases in *s* resulting in the same edited sequence. To ensure that such equivalent events are not represented multiple times in the POA, and to ensure we sum their weights correctly, we ‘left shift’ indels to their maximum extent. When shifting an indel results in multiple equivalent deletion edges or insertions we remove the duplicate elements, updating the weight of the residual element to include the sum of the weights of the removed elements. For example, the insertion of ‘AT’ in Fig. 14 is shifted left to its maximal extent, and could include the merger of an equivalent ‘AT’ insertion starting two backbone nodes to the right.

### Local Haplotype Proposal

After constructing the POA we use it to sample alternative assemblies. We first prune the POA to mark indels and base substitutions with weight below a threshold, which are generally the result of sequencing errors (Fig. 14(B)). Currently this threshold (`polish.candidateVariantWeight=0.18`, established empirically) is normalized as a fraction of the estimated coverage at the site, which is calculated in a running window around each node in the backbone of 100 bases. Consequently if fewer than 18% of the reads are expected to include the change then the edit is pruned from consideration.

To further avoid a combinatorial explosion we sample alternative assemblies locally. We identify subgraphs of *s* containing indels and substitutions to *s* then in each subgraph, defined by a start and end backbone vertex, we enumerate all possible paths between the start and end vertex and all plausible base substitutions from the backbone sequence. The rationale for heuristically doing this locally is that two subgraphs separated by one or more *anchor* backbone sites with no plausible edits are conditionally independent of each other given the corresponding interstitial anchoring substring of *s* and the substrings of the reads aligning to it. Currently, any backbone site more than `polish.columnAnchorTrim=5` nodes (equivalent to bases) in the backbone from a node overlapping a plausible edit (either substitution or indel) is considered an anchor. This heuristic allows for some exploration of alignment uncertainty around a potential edit. Given the set of anchors computation proceeds by identifying successive pairs of anchors separated by subgraphs containing the potential edits, with the two anchors considered the source and sink vertex.

### A Simple Bayesian Model for Run-length Decoding

Run-length encoding allows for separate modelling of length and nucleotide error profiles. In particular, length predictions are notoriously error prone in nanopore basecalling. Since homopolymers produce continuous signals, and DNA translocates at a variable rate through the pore, the basecaller often fails to infer the true number of bases given a single sample. For this reason, a Bayesian model is used for error correction in the length domain, given a distribution of repeated samples at a locus.

To model the error profile, a suitable reference sequence is selected as the truth set. Reads and reference are run-length encoded and aligned by their nucleotides. The alignment is used to generate a mapping of observed lengths to their true length (*y, x*) where *y* = *true* and *x* = *observed* for each position in the alignment. Observations from alignment are tracked using a matrix of predefined size (*y*_*max*_ = 50*, x*_*max*_ = 50) in which each coordinate contains the corresponding count for (*y, x*). Finally the matrix is normalized along one axis to generate a probability distribution of *P* (*X|y*_*j*_) for *j* in [1, *y*_*max*_]. This process is performed for each of the 4 bases.

With enough observations, the model can be used to find the most probable true run length given a vector of observed lengths *X*. This is done using a simple log likelihood calculation over the observations *x*_*i*_ for all possible true lengths *y*_*j*_ in *Y*, assuming the length observations to be independent and identically distributed. The length *y*_*j*_ corresponding to the greatest likelihood *P* (*X|y*_*j*_, *Base*) is chosen as the consensus length for each alignment position (Fig. 15).

**Figure 15:**
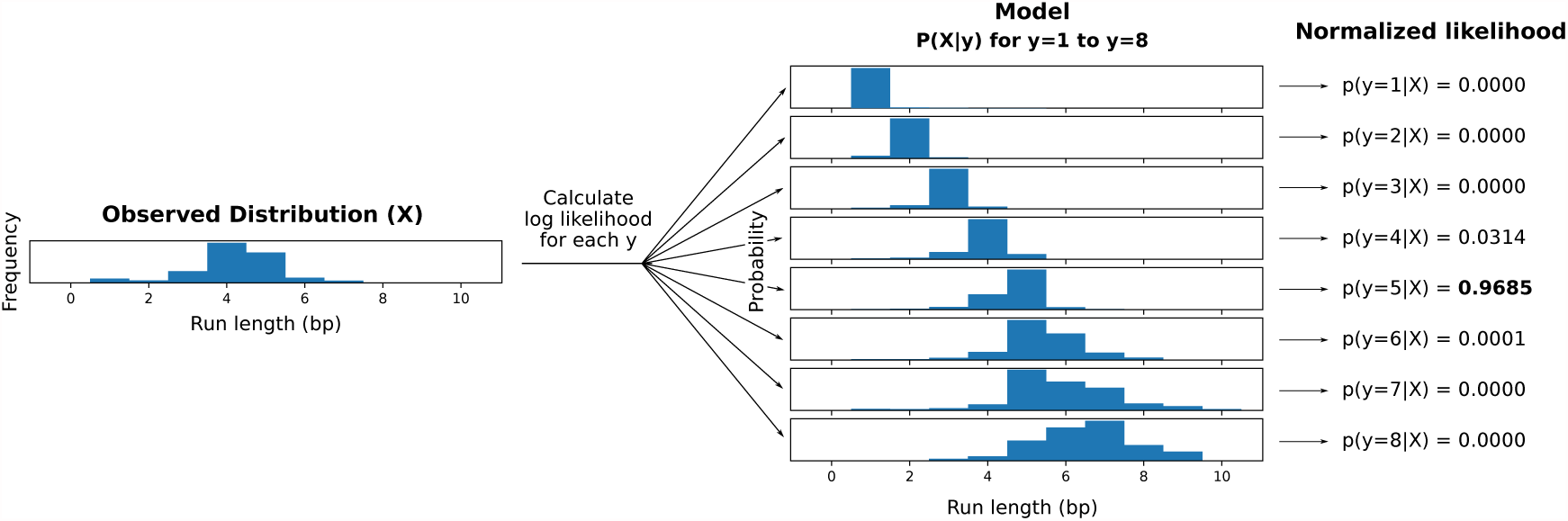
Visual representation of run length inference. This diagram shows how a consensus run length is inferred for a set of aligned lengths (X) that pertain to a single position. The lengths are factored and then iterated over, and log likelihood is calculated for every possible true length up to a predefined limit. Note that in this example, the most frequent observation (4bp) is not the most likely true length (5bp) given the model.

### Training

To generate a model, we ran MarginPolish with reads from a specific basecaller version aligned to a reference (GRCh38) and specified the `–outputRepeatCounts` flag. This option produces a TSV for each chunk describing all the observed repeat counts aligned to each backbone node in the POA. These files are consumed by a script in the https://github.com/rlorigro/runlength_analysis repository, which generates a RLE consensus sequence, aligns to the reference, and performs the described process to produce the model.

The `allParams.np.human.guppy-ff-235.json` model used for most of the analysis was generated from HG00733 reads basecalled with Guppy Flipflop v2.3.5 aligned to GRCh38, with chromosomes 1, 2, 3, 4, 5, 6, and 12 selected. The model `allParams.np.human.guppy-ff-233.json` was generated from Guppy Flipflop v2.3.3 data and chromosomes 1-10 were used. This model was also used for the CHM13 analysis, as the run-length error profile is very similar between v2.3.3 and v2.3.1 (v2.3.5 has a drastically different error profile, as is shown below in Fig. 18).

### Parallelization and Computational Considerations

To parallelize MarginPolish we break the assembly up into chunks of size `polish.chunkSize`=1000 bases, with an overlap of `polish.chunkBoundary`=50 bases. We then run the MarginPolish algorithm on each chunk independently and in parallel, stitching together the resulting chunks after finding an optimal pairwise alignment (using the default hmm described earlier) of the overlaps that we use to remove the duplication. We can further parallelize the algorithm across machines or processes using a provided Toil script CITE:PMID: 28398314.

Memory usage scales with thread count, read depth, and chunk size. For this reason, we downsample reads in a chunk to `polish.maxDepth=50`× coverage by counting total nucleotides in the chunk *N*_*c*_ and discarding reads with likelihood 1 −(`chunkSize` + 2 ∗ `chunkBoundary`) ∗ `maxDepth`*/N*_*c*_. With these parameters, we find that 2GB of memory per thread is sufficient to run MarginPolish on genome-scale assemblies. Across 13 whole-genome runs, we averaged roughly 350 CPU hours per gigabase of assembled sequence.

## HELEN: Homopolymer Encoded Long-read Error-corrector for Nanopore

HELEN is a deep neural network based haploid consensus sequence polisher. HELEN employs a multi-task recurrent neural network (RNN) [39] that takes the weights of the partial order alignment (POA) graph of MarginPolish to predict a base and a run-length for each genomic position. MarginPolish constructs the POA graph by performing multiple possible alignments of a single read that makes the weights associative to the correct underlying base and a run-length. The RNN employed in HELEN takes advantage of the transitive relationship of the genomic sequence and associative coupling of the POA weights to the correct base and run-length to produce a consensus sequence with higher accuracy.

The error-correction with HELEN is done in three steps. First, we generate tensor-like images of genomic segments with MarginPolish that encodes POA graph weights for each genomic position. Then we use a trained RNN model to produce predicted bases and run-lengths for each of the generated images. Finally, we stitch the chunked sequences to get a contiguous polished sequence.

### Image Generation

MarginPolish produces an image-like summary of the final POA state for use by HELEN. At a high level, the image summarizes the weighted alignment likelihoods of all reads divided into nucleotide, orientation, and run-length.

The positions of the POA nodes are recorded using three coordinates: the position in the backbone sequence of the POA, the position in the insert sequences between backbone nodes, and the index of the run-length block. All backbone positions have an insert coordinate of 0. Each backbone and insert coordinate includes one or more run-length coordinate.

When encoding a run-length, we divide all read observations into blocks from 0 to 10 inclusive (this length is configurable). For cases where no observations exceed the maximum run-length, a single run-length image can describe the POA node. When an observed run-length exceeds the length of the block, the run-length is encoded as that block’s maximum (10), and the remaining run-length is encoded in successive blocks. For a run-length that terminates in a block, its weight is contributed to the run-length 0 column in all successive blocks. This means that the records for all run-length blocks of a given backbone and insert position have the same total weight. As an example, consider three read positions aligned to a node with run-lengths of 8, 10, and 12. These require two run-length blocks to describe: the first block includes one 8 and two 10s, and the second includes two 0s and one 2.

The information described at each position (backbone, insert, and run-length) is encoded in 92 features: each nucleotide {A, C, T, G} and run-length {0, 1,.., 10}, plus a gap weight (for deletions in read alignments). The weights for each of these 45 observations are separated into forward and reverse strand for a total of 90 features. The weights for each of these features are normalized over the total weight for the record and accompanied by an additional data point describing the total weight of the record. This normalization column for the record is an approximation of the read depth aligned to that node. Insert nodes are annotated with a binary feature (for a final total of 92); weights for an insert node’s alignments are normalized over total weight at the backbone node it is rooted at (not the weight of the insert node itself) and gap alignment weights are not applied to them.

Labeling nodes for training requires a truth sequence aligned to the assembly reference. This provides a genome-scale location for the true sequence and allows the its length to help in the resolution of segmental duplications or repetitive regions. When a region of the assembly is analyzed with MarginPolish, the truth sequences aligned to that region are extracted. If there is not a single truth sequence which approximately matches the length of the consensus for this region, we treat it as an uncertain region and no training images are produced. Having identified a suitable truth sequence, it is aligned to the final consensus sequence in non-run-length space with Smith-Waterman. Both sequences and the alignment are then run-length encoded, and true labels are matched with locations in the images. All data between the first and last matched nodes are used in the final training images (leading and trailing inserts or deletes are discarded). For our training, we aligned the truth sequences with `minimap2` using the `asm20` preset and filtered the alignments to include only primary and supplementary alignments (no secondary alignments).

Fig. 16 shows a graphical representation of the images. On the y-axis we display true nucleotide labels (with the dash representing no alignment / gap) and true run-length. On the x-axis the features used as input to HELEN are displayed: first the normalization column (the total weight at the backbone position), second the insert column (the binary feature encoding whether the image is for a backbone or insert node), forty-eight columns describing the weights associated with read observations (stratified by nucleotide, run-length, strand), and two columns describing weights for gaps in read alignments (stratified by strand). In this example, we have reduced the maximum run-length per block from 10 to 5 for demonstrative purposes.

**Figure 16:**
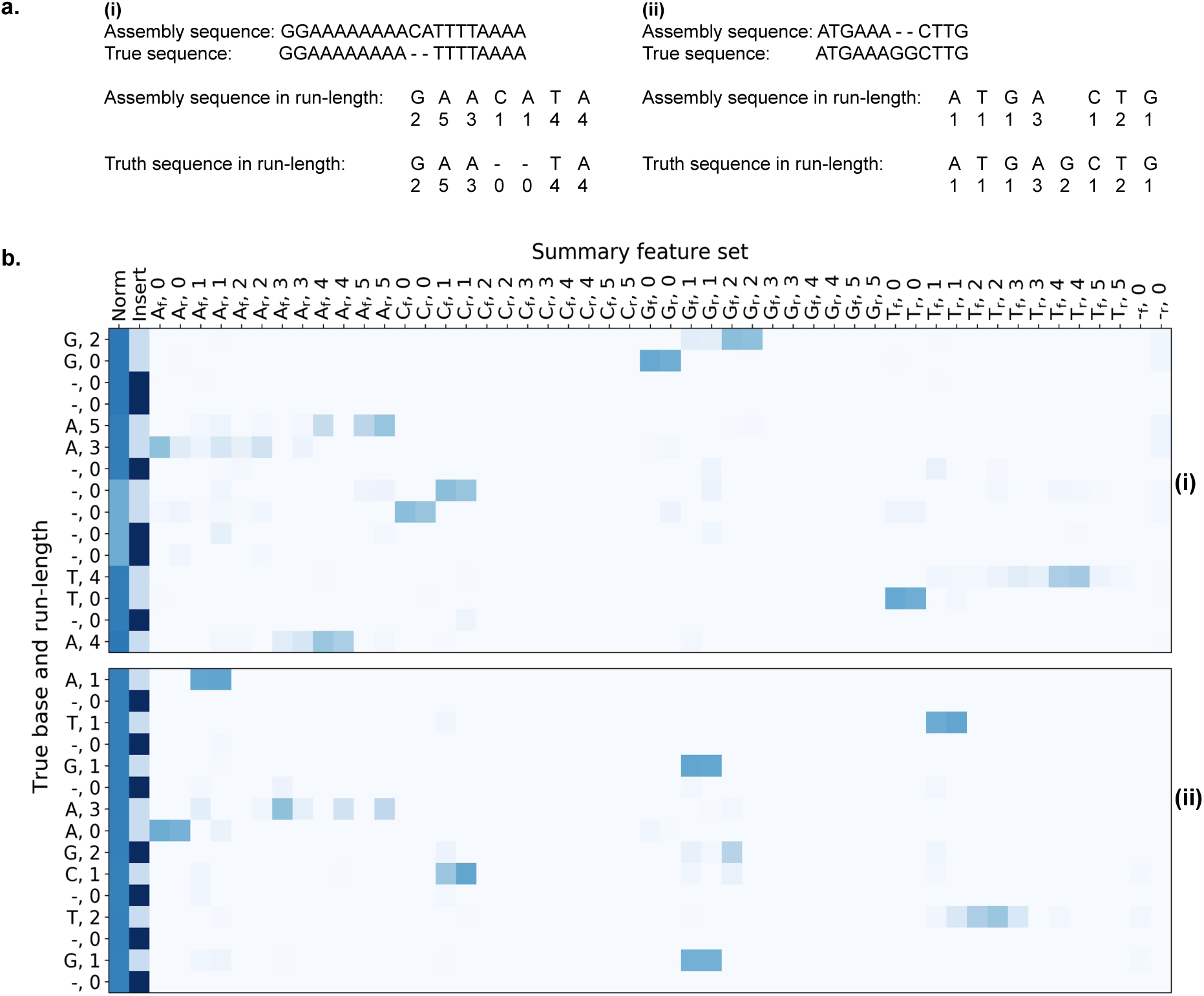
MarginPolish Images. A graphical representation of images from two labeled regions selected to demonstrate: the encoding of a single POA node into two run-length blocks (i), a true deletion (i), and a true insert (ii). The y-axis shows truth labels for nucleotides and run-lengths, the x-axis describes features in the images, and colors show associated weights.

We selected these two images to highlight three features of the model: the way multiple run-length blocks are used to encode observations for a single node, and the relevant features around a true gap and a true insert that enable HELEN to correct these errors.

To illustrate multiple run length blocks, we highlight two locations on on image (i). The first are the nodes labeled (A,5) and (A,3). This is the labeling for a true (A,8) sequence separated into two blocks. See that the bulk of the weight is on the (A,5) features on the first block, with most of that distributed across the (A,1-3) features on the second. Second, observe the nodes on (i) labeled (T,4) and (T,0). Here we show the true labeling of a (T,4) sequence where there are some read observations extending into a second run-length block.

To show a features of a true gap, note on (i) the non-insert nodes labeled (-,0). We know that MarginPolish predicted a single cytosine nucleotide (as it is a backbone node and the (C,1) nodes have the bulk of the weight. Here, HELEN is able to use the low overall weight (the lighter region in the normalization column) at this location as evidence of fewer supporting read alignments and can correct the call.

The position labeled (G,2) on (ii) details a true insertion. It is not detected by MarginPolish (as all insert nodes are not included in the final consensus sequence). Read support is present for the insert, less than the backbone nodes in this image but more than the other insert nodes. HELEN can identify this sequence and correct it.

**Figure 17:**
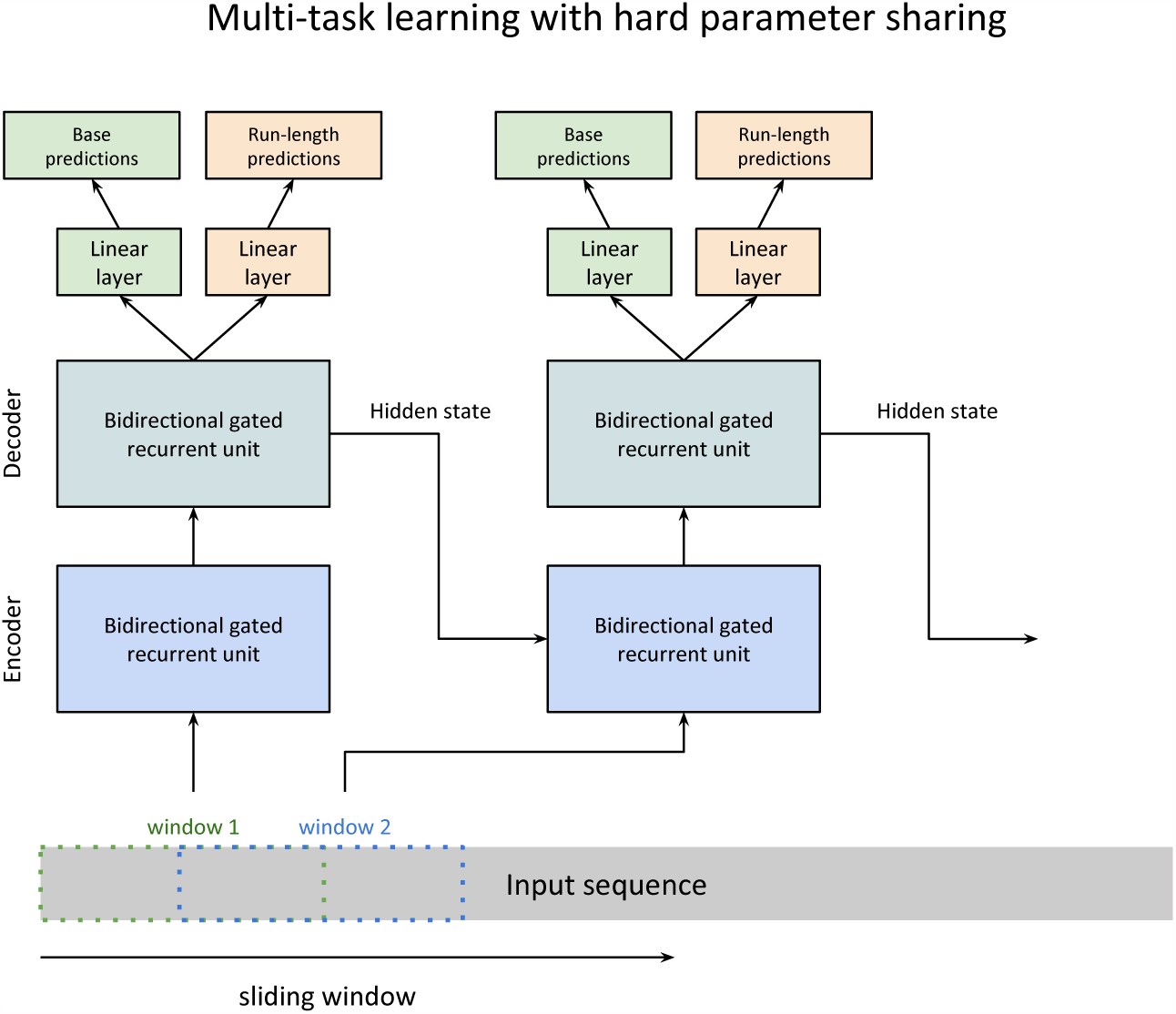
The sequence-to-sequence model implemented in Helen.

### The model

We use a sequence transduction model for consensus polishing. The model structure consists of two single-layer gated recurrent neural units (GRU) for encoding and decoding on top of two linear transformation layers. The two linear transformation layers independently predict a base and a run-length for each position in the input sequence. Each unit of the GRU can be described using the four functions it calculates:

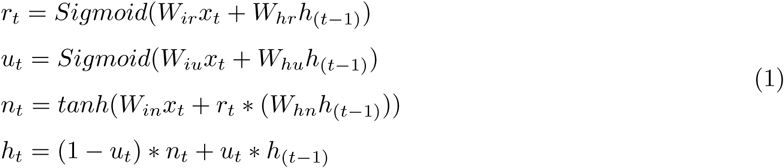

For each genomic position *t*, we calculate the current state *h*_*t*_ from the new state *n*_*t*_ and the update value *u*_*t*_ applied to the output state of previous genomic position *h*_(t−1)_. The update function *u*_*t*_ decides how much past information to propagate to the next genomic position. It multiplies the input *x*_*t*_ with the weight vector *W*_*iu*_ and multiplies the hidden state of the previous genomic position *h*_(*t*−1)_. The weight vectors decide how much from the previous state to propagate to the next state. The reset function *r*_*t*_ decides how much information to dissolve from the previous state. Using a different weight vector, the *r*_*t*_ function decides how much information to dissolve from the past. The new memory state *n*_*t*_ is calculated by multiplying the input *x*_*t*_ with the weight vector *W*_*in*_ and applying a Hadamard multiplication ∗ between the reset function value and a weighted state of the previous hidden state *h*_(*t*−1)_. The new state captures the associative relationship between the input weights and true prediction. In this setup, we can see that *r*_*t*_ and *u*_*t*_ can decide to hold memory from distant locations while *n*_*t*_ captures the associative nature of the weights to the prediction, helping the model to decide how to propagate genomic information in the sequence. The output of each genomic position *h*_*t*_ can be then fed to the next genomic position as a reference to the previously decoded genomic position. The final two layers apply linear transformation functions:

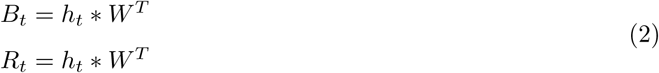

The two linear transformation functions independently calculate a base prediction *B*_*t*_ and a run-length prediction *R*_*t*_ from the hidden state output of that genomic position *h*_*t*_. The model operates in hard parameter sharing mode where the model learns to perform two tasks in equation 2 using the same set of underlying parameters from equation 1. The ability of the model to reduce the error rate of the assemblies from multiple samples with multiple assemblers shows the generalizability and robustness we achieve with this method.

### Sliding window mechanism

One of the challenges of this setup is the sequence length. From the functions of recurrent units in equation 1, we see that each state is updated based on the previous state and associated weight. Due to the noisy nature of the data, if the sequence length is too long, the back-propagation becomes difficult over noisy regions. On the other hand, a small sequence length would make the program very slow. We balance the run-time and accuracy by using a sliding window approach.

During the sliding-window, we chunk the sequence of thousand bases to multiple overlapping windows of length 100. Starting from the leftmost window, we perform prediction on sequence pileups of the window and transmit the hidden state of the current window to the next window and slide the window by 50 bases to the right. For each window, we collect all the predicted values and add it to a global sequence inference counter that can keep track of predicted probabilities of base and run-length at each position. Lastly, we aggregate the probabilities from the global inference counter to generate a sequence. This setup allows us to utilize the minibatch feature of the popular neural network libraries allowing inference on a batch of inputs instead of performing inference one at a time.

### Training the model

HELEN is trained with a gradient descent method. We use Adaptive Moment Estimation (Adam) method to compute gradients for each of the parameters in the model based on a target loss function. Adam uses both decaying squared gradients and the decaying average of gradients, making it suitable to use with recurrent neural networks[39]. Adam performs gradient optimization by adapting the parameters to set in a way that minimizes the value of the loss function.

We perform optimization through back-propagation per window of the input sequence. From equation 2, we see that we get two vectors *B* = [*B*_1_, *B*_2_, *B*_3_…*B*_*n*_] and *R* = [*R*_1_, *R*_2_, *R*_3_*…R*_*n*_] containing base and run-length predictions for each window of size *n*. From the labeled data we get two more such vectors *T*_*B*_ = [*T*_*B*1_, *T*_*B*2_, *T*_*B*3_, …*T*_*Bn*_] and *T*_*R*_ = [*T*_*R*1_, *T*_*R*2_, *T*_*R*3_, …*T*_*Rn*_] containing the true base and true rle values of each position in the window. From these loss function the loss *L* is calculated:

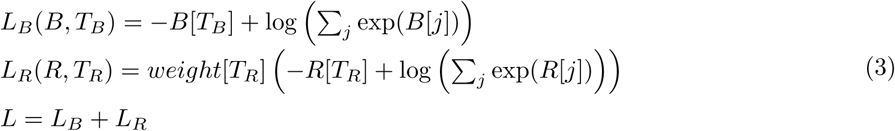

In equation 3, *L*_*B*_ calculates the base prediction loss and *L*_*R*_ calculates the rle prediction loss. The rle class distribution is heavily biased toward lower run-length values, so, we apply class-wise weights depending on the observation of per class to make the learning process balanced between classes. The optimizer then updates the parameters or weights *W* of the model from equation 1 and equation 2 in a way that minimizes the value of the loss function. We can see that the loss function is a summation of the two independent loss functions but the underlying weights from the recurrent neural network belongs to the same set of elements in the model. In this setting, the model optimizes to learn both task simultaneously by updating the same set of weights.

### Sequence stitching

To parallelize the polishing pipeline, MarginPolish chunks the genome into smaller segments while generating images. Each image segment encodes a thousand nucleotide bases, and two adjacent chunks have 50 nucleotide bases overlap between them. During the inference step, we save all run-length and base predictions of the images, including their start and end genomic positions.

For stitching, we load all the image predictions and sort them based on the genomic start position of the image chunk and stitch them in parallel processes. For example, if there are *n* predictions from *n* images of a contig and we have *t* available threads, we divide *n* prediction chunks into *t* buckets each containing approximately *n/t* predicted sequences. Then we start *t* processes in parallel where each process stitches all the sequences assigned to it and returns a longer sequence. For stitching two adjacent sequences, we take the overlapping sequences between the two sequence and perform a pairwise Smith-Waterman alignment. From the alignment, we pick an anchor position where both sequences agree the most and create one sequence. After all the processes finish stitching the buckets, we get *t* longer sequences generated by each process. Finally, we iteratively stitch the *t* sequences using the same process and get one contiguous sequence for the contig.

### Generating trained models

In supplementary tables 13, 16 and 15 we report several models for HELEN. The models are trained on different sets of data with varying Guppy base-caller versions. We discuss three trained models `r941_flip235_v001.pkl, r941_flip233_v001.pkl, and r941_flip231_v001.pkl` to use with HELEN for different versions of the ONT Guppy base-callers. Due to the difference in the error profile of different versions of the Guppy base-caller, we trained three different models.

**Table 3:** Description of trained models for HELEN.

| Model Name | Base caller version | Training sample | Training region | Testing region |
| --- | --- | --- | --- | --- |
| `r941_flip235_v001.pkl` | Guppy 2.3.5 | HG002 | Chr1-19, Chr21-22 | Chr20 |
| `r941_flip233_v001.pkl` | Guppy 2.3.3 | HG002 | Chr1-19, Chr21-22 | Chr20 |
| `r941_flip231_v001.pkl` | Guppy 2.3.1 | CHM13 | Chr1-6 | Chr20 |

The `r941_flip235_v001.pkl` is trained on HG002 base called with Guppy 2.3.5. The model is trained on the high confidence regions of all autosomes and tested on Chr20. The training script trained the model for 80 hours on 10 epochs, which generated 10 trained models. We picked the model that has the best performance on Chr20 as the final model.

The CHM13 data from T2T consortium [31] were base called with Guppy 2.3.1. The error profile of Guppy 2.3.1 is significantly different than Guppy 2.3.5. Figure 18 shows the difference in underlying error profile of HG00733 sample for two different versions of Guppy. We trained `r941_flip233_v001.pkl` Model on HG002 Guppy 2.3.3 data. Although the error profile of Guppy 2.3.1 and Guppy 2.3.3 are similar, the reported base qualities are different. So, we trained another model `r941_flip231_v001.pkl` on Chr1-6 of CHM13 to see further improvement in the consensus quality of CHM13.

**Figure 18:**
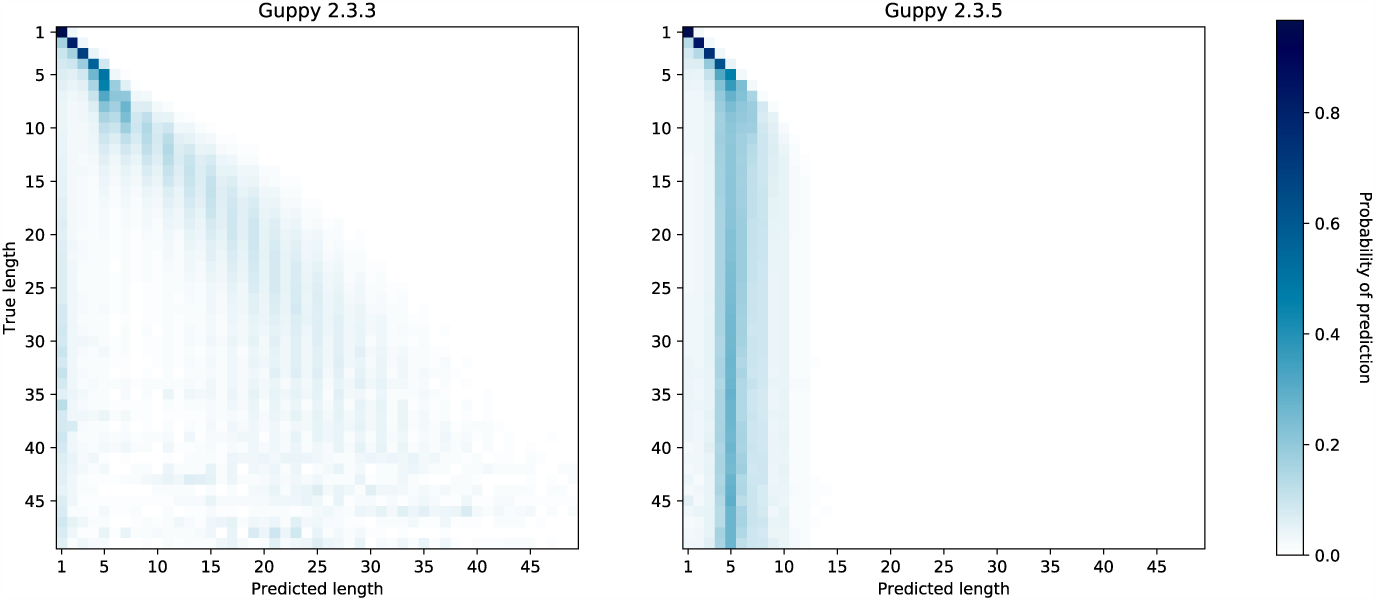
Run-length confusion in different versions of Guppy base caller

### Implementation notes

We have implemented HELEN using python and C++ programming language. We use PyTorch [69] deep neural network library for the model implementation. We also use the Striped-Smith Waterman algorithm implementation to use during stitching and Pybind11 [70] as a bridge between C++ and python methods. The image data is saved using HDF5 file format. The implementation is publicly available via GitHub (https://github.com/kishwarshafin/helen).

