## Supplement for "Efficient *de novo* assembly of eleven human genomes using PromethION sequencing and a novel nanopore toolkit"

A Preprint

### Supplementary Notes

#### Execution Parameters

#### Shasta

All Shasta runs used Shasta version 0.1.0 built from `https://github.com/chanzuckerberg/shasta`. Rather than using the distributed version of the release, the source code was rebuilt locally for best performance as recommended by Shasta documentation.

The Shasta executable was run with the following command:

```
shasta \
--memoryMode filesystem \
--memoryBacking 2M
```

#### Canu

Canu 1.8 from `https://github.com/marbl/canu` was run with the following command:

```
canu \
-p asm \
-d asm \
genomeSize=3.1g \
'corMhapOptions=--threshold 0.8 --num-hashes 512
                --ordered-sketch-size 1000 --ordered-kmer-size 14' \
'gridOptionsJobName=mom' \
'gridOptions=--time=240:00:00 --partition=norm' \
'stageDirectory=/lscratch/$SLURM_JOBID' \
'gridEngineStageOption=--gres=lscratch:100' \
'correctedErrorRate=0.105' \
-nanopore-raw input.fastq.gz
```

#### Wtdbg2

Wtdbg2 version 2.3 from `https://github.com/ruanjue/wtdbg2` was run with the following commands:

```
wtdbg2 \
-t 0 \
-x ont \
-L 10000 \
-g 3.3g \
-i reads1.fastq.gz \
```

```
-i reads2.fastq.gz \
-i reads3.fastq.gz \
-o wtdbg2-assembly

wtpoa-cns \
-t 31 \
-i wtdbg2-assembly.ctg.lay.gz \
-f \
-o wtdbg2-assembly.fa
```

#### Flye

Flye version 2.4.2 from `https://github.com/fenderglass/Flye` was run with the following command:

```
flye \
--nano-raw reads1.10kb.fastq.gz reads2.10kb.fastq.gz reads3.10kb.fastq.gz
--genome-size 3.3g \
--out-dir flye \
--threads 123
```

#### Racon

We used a home-grown script to manage running 4 iterations of racon. The code can be found here `https://github.com/rlorigro/nanopore_assembly_and_polishing_assessment`, and was run with the following command:

```
python3 /home/ubuntu/software/nanopore_assembly_and_polishing_assessment/polish.py \
--true_ref hg38.fa \
--contigs assembly.fasta \
--sequences reads.fasta \
--output_dir racon \
--n_passes 4
```

#### Medaka

Medaka version 0.6.0-alpha.3 from `https://github.com/nanoporetech/medaka` was run with the following commands:

```
medaka consensus \
-i reads5.fasta \
-d assembly_racon4x.fasta \
-o medaka \
-t 64 \
-m r941_flip235

medaka stitch \
medaka/consensus_probs.hdf \
medaka/consensus.fasta
```

#### Minialign

Minialign is bundled with Medaka, and was run with the following commands:

```
mini_align \
-i reads.fasta \
-r assembly.fasta \
-P \
-m \
-p medaka/calls_to_draft \
-t 60
```

#### Minimap2, Samtools

Minimap2 version 2.15-r908-dirty from https://github.com/lh3/minimap2. We used samtools 1.7 using htslib 1.7-2 for sorting and filtering. The following three commands were piped into each other:

```
minimap2 \
-ax map-ont \
-t 70 \
assembly.fasta \
reads.fasta

samtools sort \
-@ 70

samtools view \
-hb \
-F 0x104 \
>align.bam
```

#### MarginPolish

MarginPolish 1.0.0 was from https://github.com/UCSC-nanopore-cgl/MarginPolish run with the following command:

```
marginPolish \
input.bam \
input.fa \
allParams.np.human.guppy-ff-235.json \
-f \
-o output\_location \
-t 70
```

#### HELEN

HELEN version 0.1 from https://github.com/kishwarshafin/helen was run with the following commands:

```
python3 /home/ubuntu/software/helen/call_consensus.py \
-i images/ \
-b 1024 \
-w 16 \
-t 32 \
-m r941_flip235_v001.pkl \
-o out \
-g

python3 /home/ubuntu/software/helen/stitch.py \
-i out/helen_predictions_05312019_183902.hdf \
-o out/ \
-p polished_assembly \
-t 32
```

#### HiRise

HiRise was run via a docker container, with access given by Dovetail Genomics. The HiRise version was v2.1.6, with the HiRise Helper version 2.1.10 and the HiRise Utils version v2.1.7-3-g98c1a1b. Default parameters were used.

#### Trio-binning

For HG00733, the parental read sample accessions were obtained from 1000 genome database:

http://www.internationalgenome.org/data-portal/sample/HG00731
http://www.internationalgenome.org/data-portal/sample/HG00732

Briefly, k-mers were counted with `meryl`, subtracted to generate maternal/paternal sets, and any k-mers occurring less than 6 times for maternal k-mers and 5 times for paternal k-mers were not used. Binning did not use normalization by k-mer set size. This resulted in 35.2x maternal, 37.3x paternal, and 5.6x unclassified. Assembly did not use the unclassified reads and ran with the command:

```
canu \
-p asm \
-d <mom/dad>
'genomeSize=3.1g' \
'corMhapOptions=--threshold 0.8 --num-hashes 512
                --ordered-sketch-size 1000 --ordered-kmer-size 14' \
'corMinCoverage=0'
```

Each haplotype assembly required approximately 100k CPU hours (4-5 days). A subsequent run using Canu 1.8 and automated binning with the command:

```
canu \
-p asm \
-d asm \
'genomeSize=3.1g' \
'corMhapOptions=--threshold 0.8 --num-hashes 512
                --ordered-sketch-size 1000 --ordered-kmer-size 14' \
'gridOptionsJobName=733_trio' \
'corMinCoverage=0' \
-haplotypeMOM hg0732/*fastq.gz \
-haplotypeDAD hg0731/*.fastq.gz
```

resulted in a similar classification split (35.1x dad, 36.7x mom, 5.6x unknown) and assembly (manual: dad=16.6 NG50, mom=18.1 NG50; automated: dad=14.1 NG50, mom=19.9 NG50).

For HG0002, illumina data for the parents was downloaded from the GIAB ftp site:

```
ftp://ftp-trace.ncbi.nlm.nih.gov/giab/ftp/data/AshkenazimTrio/HG003_NA24149_father \
    /NIST_HiSeq_HG003_Homogeneity-12389378/HG003_HiSeq300x_fastq/
ftp://ftp-trace.ncbi.nlm.nih.gov/giab/ftp/data/AshkenazimTrio/HG004_NA24143_mother \
    /NIST_HiSeq_HG004_Homogeneity-14572558/HG004_HiSeq300x_fastq/
```

K-mers were counted as before, subtracted, and filtered to exclude k-mers occuring less than 25 times in the maternal or paternal set. The classification resulted in 24x maternal, 23x paternal, and 3.5x unknown. Only classified reads were used for assembly with the command:

```
canu \
-p asm \
-d <mom/dad> \
'genomeSize=3.1g' \
'corMhapOptions=--threshold 0.8 --num-hashes 512
                --ordered-sketch-size 1000 --ordered-kmer-size 14'  \
'corMinCoverage=0'
```

Each haplotype assembly required approximately 100k cpu hours (4-5 days).

#### Benchmarking assemblies using Pomoxis

The truth assembly files and the reported error-rates are described in Online methods.

To benchmark the assemblies, we used `assess_assembly` version 0.2.2 from Pomoxis. This tool is developed and suggested by the research group of Oxford Nanopore Technology. The installation instruction of Pomoxis can be found on the github page https://github.com/nanoporetech/pomoxis. The parameters we used are:

- -i: The input assembly (fasta).
- -r: The reference fasta file. (The truth assembly)

- -b: Bed file containing reference regions to assess.
- -p: Prefix of the output file names.
- -t: Number of threads to use.
- -T: Trim consensus to primary alignments of truth to assembly.

We compared the HG002 samples, we gathered the truth assembly `hg002_truth_assembly.fa`, a bed file `hg002_confident.bed` describing the confident regions and a shasta assembly `hg002_shasta_assembly.fa` and ran the following command.

```
assess_assembly \
-i hg002_shasta_assembly.fa \
-r hg002_truth_assembly.fa \
-b hg002_confident.bed \
-p hg002_shasta_assessment \
-t 32 \
-T
```

In this setup, the `assess_assembly` module computes the error rate of the input `hg002_shasta_assembly.fa` that aligns to the high-confidence region defined in the `hg002_confident.bed` of `hg002_truth_assembly.fa` assembly. Also, the `-T` parameter limits the assessment to regions where there is an alignment between the truth and the input assembly.

For HG00733 sample, we used the high-quality phased PacBio assembly. We got `hg00733_truth_assembly.fa` and the `hg00733_shasta_assembly.fa` and ran the following command for assessment.

```
assess_assembly \
-i hg00733_shasta_assembly.fa \
-r hg00733_truth_assembly.fa \
-p hg00733_shasta_assessment \
-t 32 \
-T
```

As the truth assembly of HG00733 does not define any high-confidence region, we do a whole genome comparison where there is an alignment between the truth and the input assembly enforced by the `-T` parameter. For CHM13 and all other assemblies, we used the same command as HG00733. The output of this program reports different error rates described in the online methods section.

### Supplementary Results

#### Nanopore sequencing eleven human genomes in nine days

Supplementary Table 1: Read N50s stratified by sample and flowcell (three for each sample) for 11 samples.

| Sample | Flowcell No. | Flowcell N50 | Sample N50 |
|---|---|---|---|
| GM24143 | 1 | 48891 | 46757 |
| | 2 | 47044 | |
| | 3 | 44335 | |
| GM24149 | 1 | 46054 | 43306 |
| | 2 | 44245 | |
| | 3 | 39618 | |
| GM24385 | 1 | 50349 | 48705 |
| | 2 | 49319 | |
| | 3 | 46448 | |
| HG00733 | 1 | 29862 | 29584 |
| | 2 | 30473 | |
| | 3 | 28417 | |
| HG01109 | 1 | 48795 | 45894 |
| | 2 | 44218 | |
| | 3 | 44670 | |
| HG01243 | 1 | 45467 | 43567 |
| | 2 | 44681 | |
| | 3 | 40554 | |
| HG02055 | 1 | 44320 | 45457 |
| | 2 | 47148 | |
| | 3 | 44902 | |
| HG02080 | 1 | 38519 | 39319 |
| | 2 | 40123 | |
| | 3 | 39315 | |
| HG02723 | 1 | 50509 | 49723 |
| | 2 | 47842 | |
| | 3 | 50817 | |
| HG03098 | 1 | 41463 | 40629 |
| | 2 | 42308 | |
| | 3 | 38115 | |
| HG03492 | 1 | 32149 | 30168 |
| | 2 | 30063 | |
| | 3 | 28292 | |
| **Average** | **-** | **41889** | **42101** |

Supplementary Table 2: Throughput stratified by sample and flowcell (three for each sample) in gigabases (Gb) for 11 samples.

| Sample | Flowcell No. | Flowcell (Gb) | Sample (Gb) | Coverage |
|---|---|---|---|---|
| GM24143 | 1 | 87 | 280 | 84.72 |
| | 2 | 97 | | |
| | 3 | 95 | | |
| GM24149 | 1 | 82 | 273 | 82.6 |
| | 2 | 107 | | |
| | 3 | 84 | | |
| GM24385 | 1 | 26 | 157 | 47.43 |
| | 2 | 71 | | |
| | 3 | 59 | | |
| HG00733 | 1 | 62 | 242 | 73.45 |
| | 2 | 90 | | |
| | 3 | 89 | | |
| HG01109 | 1 | 71 | 219 | 66.48 |
| | 2 | 79 | | |
| | 3 | 70 | | |
| HG01243 | 1 | 71 | 187 | 56.68 |
| | 2 | 73 | | |
| | 3 | 43 | | |
| HG02055 | 1 | 71 | 202 | 61.33 |
| | 2 | 67 | | |
| | 3 | 65 | | |
| HG02080 | 1 | 71 | 172 | 52.21 |
| | 2 | 42 | | |
| | 3 | 59 | | |
| HG02723 | 1 | 81 | 227 | 68.7 |
| | 2 | 69 | | |
| | 3 | 78 | | |
| HG03098 | 1 | 79 | 177 | 53.63 |
| | 2 | 40 | | |
| | 3 | 58 | | |
| HG03492 | 1 | 61 | 158 | 47.74 |
| | 2 | 45 | | |
| | 3 | 51 | | |
| **Average** | **-** | **69** | **208** | **63.18** |

Supplementary Table 3: Mean, median, and modal values for read alignment identities of 11 samples, aligned to GRCh38.

| Sample | Mean | Median | Mode |
|---|---|---|---|
| GM24143 | 0.87188 | 0.89651 | 0.920 |
| GM24149 | 0.87665 | 0.90511 | 0.930 |
| GM24385 | 0.88276 | 0.91143 | 0.935 |
| HG00733 | 0.87165 | 0.89682 | 0.925 |
| HG01109 | 0.87033 | 0.89845 | 0.930 |
| HG01243 | 0.88525 | 0.91435 | 0.935 |
| HG02055 | 0.87215 | 0.90572 | 0.930 |
| HG02080 | 0.88188 | 0.91259 | 0.935 |
| HG02723 | 0.84914 | 0.87565 | 0.920 |
| HG03098 | 0.85522 | 0.88156 | 0.915 |
| **All samples:** | **0.87251** | **0.90068** | **0.930** |

**Shasta: assembling a human genome from nanopore reads in under 6 hours**

Supplementary Table 4: QUAST assembly metrics of three samples on four assemblers, before polishing.

| Sample | Metric | Shasta | Wtdbg2 | Flye | Canu |
|---|---|---|---|---|---|
| HG00733 | # contigs | 2,150 | 5,086 | 1,852 | 778 |
| | Total length | 2,783,599,890 | 2,792,376,827 | 2,816,034,584 | 2,900,719,051 |
| | N50 | 24,429,871 | 18,763,119 | 28,763,002 | 44,759,083 |
| | NG50 | 21,088,309 | 15,338,021 | 25,227,330 | 40,627,903 |
| | # misassemblies | 814 | 3,985 | 6,555 | 4,570 |
| | Genome fraction (%) | 94.982 | 92.938 | 95.763 | 96.404 |
| | Duplication ratio | 0.995 | 1.005 | 0.986 | 1.014 |
| | # mismatches per 100 kbp | 156.21 | 248.78 | 506.12 | 231.24 |
| | # indels per 100 kbp | 453.97 | 664.90 | 1,480.91 | 677.26 |
| | Total aligned length | 2,775,307,347 | 2,742,343,142 | 2,769,440,009 | 2,858,769,830 |
| | NA50 | 16,052,981 | 9,106,500 | 18,577,806 | 21,157,324 |
| | NGA50 | 12,765,264 | 7,787,949 | 16,267,214 | 19,945,150 |
| HG002 | # contigs | 1,847 | 5,310 | 1,627 | 767 |
| | Total length | 2,801,200,983 | 2,793,889,694 | 2,819,241,152 | 2,901,099,163 |
| | N50 | 23,346,484 | 15,380,722 | 31,253,170 | 33,064,788 |
| | NG50 | 20,205,529 | 13,750,884 | 25,917,293 | 32,340,595 |
| | # misassemblies | 901 | 3,572 | 5,881 | 3,882 |
| | Genome fraction (%) | 95.622 | 93.136 | 96.228 | 96.959 |
| | Duplication ratio | 0.995 | 1.004 | 0.981 | 1.009 |
| | # mismatches per 100 kbp | 167.75 | 261.72 | 549.10 | 231.39 |
| | # indels per 100 kbp | 520.33 | 796.71 | 1,650.63 | 792.45 |
| | Total aligned length | 2,792,458,737 | 2,743,401,414 | 2,768,347,339 | 2,863,787,213 |
| | NA50 | 16,068,951 | 8,564,600 | 18,803,788 | 21,330,391 |
| | NGA50 | 14,189,972 | 7,361,363 | 16,079,132 | 18,175,258 |
| CHM13 | # contigs | 1,236 | 6,428 | 1,269 | 558 |
| | Total length | 2,809,087,051 | 2,836,802,421 | 2,857,931,691 | 2,919,690,848 |
| | N50 | 46,037,322 | 15,522,332 | 36,829,446 | 80,507,947 |
| | NG50 | 41,091,906 | 14,039,241 | 35,319,460 | 79,504,166 |
| | # misassemblies | 1,051 | 4,202 | 5,452 | 4,768 |
| | Genome fraction (%) | 95.307 | 93.124 | 96.022 | 96.553 |
| | Duplication ratio | 1.000 | 1.017 | 0.997 | 1.014 |
| | # mismatches per 100 kbp | 155.15 | 256.17 | 443.85 | 226.04 |
| | # indels per 100 kbp | 358.45 | 535.46 | 1,023.79 | 484.46 |
| | Total aligned length | 2,798,043,587 | 2,780,449,715 | 2,807,157,420 | 2,864,418,837 |
| | NA50 | 23,475,255 | 6,786,237 | 18,991,999 | 25,611,947 |
| | NGA50 | 18,990,051 | 5,892,796 | 17,032,972 | 23,819,455 |

Supplementary Table 5: QUAST misassembly count for four assemblers on different regions of the genome for four samples. We report misassemblies that happen in whole genome then we incrementally exclude Centromeric regions, segmental duplication regions (Seg Dups) and regions where Structural Variations (SVs) occur from the total misassemblies. We only have known SV regions for HG002 so other numbers are missing from the table.

| Sample | Assembler | Misassemblies in GRCh38 autosomes and chrX, chrY | Misassemblies outside Centromeres | Misassemblies outside Centromeres and Seg Dups | Misassemblies outside Centromeres and Seg Dups and known SVs |
|---|---|---|---|---|---|
| HG002 | Shasta | 901 | 755 | 284 | 111 |
| | Flye | 5881 | 1226 | 513 | 360 |
| | Canu | 3882 | 2347 | 689 | 320 |
| | Wtdbg2 | 3572 | 1213 | 484 | 254 |
| HG00733 | Shasta | 814 | 662 | 256 | - |
| | Flye | 6555 | 1261 | 604 | - |
| | Canu | 4570 | 2791 | 755 | - |
| | Wtdbg2 | 3985 | 1166 | 474 | - |
| CHM13 | Shasta | 1051 | 795 | 333 | - |
| | Flye | 5452 | 1228 | 448 | - |
| | Canu | 4768 | 2764 | 864 | - |
| | Wtdbg2 | 4202 | 1519 | 592 | - |

Supplementary Table 6: BAC analysis on selected dataset. BACs were selected (31 of CHM13 and 16 of HG00733) for falling within unique regions of the genome, specifically >10 Kb away from the closest segmental duplication. *Closed* refers to the number of BACs for which 99.5% of their length aligns to a single locus in the assembly. *Attempted* refers to the number of BACs which have an alignment for >5 Kb of sequence with >90% identity to only one contig (BACs which have such alignments to multiple contigs are excluded). Identity metrics are for *closed* BACs.

| Sample | Assembler | BAC counts | | | | Median Quality | | Mean Quality | |
|---|---|---|---|---|---|---|---|---|---|
| | | Total | Attempted | Closed | Closed of attempted % | Identity % | QV | Identity % | QV |
| CHM13 | Canu | 31 | 31 | 30 | 96.77 | 99.40 | 22.18 | 99.34 | 21.84 |
| | Flye | 31 | 31 | 31 | 100.00 | 97.58 | 16.17 | 97.65 | 16.28 |
| | Shasta | 31 | 31 | 31 | 100.00 | 99.55 | 23.51 | 99.51 | 23.07 |
| | Wtdbg2 | 31 | 29 | 28 | 96.55 | 99.46 | 22.71 | 99.39 | 22.15 |
| HG00733 | Canu | 16 | 16 | 15 | 93.75 | 98.74 | 18.98 | 98.61 | 18.56 |
| | Flye | 16 | 16 | 15 | 93.75 | 98.02 | 17.03 | 98.06 | 17.12 |
| | Shasta | 16 | 16 | 15 | 93.75 | 98.89 | 19.53 | 98.85 | 19.4 |
| | Wtdbg2 | 16 | 16 | 15 | 93.75 | 98.82 | 19.29 | 98.84 | 19.37 |

Supplementary Table 7: BAC analysis on full dataset, 341 on CHM13 and 179 on HG00733. *Closed* refers to the number of BACs for which 99.5% of their length aligns to a single locus. *Attempted* refers to the number of BACs which have an alignment for >5Kb of sequence with >90% identity to only one contig (BACs which have such alignments to multiple contigs are excluded). Identity metrics are for *closed* BACs.

| Sample | Assembler Polisher | BAC counts | | | | Median Quality | | Mean Quality | |
|---|---|---|---|---|---|---|---|---|---|
| | | Total | Attempted | Closed | Closed of attempted % | Identity % | QV | Identity % | QV |
| CHM13 | Canu | 341 | 315 | 282 | 89.52 | 99.22 | 21.07 | 98.93 | 19.7 |
| | Flye | 341 | 235 | 202 | 85.95 | 97.54 | 16.09 | 97.51 | 16.03 |
| | Shasta | 341 | 104 | 92 | 88.46 | 99.47 | 22.74 | 99.37 | 21.99 |
| | Wtdbg2 | 341 | 89 | 62 | 69.66 | 99.36 | 21.96 | 99.28 | 21.43 |
| HG00733 | Canu | 179 | 140 | 119 | 85 | 98.73 | 18.95 | 98.43 | 18.05 |
| | Flye | 179 | 99 | 78 | 78.78 | 98.09 | 17.18 | 97.76 | 16.49 |
| | Shasta | 179 | 46 | 39 | 84.78 | 98.78 | 19.12 | 98.14 | 17.31 |
| | Wtdbg2 | 179 | 74 | 45 | 60.81 | 98.71 | 18.91 | 98.02 | 17.04 |

Supplementary Table 8: Base-level accuracies on four different assemblers for two samples. Analysis is performed with whole-genome truth sequences.

| Sample | Assembler | Percentage Errors | | | |
|---|---|---|---|---|---|
| | | Balanced | Identity | Deletion | Insertion |
| HG00733 Guppy 2.3.5 | Shasta | 1.217% | 0.084% | 0.963% | 0.170% |
| | Wtdbg2 | 1.381% | 0.122% | 1.159% | 0.101% |
| | Canu | 1.499% | 0.090% | 1.328% | 0.082% |
| | Flye | 2.030% | 0.100% | 0.536% | 1.395% |
| CHM13 Guppy 2.3.1 | Shasta | 0.626% | 0.048% | 0.469% | 0.109% |
| | Wtdbg2 | 0.907% | 0.109% | 0.709% | 0.090% |
| | Canu | 0.800% | 0.056% | 0.692% | 0.052% |
| | Flye | 2.363% | 0.071% | 0.501% | 1.791% |

Supplementary Table 9: Runtime and cost of three assembly workflows on Amazon Web Services (AWS) platform.

| Method | Sample | Minutes | Threads Used | Peak Memory | AWS Instance Type | AWS Instance Cost |
|---|---|---|---|---|---|---|
| WTDBG2 | HG00733 | 2971 | 63 | 365 | x1.16xlarge | $6.67 |
| | GM24385 | 1752 | 63 | 293 | x1.16xlarge | $6.67 |
| | CHM13 | 1655 | 63 | 312 | x1.16xlarge | $6.67 |
| WTDBG2 (wtpoa-cns) | HG00733 | 248 | 31 | 12 | x1.16xlarge | $6.67 |
| | GM24385 | 274 | 24 | 12 | x1.16xlarge | $6.67 |
| | CHM13 | 257 | 31 | 12 | x1.16xlarge | $6.67 |
| Flye | HG00733 | 3421 | 123 | 1013 | x1.16xlarge | $6.67 |
| | GM24385 | 3749 | 64 | 727 | x1.16xlarge | $6.67 |
| | CHM13 | 4084 | 126 | 911 | x1.16xlarge | $6.67 |
| Shasta | HG00733 | 300 | 128 | 966 | x1.32xlarge | $13.34 |
| | HG01109 | 354 | 128 | - | x1.32xlarge | $13.34 |
| | HG01243 | 294 | 128 | - | x1.32xlarge | $13.34 |
| | HG02055 | 312 | 128 | - | x1.32xlarge | $13.34 |
| | HG02080 | 276 | 128 | - | x1.32xlarge | $13.34 |
| | HG02723 | 372 | 128 | - | x1.32xlarge | $13.34 |
| | HG03098 | 240 | 128 | - | x1.32xlarge | $13.34 |
| | HG03492 | 198 | 128 | - | x1.32xlarge | $13.34 |
| | GM24385 | 240 | 128 | 692 | x1.32xlarge | $13.34 |
| | GM24149 | 426 | 128 | - | x1.32xlarge | $13.34 |
| | GM24143 | 450 | 128 | - | x1.32xlarge | $13.34 |
| | CHM13 | 336 | 128 | - | x1.32xlarge | $13.34 |

#### Contiguously assembling MHC haplotypes

Supplementary Table 10: CHM13 MHC unpolished Shasta assembly as compared to the nearest matching haplotype in hg38 (GL000251.2)

| Assembler | Best Contig | Misassemblies | Largest Aligned | Mismatch Rate | Indel Rate |
|---|---|---|---|---|---|
| Shasta | 62 | 6 | 2,788,362 | 0.00296 | 0.00399 |
| Canu | tig00589784 | 5 | 2,792,139 | 0.00331 | 0.00607 |
| Flye | contig_115 | 6 | 2,787,570 | 0.00543 | 0.01106 |
| wtdbg2 | ctg25 | 32 | 1,819,753 | 0.00553 | 0.00576 |

Supplementary Table 11: QUAST results for the HG00733 trio-binned maternal reads, using all four assemblers.

| Metric | HG00733-Mother | | | |
|---|---|---|---|---|
| | Shasta | Wtdbg2 | Flye (initial) | Canu |
| # contigs | 1,934 | 4,028 | 1,634 | 877 |
| Total length | 2,754,225,214 | 2,690,619,717 | 2,791,893,188 | 2,829,920,708 |
| N50 | 9,071,623 | 14,125,235 | 25,658,831 | 19,451,828 |
| NG50 | 7,702,138 | 10,217,387 | 23,775,989 | 16,507,795 |
| # misassemblies | 705 | 3,661 | 6,082 | 2,161 |
| Genome fraction (%) | 90.824 | 87.373 | 92.121 | 92.298 |
| Duplication ratio | 0.993 | 0.996 | 0.982 | 0.999 |
| # mismatches per 100 kbp | 194.15 | 287.89 | 549.61 | 232.72 |
| # indels per 100 kbp | 576.55 | 859.83 | 1585.30 | 724.67 |
| Total aligned length | 2,748,135,723 | 2,650,821,801 | 2,751,532,754 | 2,798,797,021 |
| NA50 | 7,805,090 | 7,615,651 | 15,615,208 | 11,947,316 |
| NGA50 | 6,339,949 | 5,584,544 | 12,833,996 | 10,085,023 |

Supplementary Table 12: HG00733 Maternal trio binned MHC unpolished Shasta assembly as compared to the nearest matching haplotype in hg38 (GL000255.1)

| Assembler | Best Contig | Misassemblies | Largest Aligned | Mismatch Rate | Indel Rate |
|---|---|---|---|---|---|
| Shasta | 226 | 0 | 4,289,729 | 0.00206 | 0.00538 |
| Canu | tig00002130 | 0 | 4,289,729 | 0.00182 | 0.00676 |
| Flye | contig_295 | 0 | 4,289,729 | 0.00579 | 0.01759 |
| wtdbg2 | ctg36 | 23 | 1,418,939 | 0.00592 | 0.00905 |

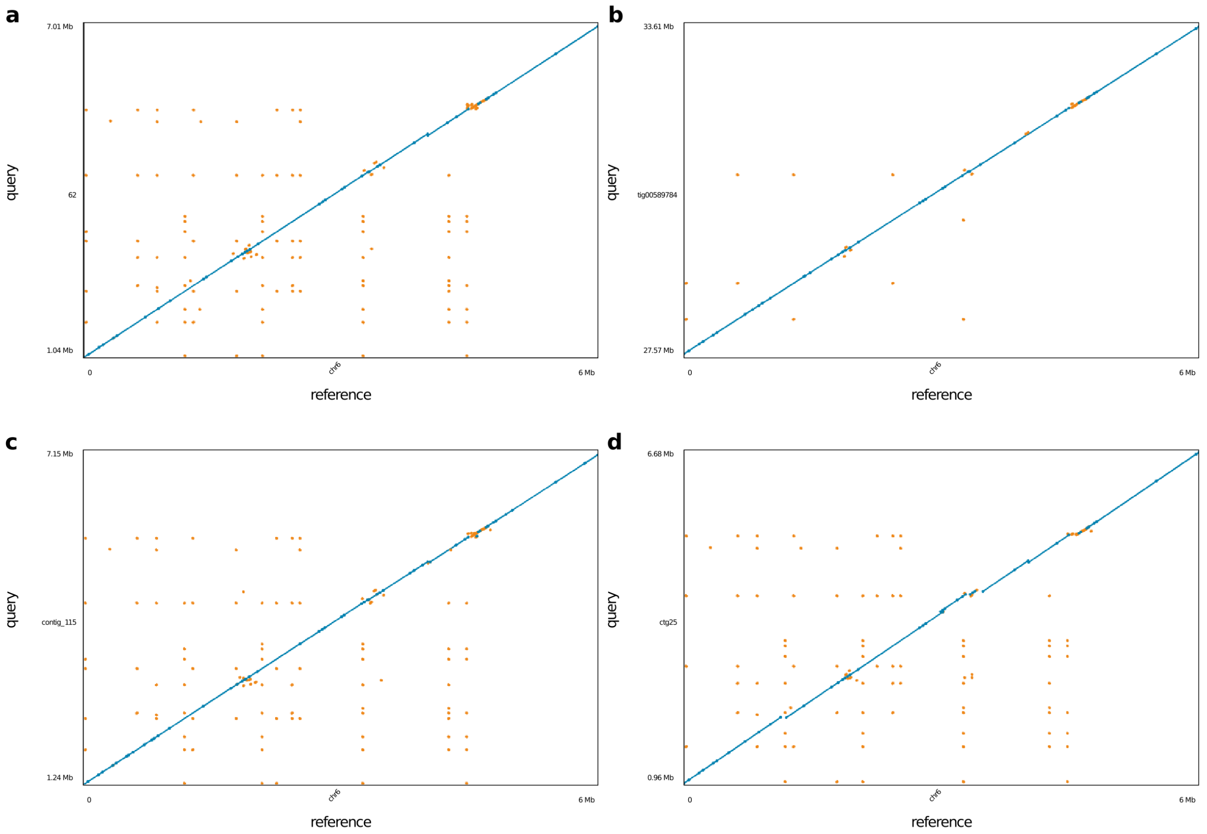


Supplementary Figure 1: Dotplot of unpolished CHM13 MHC assembly vs hg38 chr6:28000000-34000000 for the each of the 4 assemblers tested. **(a)** Shasta **(b)** Canu **(c)** Flye (no native polish) **(d)** wtdbg2. Blue dots represent unique alignments and orange dots represent repetitive alignments.

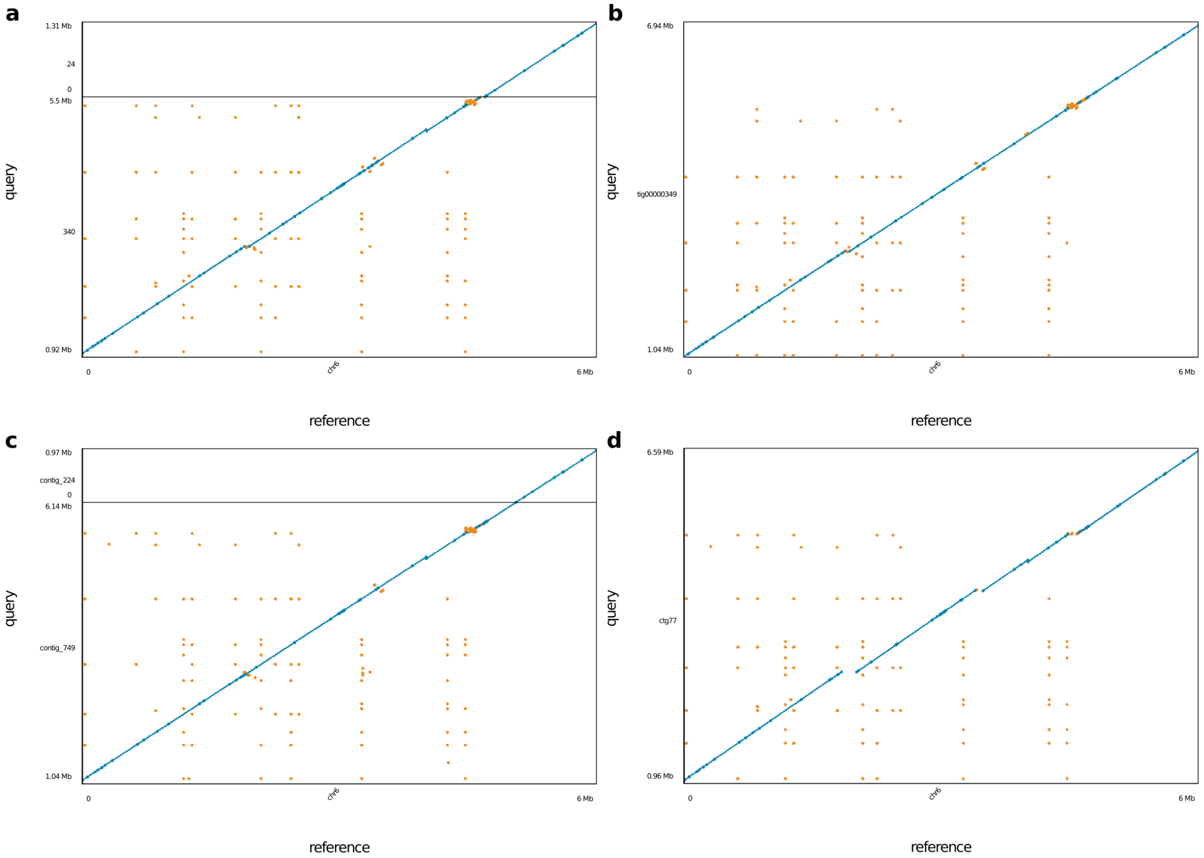


Supplementary Figure 2: Dotplot of unpolished HG00733 diploid MHC assembly vs hg38 chr6:28000000-34000000 for the each of the 4 assemblers tested. **(a)** Shasta **(b)** Canu **(c)** Flye (no native polish) **(d)** wtdbg2. Blue dots represent unique alignments and orange dots represent repetitive alignments.

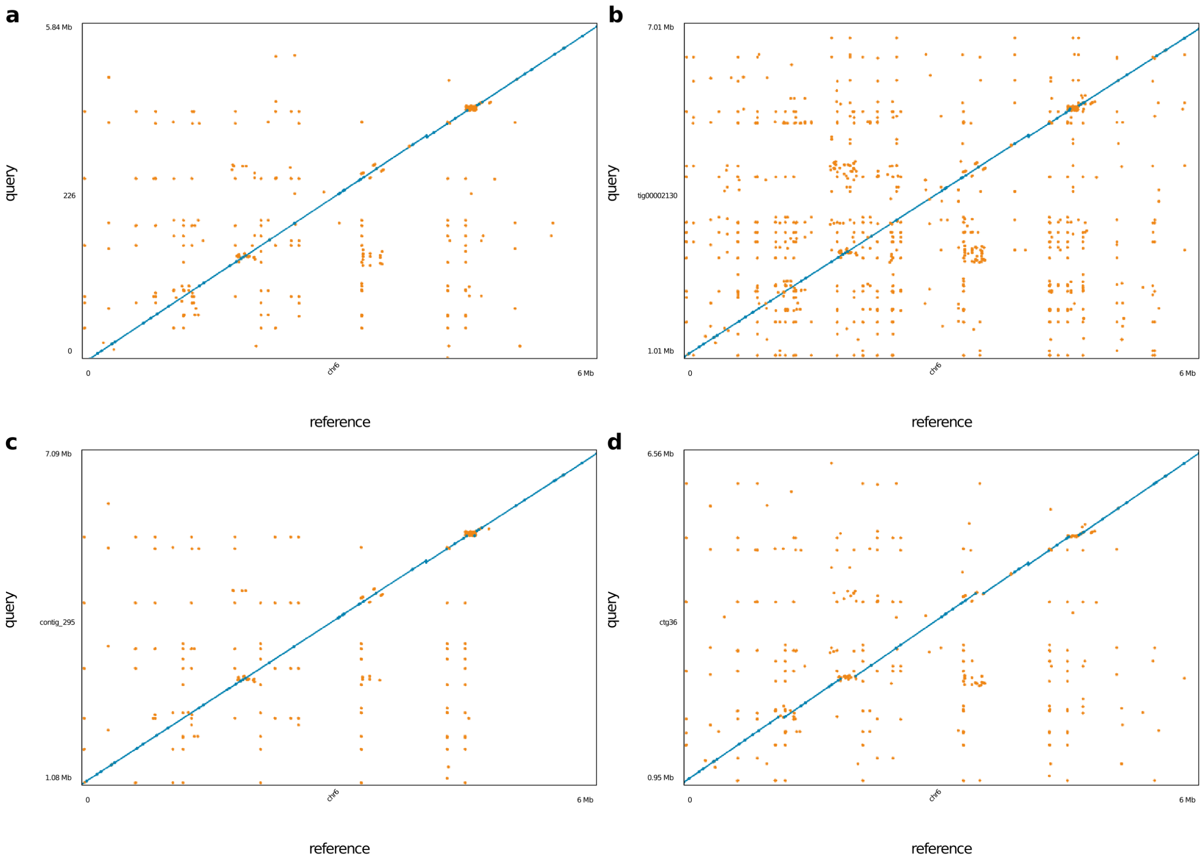


Supplementary Figure 3: Dotplot of unpolished HG00733 maternal haploid MHC assembly vs hg38 chr6:28000000-34000000 for the each of the 4 assemblers tested. **(a)** Shasta **(b)** Canu **(c)** Flye (no native polish) **(d)** wtdbg2. Blue dots represent unique alignments and orange dots represent repetitive alignments.

**Deep neural network based polishing achieves QV30 long-read only polishing accuracy**

Supplementary Table 13: Base-level accuracies comparing Racon & Medaka and MarginPolish & HELEN pipelines on Shasta assemblies for three samples. Analysis is performed with whole-genome truth sequences.

| Sample | Polisher | | Percentage Errors | | | |
|---|---|---|---|---|---|---|
| | Method | Model | Balanced | Identity | Deletion | Insertion |
| HG002 Guppy 2.3.5 | Shasta | Unpolished | 1.030% | 0.058% | 0.861% | 0.111% |
| | Racon | 4x | 0.711% | 0.052% | 0.584% | 0.075% |
| | Medaka | r941_flip235 | 0.447% | 0.049% | 0.324% | 0.074% |
| | MarginPolish | guppy_ff235 | 0.432% | 0.042% | 0.268% | 0.122% |
| | HELEN | rl941_flip235 | 0.347% | 0.042% | 0.186% | 0.119% |
| HG00733 Guppy 2.3.5 | Shasta | Unpolished | 1.217% | 0.084% | 0.963% | 0.170% |
| | Racon | 4x | 0.839% | 0.085% | 0.624% | 0.131% |
| | Medaka | r941_flip235 | 0.579% | 0.082% | 0.382% | 0.116% |
| | MarginPolish | guppy_ff235 | 0.586% | 0.069% | 0.340% | 0.177% |
| | HELEN | rl941_flip235 | 0.501% | 0.069% | 0.262% | 0.170% |
| CHM13 Guppy 2.3.1 | Shasta | Unpolished | 0.626% | 0.048% | 0.469% | 0.109% |
| | Racon | 4x | 0.498% | 0.074% | 0.241% | 0.183% |
| | Medaka | r941_flip213 | 0.417% | 0.051% | 0.065% | 0.302% |
| | MarginPolish | guppy_ff233 | 0.384% | 0.052% | 0.107% | 0.224% |
| | HELEN | rl941_flip233 | 0.299% | 0.052% | 0.101% | 0.146% |

Supplementary Table 14: QUAST results for the Shasta assemblies for all samples, post polishing with MarginPolish-HELEN.

| Sample | # contigs | Total length | N50 | NG50 | # mis-assemblies | Genome fraction (%) | # mismatches per 100 kbp | # indels per 100 kbp | Total aligned length | NA50 | NGA50 |
|---|---|---|---|---|---|---|---|---|---|---|---|
| GM24143 | 2,042 | 2,802,437,249 | 23,531,777 | 19,936,924 | 970 | 95.025 | 128.63 | 142.77 | 2,794,379,803 | 16,323,510 | 13,840,294 |
| GM24149 | 2,368 | 2,816,566,939 | 20,798,256 | 17,752,973 | 990 | 95.416 | 130.54 | 134.60 | 2,806,847,428 | 13,174,778 | 12,128,076 |
| GM24385 | 1,685 | 2,819,474,365 | 23,520,830 | 20,346,145 | 960 | 95.609 | 127.44 | 152.17 | 2,810,951,083 | 16,200,287 | 14,315,298 |
| HG00733 | 1,962 | 2,800,357,697 | 24,600,414 | 21,701,762 | 877 | 94.976 | 126.23 | 137.92 | 2,792,792,711 | 16,156,822 | 12,971,070 |
| HG01109 | 2,111 | 2,820,988,852 | 21,532,001 | 18,279,481 | 1,033 | 95.564 | 136.51 | 140.59 | 2,811,696,923 | 13,162,850 | 12,012,786 |
| HG01243 | 1,936 | 2,819,065,027 | 22,753,128 | 20,884,160 | 920 | 95.521 | 137.50 | 143.02 | 2,810,262,570 | 16,040,951 | 14,115,348 |
| HG02055 | 1,903 | 2,819,836,390 | 17,485,643 | 16,302,857 | 971 | 95.592 | 142.23 | 162.43 | 2,810,300,557 | 13,840,319 | 12,123,357 |
| HG02080 | 1,814 | 2,803,471,776 | 18,701,305 | 15,584,440 | 920 | 95.045 | 128.16 | 134.35 | 2,794,749,368 | 12,401,739 | 11,561,569 |
| HG02723 | 1,813 | 2,805,268,038 | 25,163,327 | 20,265,678 | 1,110 | 95.062 | 143.30 | 147.09 | 2,796,332,696 | 15,390,923 | 13,175,818 |
| HG03098 | 1,790 | 2,811,295,217 | 22,571,315 | 19,620,076 | 986 | 95.395 | 144.36 | 170.40 | 2,802,844,336 | 14,045,283 | 12,089,849 |
| HG03492 | 1,811 | 2,811,690,127 | 24,629,163 | 22,891,947 | 854 | 95.364 | 126.61 | 147.22 | 2,804,103,412 | 16,317,390 | 12,930,516 |
| CHM13 | 1,186 | 2,819,245,173 | 46,206,794 | 41,255,275 | 1,107 | 95.281 | 136.58 | 140.38 | 2,808,536,514 | 23,540,225 | 19,532,176 |

Supplementary Table 15: Base-level accuracies comparing Racon & Medaka and MarginPolish & HELEN pipelines against CHM13 Chromosome-X. The truth Chromosome-X sequence used reflects the most accurate haploid truth sequence available.

| Sample | Polisher | | Percentage Errors | | | |
|---|---|---|---|---|---|---|
| | Method | Model | Balanced | Identity | Deletion | Insertion |
| CHM-13 Chromosome-X | Shasta | Unpolished | 0.494% | 0.015% | 0.419% | 0.060% |
| | Racon | 4x | 0.340% | 0.021% | 0.204% | 0.115% |
| | Medaka | r941_flip213 | 0.127% | 0.013% | 0.044% | 0.070% |
| | MarginPolish | guppy_ff233 | 0.269% | 0.024% | 0.067% | 0.178% |
| | HELEN | rl941_flip233 | 0.167% | 0.011% | 0.055% | 0.101% |
| | | rl941_flip231 | 0.095% | 0.009% | 0.053% | 0.034% |

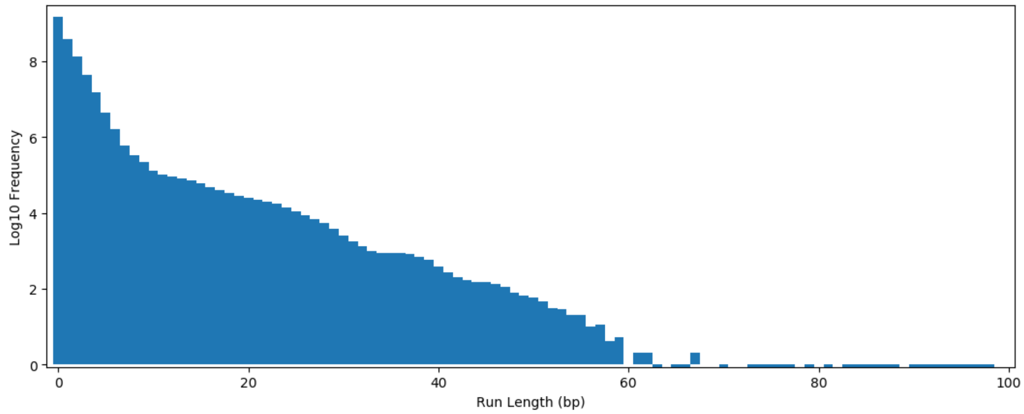


Supplementary Figure 4: Log frequency of each run length as found in the GRCh38 reference for all bases A,C,G,T up to 100bp. Run lengths greater than 15 account for approximately 0.012% of all homopolymer runs in GRCh38.

Supplementary Table 16: Base-level accuracies improvements with MarginPolish and HELEN pipeline on four different assemblers for two samples. Analysis is performed with whole-genome truth sequences.

| Sample | Polisher | | Percentage Errors | | | |
|---|---|---|---|---|---|---|
| | Method | Model | Balanced | Identity | Deletion | Insertion |
| HG00733 Guppy 2.3.5 | Shasta | Unpolished | 1.217% | 0.084% | 0.963% | 0.170% |
| | MarginPolish | guppy_ff235 | 0.586% | 0.069% | 0.340% | 0.177% |
| | HELEN | rl941_flip235 | 0.501% | 0.069% | 0.262% | 0.170% |
| | Wtdbg2 | Unpolished | 1.381% | 0.122% | 1.159% | 0.101% |
| | MarginPolish | guppy_ff235 | 0.665% | 0.080% | 0.422% | 0.163% |
| | HELEN | rl941_flip235 | 0.587% | 0.081% | 0.341% | 0.164% |
| | Canu | Unpolished | 1.499% | 0.090% | 1.328% | 0.082% |
| | MarginPolish | guppy_ff235 | 0.576% | 0.064% | 0.364% | 0.148% |
| | HELEN | rl941_flip235 | 0.491% | 0.063% | 0.279% | 0.149% |
| | Flye | Unpolished | 2.030% | 0.100% | 0.536% | 1.395% |
| | MarginPolish | guppy_ff235 | 0.571% | 0.075% | 0.339% | 0.157% |
| | HELEN | rl941_flip235 | 0.494% | 0.075% | 0.263% | 0.157% |
| CHM13 Guppy 2.3.1 | Shasta | Unpolished | 0.626% | 0.048% | 0.469% | 0.109% |
| | MarginPolish | guppy_ff233 | 0.384% | 0.052% | 0.107% | 0.224% |
| | HELEN | rl941_flip233 | 0.299% | 0.052% | 0.101% | 0.146% |
| | Wtdbg2 | Unpolished | 0.907% | 0.109% | 0.709% | 0.090% |
| | MarginPolish | guppy_ff233 | 0.530% | 0.056% | 0.235% | 0.239% |
| | HELEN | rl941_flip233 | 0.468% | 0.057% | 0.243% | 0.168% |
| | Canu | Unpolished | 0.800% | 0.056% | 0.692% | 0.052% |
| | MarginPolish | guppy_ff233 | 0.340% | 0.027% | 0.107% | 0.206% |
| | HELEN | rl941_flip233 | 0.268% | 0.029% | 0.097% | 0.143% |
| | Flye | Unpolished | 2.363% | 0.071% | 0.501% | 1.791% |
| | MarginPolish | guppy_ff233 | 0.344% | 0.041% | 0.088% | 0.216% |
| | HELEN | rl941_flip233 | 0.277% | 0.041% | 0.086% | 0.150% |

Supplementary Table 17: Runtime and cost of two polishing workflows on Amazon Web Services (AWS) platform.

| Method | Sample | Minutes | Threads Used | Peak Memory | Instance Type | Instance Cost |
|---|---|---|---|---|---|---|
| Racon (4x) | HG00733 | 3099 | 62 | 574 | x1.16xlarge | $6.67 |
| | GM24385 | 2342 | 62 | 501 | x1.16xlarge | $6.67 |
| | CHM13 | 3700 | 62 | 281 | x1.16xlarge | $6.67 |
| Medaka mini_align | HG00733 | 611 | 62 | 101 | x1.16xlarge | $6.67 |
| | GM24385 | 489 | 62 | 115 | x1.16xlarge | $6.67 |
| | CHM13 | 810 | 60 | 143 | x1.16xlarge | $6.67 |
| Medaka call_consensus | HG00733 | 8611 | 62 | 164 | c5n.18xlarge | $3.89 |
| | GM24385 | 3355 | 62 | 150 | c5n.18xlarge | $3.89 |
| | CHM13 | 2532 | 62 | 149 | c5n.18xlarge | $3.89 |
| MarginPolish | HG00733 | 680 | 90 | 66 | m5.metal | $4.61 |
| | HG01109 | 912 | 70 | 57 | c5.18xlarge | $3.06 |
| | HG01243 | 835 | 70 | 65 | c5.18xlarge | $3.06 |
| | HG02055 | 733 | 70 | 77 | c5.18xlarge | $3.06 |
| | HG02080 | 793 | 70 | 64 | c5.18xlarge | $3.06 |
| | HG02723 | 1000 | 64 | 60 | c5.18xlarge | $3.06 |
| | HG03098 | 852 | 70 | 78 | c5.18xlarge | $3.06 |
| | HG03492 | 777 | 70 | 80 | c5.18xlarge | $3.06 |
| | GM24385 | 842 | 70 | 66 | c5.18xlarge | $3.06 |
| | GM24149 | 1037 | 64 | 103 | c5.18xlarge | $3.06 |
| | GM24143 | 1051 | 64 | 84 | c5.18xlarge | $3.06 |
| | CHM13 | 739 | 70 | 65 | c5.18xlarge | $3.06 |
| HELEN consensus | HG00733 | 216 | 8 GPUs | - | p2.8xlarge | $7.20 |
| | HG01109 | 204 | 8 GPUs | - | p2.8xlarge | $7.20 |
| | HG01243 | 233 | 8 GPUs | - | p2.8xlarge | $7.20 |
| | HG02080 | 212 | 8 GPUs | - | p2.8xlarge | $7.20 |
| | HG03098 | 216 | 8 GPUs | - | p2.8xlarge | $7.20 |
| | GM24385 | 208 | 8 GPUs | - | p2.8xlarge | $7.20 |
| | GM24143 | 226 | 8 GPUs | - | p2.8xlarge | $7.20 |
| HELEN stitch | HG00733 | 59 | 32 | - | p2.8xlarge | $7.20 |
| | HG01109 | 50 | 32 | - | p2.8xlarge | $7.20 |
| | HG01243 | 49 | 32 | - | p2.8xlarge | $7.20 |
| | HG02080 | 54 | 32 | - | p2.8xlarge | $7.20 |
| | HG03098 | 65 | 32 | - | p2.8xlarge | $7.20 |
| | GM24385 | 68 | 32 | - | p2.8xlarge | $7.20 |
| | GM24143 | 62 | 32 | - | p2.8xlarge | $7.20 |

**Long-read assemblies contain nearly all human coding genes**

Supplementary Table 18: Transcript-level analysis with Comparative Annotation Toolkit (CAT) of MarginPolish & HELEN and Racon & Medaka on three samples from Shasta assemblies.

| Metric | | HG002 | | HG00733 | | CHM13 | |
|---|---|---|---|---|---|---|---|
| | | HELEN | MEDAKA | HELEN | MEDAKA | HELEN | MEDAKA |
| Transcripts Found | Total | 83093 | 83105 | 83002 | 82928 | 82833 | 82807 |
| | Percent | 99.536 | 99.551 | 99.427 | 99.339 | 99.225 | 99.194 |
| Full mRNA Coverage | Total | 25721 | 20367 | 28612 | 26573 | 40132 | 38081 |
| | Percent | 30.811 | 24.397 | 34.274 | 31.832 | 48.074 | 45.617 |
| Full CDS Coverage | Total | 41396 | 36248 | 45104 | 43956 | 53089 | 52297 |
| | Percent | 49.588 | 43.421 | 54.030 | 52.655 | 63.595 | 62.646 |
| Transcripts With Frameshift | Total | 35339 | 40783 | 31333 | 32647 | 23261 | 24441 |
| | Percent | 42.332 | 48.854 | 37.534 | 39.108 | 27.864 | 29.278 |
| Transcripts With Original Introns | Total | 76880 | 76883 | 76618 | 76463 | 76807 | 76803 |
| | Percent | 92.094 | 92.098 | 91.780 | 91.594 | 92.006 | 92.002 |
| Transcripts With Full CDS Coverage | Total | 41396 | 36248 | 45104 | 43956 | 53089 | 52297 |
| | Percent | 49.588 | 43.421 | 54.030 | 52.655 | 63.595 | 62.646 |
| Transcripts With Full CDS Coverage And No Frameshifts | Total | 41245 | 36158 | 44982 | 43860 | 52966 | 52160 |
| | Percent | 49.407 | 43.313 | 53.884 | 52.540 | 63.448 | 62.482 |
| Transcripts With Full CDS Coverage And No Frameshifts And Original Introns | Total | 41021 | 35952 | 44692 | 43546 | 52616 | 51807 |
| | Percent | 49.139 | 43.067 | 53.536 | 52.163 | 63.028 | 62.059 |

Supplementary Table 19: Gene-level analysis with Comparative Annotation Toolkit (CAT) of MarginPolish & HELEN and Racon & Medaka on three samples from Shasta assemblies.

| Metric | | HG002 | | HG00733 | | CHM13 | |
|---|---|---|---|---|---|---|---|
| | | HELEN | MEDAKA | HELEN | MEDAKA | HELEN | MEDAKA |
| Genes Found | Total | 19536 | 19531 | 19537 | 19511 | 19505 | 19490 |
| | Percent | 99.268 | 99.243 | 99.273 | 99.141 | 99.111 | 99.035 |
| Genes With Frameshift | Total | 10933 | 12165 | 9941 | 10081 | 7300 | 7564 |
| | Percent | 55.554 | 61.814 | 50.513 | 51.225 | 37.093 | 38.435 |
| Genes With Original Introns | Total | 18212 | 18198 | 18151 | 18113 | 18217 | 18202 |
| | Percent | 92.541 | 92.47 | 92.231 | 92.038 | 92.566 | 92.49 |
| Genes With Full CDS Coverage | Total | 11070 | 10066 | 11812 | 11756 | 13648 | 13534 |
| | Percent | 56.25 | 51.148 | 60.02 | 59.736 | 69.35 | 68.77 |
| Genes With Full CDS Coverage And No Frameshifts | Total | 12454 | 11570 | 13127 | 13081 | 14625 | 14562 |
| | Percent | 63.283 | 58.791 | 66.702 | 66.468 | 74.314 | 73.994 |
| Genes With Full CDS Coverage And No Frameshifts And Original Introns | Total | 12422 | 11539 | 13098 | 13042 | 14603 | 14531 |
| | Percent | 63.12 | 58.633 | 66.555 | 66.27 | 74.202 | 73.836 |
| Missing Genes | Total | 144 | 149 | 143 | 169 | 175 | 190 |
| | Percent | 0.732 | 0.757 | 0.727 | 0.859 | 0.889 | 0.965 |

Supplementary Table 20: Transcript-level analysis with Comparative Annotation Toolkit (CAT) of four HG00733 assemblies polished with MarginPolish and HELEN.

| Metric | | HG00733 | | | |
|---|---|---|---|---|---|
| | | Flye HELEN | Canu HELEN | Wtdbg2 HELEN | Shasta HELEN |
| Transcripts Found | Total | 83267 | 83334 | 81484 | 82974 |
| | Percent | 99.745 | 99.825 | 97.609 | 99.394 |
| Full mRNA Coverage | Total | 33078 | 28488 | 28889 | 30378 |
| | Percent | 39.624 | 34.126 | 34.606 | 36.390 |
| Full CDS Coverage | Total | 41396 | 44877 | 45321 | 46965 |
| | Percent | 59.754 | 53.758 | 54.290 | 56.259 |
| Transcripts With Frameshift | Total | 27293 | 32230 | 29525 | 29657 |
| | Percent | 32.694 | 38.608 | 35.368 | 35.526 |
| Transcripts With Original Introns | Total | 77412 | 77583 | 74683 | 76613 |
| | Percent | 92.731 | 92.936 | 89.462 | 91.774 |
| Transcripts with Full CDS Coverage | Total | 49883 | 44877 | 45321 | 46965 |
| | Percent | 59.754 | 53.758 | 54.290 | 56.259 |
| Transcripts with Full CDS Coverage And No Frameshifts | Total | 49766 | 44737 | 45217 | 46802 |
| | Percent | 59.614 | 53.590 | 54.165 | 56.064 |
| Transcripts with Full CDS Coverage And No Frameshifts And Original Introns | Total | 49459 | 44412 | 44924 | 46505 |
| | Percent | 59.247 | 53.201 | 53.814 | 55.708 |

Supplementary Table 21: Gene-level analysis with Comparative Annotation Toolkit (CAT) of four HG00733 assemblies polished with MarginPolish and HELEN

| Metric | | HG00733 | | | |
|---|---|---|---|---|---|
| | | Flye HELEN | Canu HELEN | Wtdbg2 HELEN | Shasta HELEN |
| Genes Found | Total | 19563 | 19629 | 19174 | 19528 |
| | Percent | 99.405 | 99.741 | 97.429 | 99.228 |
| Genes With Frameshift | Total | 8698 | 10160 | 9323 | 9464 |
| | Percent | 44.197 | 51.626 | 47.373 | 48.089 |
| Genes With Original Introns | Total | 18345 | 18460 | 17709 | 18154 |
| | Percent | 93.216 | 93.801 | 89.985 | 92.246 |
| Genes With Full CDS Coverage | Total | 12966 | 11889 | 11817 | 12207 |
| | Percent | 65.884 | 60.412 | 60.046 | 62.027 |
| Genes With Full CDS Coverage And No Frameshifts | Total | 14145 | 13221 | 13047 | 13419 |
| | Percent | 71.875 | 67.18 | 66.296 | 68.186 |
| Genes With Full CDS Coverage And No Frameshifts And Original Introns | Total | 14124 | 13193 | 13017 | 13396 |
| | Percent | 71.768 | 67.038 | 66.143 | 68.069 |
| Missing Genes | Total | 117 | 51 | 506 | 152 |
| | Percent | 0.595 | 0.259 | 2.571 | 0.772 |

Supplementary Table 22: BUSCO results of three samples using two polishing workflows on Shasta assemblies.

| Sample | Metric | Shasta MarginPolish HELEN | Shasta Racon (4x) Medaka |
|---|---|---|---|
| HG00733 | Complete BUSCOs (C) | 87.20% | 87.10% |
| | Complete and single-copy BUSCOs (S) | 84.20% | 83.80% |
| | Complete and duplicated BUSCOs (D) | 3.00% | 3.30% |
| | Fragmented BUSCOs (F) | 4.60% | 5.30% |
| | Missing BUSCOs (M) | 8.20% | 7.60% |
| HG002 | Complete BUSCOs (C) | 89.40% | 88.80% |
| | Complete and single-copy BUSCOs (S) | 84.80% | 85.80% |
| | Complete and duplicated BUSCOs (D) | 4.60% | 3.00% |
| | Fragmented BUSCOs (F) | 3.60% | 4.30% |
| | Missing BUSCOs (M) | 7.00% | 6.90% |
| CHM13 | Complete BUSCOs (C) | 86.50% | 86.80% |
| | Complete and single-copy BUSCOs (S) | 82.50% | 82.80% |
| | Complete and duplicated BUSCOs (D) | 4.00% | 4.00% |
| | Fragmented BUSCOs (F) | 5.90% | 5.30% |
| | Missing BUSCOs (M) | 7.60% | 7.90% |

Supplementary Table 23: BUSCO results for four assemblers on HG00733, post polishing with MarginPolish and HELEN.

| Metric | HG00733 | | | |
|---|---|---|---|---|
| | Flye | Canu | Wtdbg2 | Shasta |
| Complete BUSCOs (C) | 87.50% | 89.80% | 85.80% | 87.20% |
| Complete and single-copy BUSCOs (S) | 84.50% | 86.80% | 82.20% | 84.20% |
| Complete and duplicated BUSCOs (D) | 3.00% | 3.00% | 3.60% | 3.00% |
| Fragmented BUSCOs (F) | 5.30% | 3.00% | 6.30% | 4.60% |
| Missing BUSCOs (M) | 7.20% | 7.20% | 7.90% | 8.20% |

#### Comparing to a PacBio HiFi Assembly

Supplementary Table 24: CHM13 QUAST results for Shasta, MarginPolish, HELEN and PacBio HiFi assembly. Stratified misassembly counts were added after manual determination.

| Metric | CHM13 | |
|---|---|---|
| | Nanopore Shasta MarginPolish, HELEN | PacBio-HiFi Canu Racon |
| # contigs | 1622 | 5206 |
| Total length | 2819245173 | 3031026325 |
| N50 | 46206794 | 29522819 |
| NG50 | 41255275 | 29092230 |
| # misassemblies | 1107 | 8666 |
| # misassemblies outside Centromeres | 801 | 2999 |
| # misassemblies outside centromeres and Seg Dups | 314 | 893 |
| Genome fraction (%) | 95.281 | 97.030 |
| # mismatches per 100 kbp | 136.58 | 274.84.91 |
| # indels per 100 kbp | 140.38 | 32.99 |
| Total aligned length | 2808536514 | 2954558720 |
| NA50 | 23540225 | 20440378 |
| NGA50 | 19532176 | 20029136 |

Supplementary Table 25: CHM13 Chromosome-X error rate analysis with Pomoxis for Shasta, MarginPolish, HELEN, and PacBio HiFi assembly.

| Sample | Sequencing Platform | Method | | Percentage errors | | | |
|---|---|---|---|---|---|---|---|
| | | Assembler | Polisher | Balanced | Identity | Deletion | Insertion |
| CHM13 Chr-X | PacBio HiFi | Canu | Racon | 0.026% | 0.007% | 0.013% | 0.006% |
| | Nanopore | Shasta | MarginPolish & HELEN | 0.081% | 0.010% | 0.043% | 0.027% |

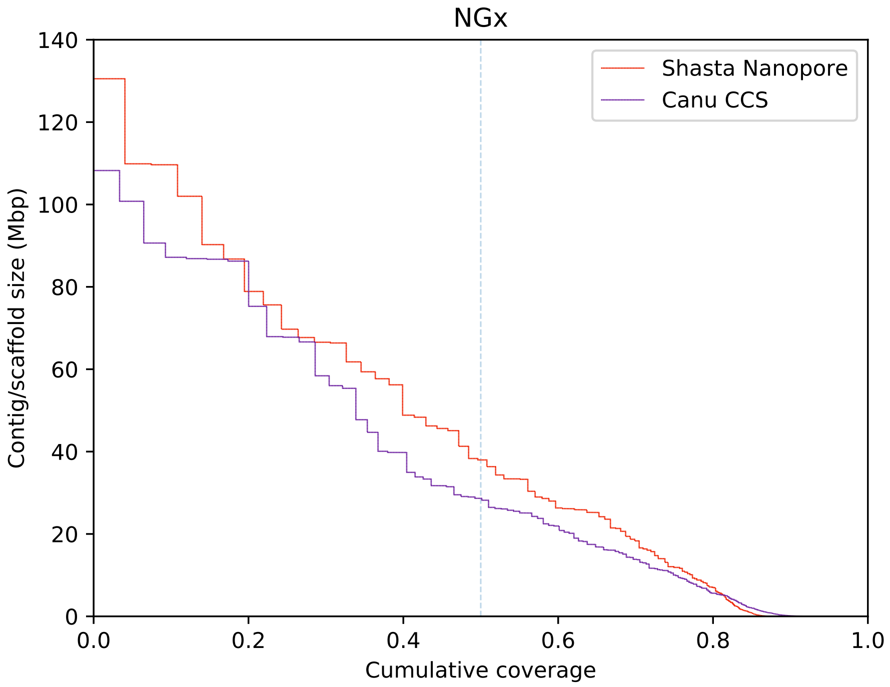


Supplementary Figure 5: Contig NGx for CHM13 Shasta-HELEN nanopore assembly vs Canu CCS (HiFi) assembly

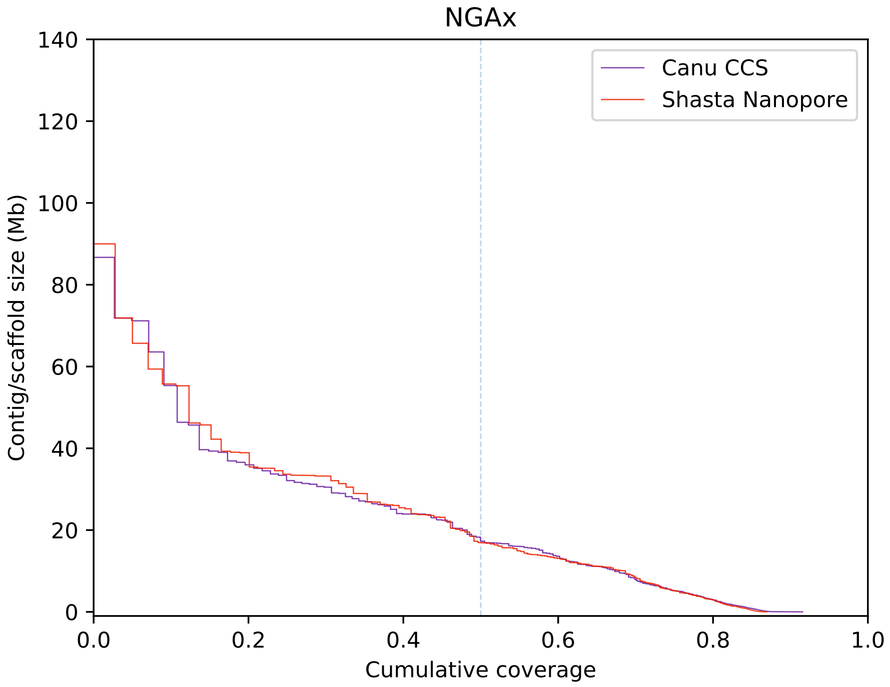


Supplementary Figure 6: Contig NGAx for CHM13 Shasta-HELEN nanopore assembly vs Canu CCS (HiFi) assembly

**Assembling, polishing and scaffolding 11 human genomes at near chromosome scale**

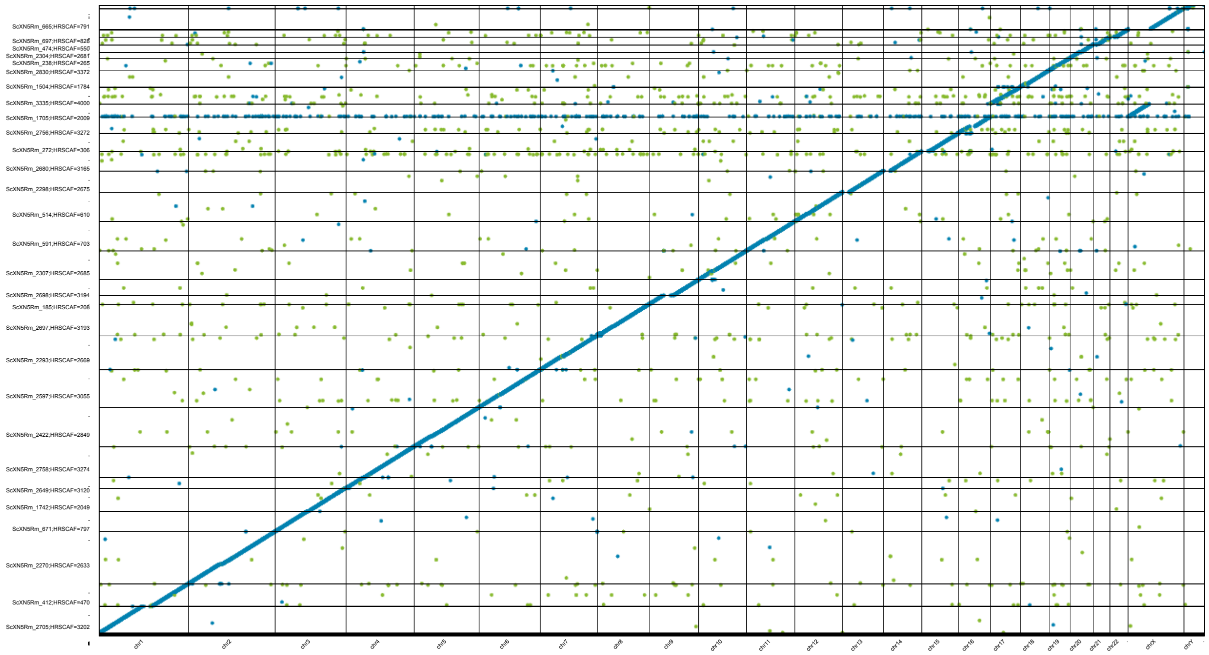

Supplementary Figure 7: Dotplot for the scaffolded HG002 assembly, aligned with GRCh38. Blue dots represent unique alignments and orange dots represent repetitive alignments.

Supplementary Table 26: QUAST results for all 11 Shasta assemblies scaffolded with HiRise, post polishing with MarginPolish-HELEN

| Sample | # contigs | Total length | N50 | NG50 | # mis-assemblies | # scaffold gap extensive mis-assembies | Genome fraction (%) | # mismatches per 100 kbp | # indels per 100 kbp | Total aligned length | NA50 | NGA50 |
|---|---|---|---|---|---|---|---|---|---|---|---|---|
| GM24143 | 1,184 | 2,802,523,049 | 129,960,437 | 128,216,303 | 1,466 | 4 | 95.027 | 128.28 | 142.79 | 2,792,775,664 | 20,657,530 | 16,966,477 |
| GM24149 | 1,323 | 2,816,683,224 | 129,643,816 | 128,275,807 | 1,530 | 11 | 95.417 | 130.24 | 134.58 | 2,804,735,382 | 18,446,390 | 15,435,923 |
| GM24385 | 1,019 | 2,819,527,260 | 118,169,209 | 102,591,941 | 1,335 | 6 | 95.606 | 127.19 | 152.25 | 2,809,570,528 | 22,369,161 | 16,601,924 |
| HG00733 | 1,056 | 2,800,455,909 | 129,857,865 | 118,785,172 | 1,337 | 8 | 94.974 | 126.16 | 138.09 | 2,791,610,554 | 22,141,375 | 17,570,210 |
| HG01109 | 1,156 | 2,821,098,626 | 130,282,751 | 130,166,418 | 1,529 | 5 | 95.559 | 136.73 | 140.63 | 2,809,413,640 | 19,932,703 | 17,228,023 |
| HG01243 | 1,006 | 2,819,162,443 | 128,571,344 | 118,762,399 | 1,381 | 7 | 95.517 | 137.47 | 143.03 | 2,808,041,766 | 22,146,722 | 17,559,055 |
| HG02055 | 977 | 2,819,933,140 | 130,184,428 | 128,180,737 | 1,387 | 8 | 95.587 | 141.91 | 162.46 | 2,809,195,864 | 21,057,279 | 18,446,049 |
| HG02080 | 934 | 2,803,570,658 | 129,931,575 | 128,451,196 | 1,470 | 9 | 95.041 | 127.98 | 134.36 | 2,793,854,132 | 20,418,609 | 16,379,851 |
| HG02723 | 982 | 2,805,356,030 | 130,365,062 | 128,975,828 | 1,499 | 9 | 95.06 | 143.45 | 147.13 | 2,794,747,200 | 20,232,566 | 17,865,825 |
| HG03098 | 926 | 2,811,385,538 | 130,040,472 | 128,535,908 | 1,439 | 4 | 95.391 | 144.36 | 170.40 | 2,801,774,564 | 22,165,948 | 17,439,948 |
| HG03492 | 901 | 2,811,782,250 | 130,277,907 | 100,251,163 | 1,381 | 7 | 95.362 | 126.54 | 147.23 | 2,803,106,787 | 20,001,587 | 16,836,756 |
